# Expanding the space of protein geometries by computational design of *de novo* fold families

**DOI:** 10.1101/2020.04.14.041772

**Authors:** Xingjie Pan, Michael Thompson, Yang Zhang, Lin Liu, James S. Fraser, Mark J. S. Kelly, Tanja Kortemme

## Abstract

Naturally occurring proteins use a limited set of fold topologies, but vary the precise geometries of structural elements to create distinct shapes optimal for function. Here we present a computational design method termed LUCS that mimics nature’s ability to create families of proteins with the same overall fold but precisely tunable geometries. Through near-exhaustive sampling of loop-helix-loop elements, LUCS generates highly diverse geometries encompassing those found in nature but also surpassing known structure space. Biophysical characterization shows that 17 (38%) out of 45 tested LUCS designs were well folded, including 16 with designed non-native geometries. Four experimentally solved structures closely match the designs. LUCS greatly expands the designable structure space and provides a new paradigm for designing proteins with tunable geometries customizable for novel functions.

**One Sentence Summary:** A computational method to systematically sample loop-helix-loop geometries expands the structure space of designer proteins.

## Main text

Design of proteins with new and useful architectures and functions requires precise control over molecular geometries ^1,2^. In nature, proteins adopt a limited set of protein fold topologies ^3-5^ that are reused and adapted for different functions. Here we define “topology” as the identity and connectivity of secondary structure elements (**Fig. 1A**). Within a given topology, geometric features including length and orientations of secondary structure elements are often highly variable ^3,4^. These considerable geometric differences between proteins with the same topology are necessary as they define the exquisite shape and physicochemical complementarity characteristic of protein functional sites. Creating proteins with new functions *de novo* therefore requires the ability to design proteins not only with different topologies, but also distinct custom-shaped geometries within these topologies optimal for each function (**Fig. 1A**).

**Figure 1.**
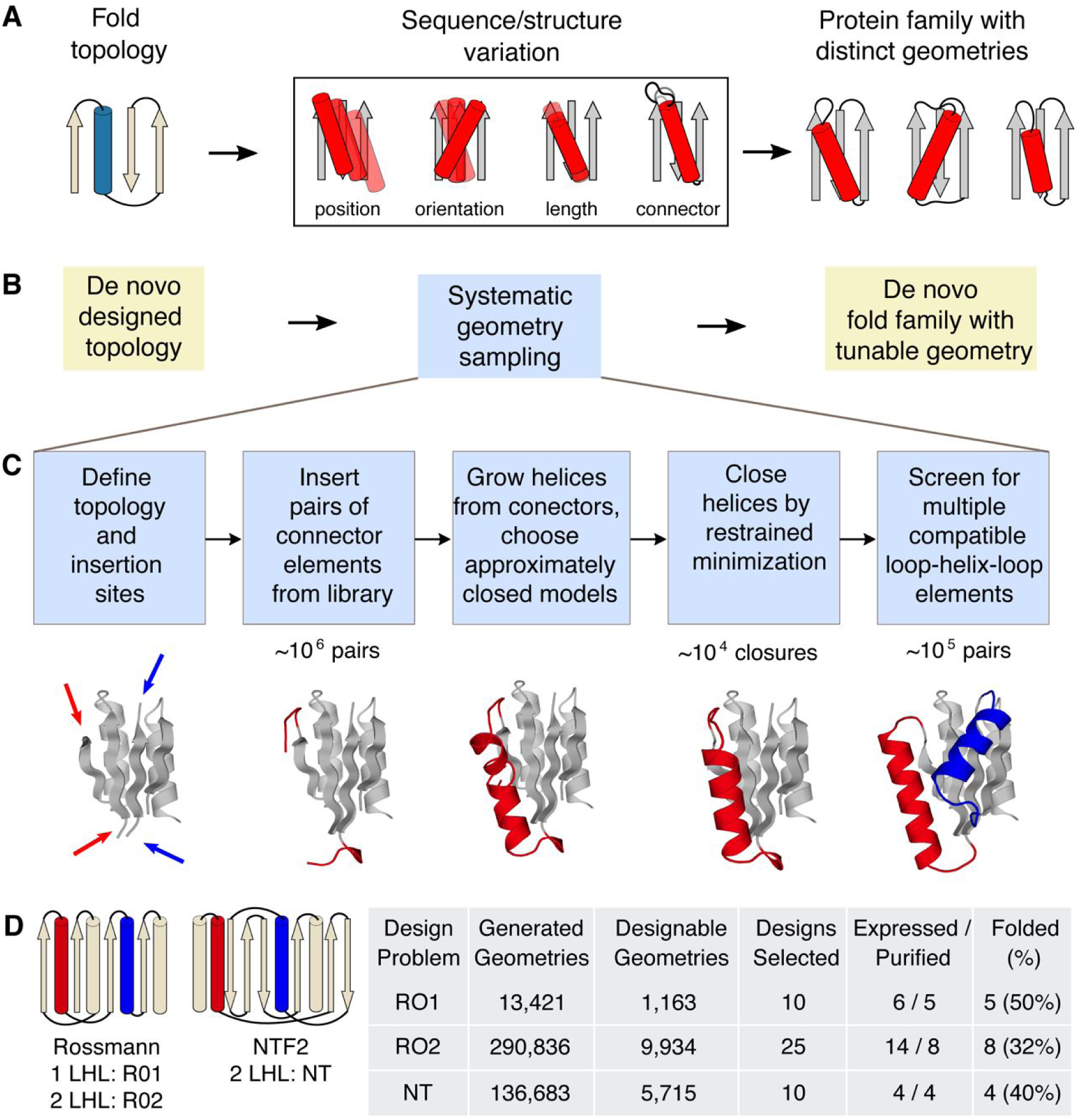
LUCS sampling strategy to create *de novo* designed protein fold families with tunable geometries. **A.** In nature, protein fold topologies (left) are diversified to create families of proteins with distinct geometries (right) optimized for function. Alpha-helices are shown as cylinders and beta-strands as arrows. The box shows schematic representations of common types of geometric variation. **B.** The LUCS computational design protocol seeks to mimic the ability of evolution to diversity protein geometries to generate *de novo* designed fold families. **C.** Schematic of the LUCS protocol for sampling LHL geometries. The reshaped LHL units are colored in red and blue. Typical numbers of models generated at major stages of the protocol are indicated. **D.** Designed fold families. Schematic shows fold topologies and design problems (Rossman fold with 1 or 2 reshaped LHL units, and NTF2 fold with 2 reshaped LHL units). Also shown are numbers for geometries generated by LUCS, designed models that passed quality filters, and experimentally characterized designs for three design problems. % folded indicates the fraction of experimentally tested designs that adopted folded structures.

Computational design has been successful in mimicking the ability of evolution to generate diverse protein structures spanning helical ^6-10^, alpha-beta ^11-13^ and beta-sheet ^14,15^ fold topologies, including novel folds ^16^. However, most design methods do not include explicit mechanisms to vary geometric features within a topology. For instance, successful design methods assemble protein structures from peptide fragments using a definition of the desired fold and topological rules derived from naturally occurring structures ^12^. Subsequent iterative cycles of fixed-backbone sequence optimization and fixed-sequence structure minimization ^16^ refine atomic packing interactions, but do not create substantial changes in geometry. An exception are methods that use parametric equations to sample backbone variation ^17^ or take advantage of modular protein elements, but these methods are restricted to helical bundles ^6,8,10^ or repeat protein ^18^ architectures, respectively.

Here we sought to develop a generalizable computational design approach that mimics the ability of evolution to create considerable geometric variation within a given fold topology (**Fig. 1**). When analyzing geometric variation in existing protein fold families, we found that 84% of naturally occurring fold families contain variations in loop-helix-loop (LHL) elements (**Supplementary Figure S1**). We hence reasoned that a method that systematically samples geometric variation in these units would not only be able to recapitulate a large fraction of geometric diversity in naturally occurring structures but also to create fold families of *de novo* designed proteins with tunable geometries (**Fig. 1B**).

To develop a generalizable method that systematically samples geometries of LHL, we first examined the connecting loop elements in native LHL units. For all LHL loop elements from all CATH superfamilies ^3^ of non-redundant structures, 72.8% contained ≤ 5 residues (**Supplementary Figure S2A**). We therefore focused on sampling LHL units with loop elements that have 2, 3, 4 and 5 residues. We extracted 313,072 loops connecting to helices from the Rosetta non-redundant fragment database ^19^ and sorted loops into 12 libraries based on loop length and type of adjacent secondary structure (**Supplementary Table S1**). For each library, only non-redundant loops were kept (**Supplementary Methods**); this procedure yielded between 224 and 5,826 loops per library. The loop libraries had degeneracies (total number of loops divided by the number of non-redundant loops in each library) ranging from 4.4 to 202 (**Supplementary Figure S2B**), indicating that evolution frequently used similar loop structures in different proteins. We therefore reasoned that the identified loop element libraries could also be used to computationally sample novel protein structures that have not been explored by nature.

We developed a protocol called loop-helix-loop unit combinatorial sampling (LUCS, **Fig. 1C, Supplementary Figure S3**). LUCS starts with an input protein fold, which can be naturally occurring or as in our case *de novo* designed, and a definition of gaps to insert LHL units. The first step systematically samples all loop element pairs in our libraries (**Supplementary Table S1**). For each gap, all pairs of loops from the libraries are inserted and any loops that clash with the input structure are removed. The second step tests all remaining pairs of loops for supporting LHL units by growing helices from each loop. If helices grown from the two ends meet in the middle, excess residues are removed in the third step and the gap closed by energy minimization with a chain-break penalty and hydrogen bond restraints. Closed LHL units with distorted hydrogen bonds geometries, steric clashes or suboptimal interactions between designed backbones and the environment are discarded (**Supplementary Methods**). In a fourth step, combinations of LHL units at different positions can be screened to yield final structures that have multiple compatible LHL units with systematically sampled lengths and orientations.

To validate the ability of LUCS to generate distinct geometries within given fold topologies, we applied the method to three design problems (**Fig. 1D**). In the first two design problems, we varied one (RO1) or two (RO2) LHL units of a *de novo* designed protein ^12^ (PDB:2LV8) with a Rossmann fold topology. In the third problem, we varied two LHL units of a *de novo* designed protein ^20^ (PDB:5TPJ) with an NTF2 fold topology (NT). In principle, LUCS can sample topologies with arbitrary number of LHL units. For the systems we tested, systematic sampling of the geometries of each LHL unit generated approximately 10 ^4^ LHL elements for each gap. To limit the required computing power, we screened 10 ^6^ random combinations of LHL units and generated between 10 ^4^-10 ^5^ final backbone structures for each design problem (**Supplementary Table S2**). We then applied the Rosetta FastDesign protocol (**Supplementary Methods**) to optimize sequences for all residue positions within 10 Å from the new LHL elements. The number of designed residues for each backbone was between 33 and 87. We note that Rosetta FastDesign also introduces structural changes outside the reshaped LHL elements of the designed fold through gradient-based torsion minimization, although these changes are small (backbone heavy atom root-mean-square deviation (RMSD) < 1 Å). Following sequence design, we filtered the design models computationally using a set of quality criteria that included a minimal number of buried unsatisfied hydrogen bond donors/acceptors, tight atomic packing interactions in the protein core, and compatibility between sequences and local structures (**Supplementary Methods**).

For each of the three design problems, we selected 50 low Rosetta energy ^21^ designs from models that passed the quality filters and had diverse conformations for further computational characterization. The Rosetta design simulations optimized low-energy sequences given a desired structure. To determine the converse, whether the desired structure is also a low energy conformation given the sequence, we conducted *ab initio* protein structure prediction simulations in Rosetta ^22^. For the Rossman fold designs, we required the lowest-energy predicted structure to be within 1 Å Cα RMSD of the design model. For the NTF2 fold designs, we used a less strict criterion requiring a number of low-energy models to be close to the design model, to account for the more difficult problem of sampling native-like structures for proteins larger than 100 amino acids. 10, 25 and 10 designs that passed these tests were chosen for experimental characterization for each of the three design problems, respectively (**Fig. 1D, Data S1, S2**). The designed proteins were recombinantly expressed in *E. coli* and purified using His-tag affinity and size exclusion chromatography. For monomeric designs, we measured near-UV circular dichroism (CD) spectra, thermal melts monitored by CD, one dimensional 1H nuclear magnetic resonance (NMR) spectra and 2-dimensional ^15^N HSQC NMR spectra to assess formation of stable secondary and tertiary structure. 5/10, 8/25 and 4/10 designs were found to be well folded for each of the three design problems, respectively (**Fig. 1D, Supplementary Figure S4, Supplementary Table S3**).

To assess whether the designed structures adopted their intended geometries, we solved structures for three designs (RO2-1, RO2-20, and RO2-25) that sampled two LHL units in the Rossmann fold topology using nuclear magnetic resonance spectroscopy (NMR), and one structure for the NTF2 fold topology designs (NT-9) by X-ray crystallography (**Supplementary Methods, Supplementary Figure S5, Supplementary Tables S4-5**). The experimentally solved Rossmann fold structures closely matched the designed models (**Fig. 2 A-C**), with backbone heavy atom RMSDs between models and solved structures within 1.3 Å, and core hydrophobic side chains in good agreements with the designed models (**Supplementary Figure S6**). Among the loops of the designed LHL units, 5 loops were well converged (pairwise backbone RMSD within the ensemble of NMR models within 1 Å). The backbone heavy atom RMSDs between the converged loops of lowest energy NMR models and designs were within 1.6 Å (**Supplementary Figure S7**). In the crystallographic electron density map obtained at 1.5 Å resolution for the NTF2 fold design (NT-9), strong signal was clearly identifiable inside a surface pocket (**Fig. 2D**), which was interpreted as a bound phospholipid (1,2-diacyl-sn-glycero-3-phosphoethanolamine, see **Supplementary Methods**). The two N- and C-terminal helices (residues 1-20 and 113-128), which had not been reshaped by LUCS, were pushed apart to accommodate the ligand, leading to an overall backbone heavy atom RMSD between design and model of 2.7 Å. However, when excluding the N- and C-termini helices and aligning the remainder of the design, the backbone heavy atom RMSD between the model and the solved structure was 1.4 Å (**Fig. 2E**). Moreover, the designed side chain packing interactions between the reshaped helices were in excellent agreement with the design (**Fig. 2F**). Taken together, our structural analysis confirmed the designed geometry in the reshaped regions for all 4 designs. The presence of a ligand in the NT-9 design is consistent with the known ability of the NTF2 fold to bind to diverse hydrophobic small molecules, and highlights the exciting possibility to introduce new functions such as ligand binding by reshaping protein geometries.

**Figure 2.**
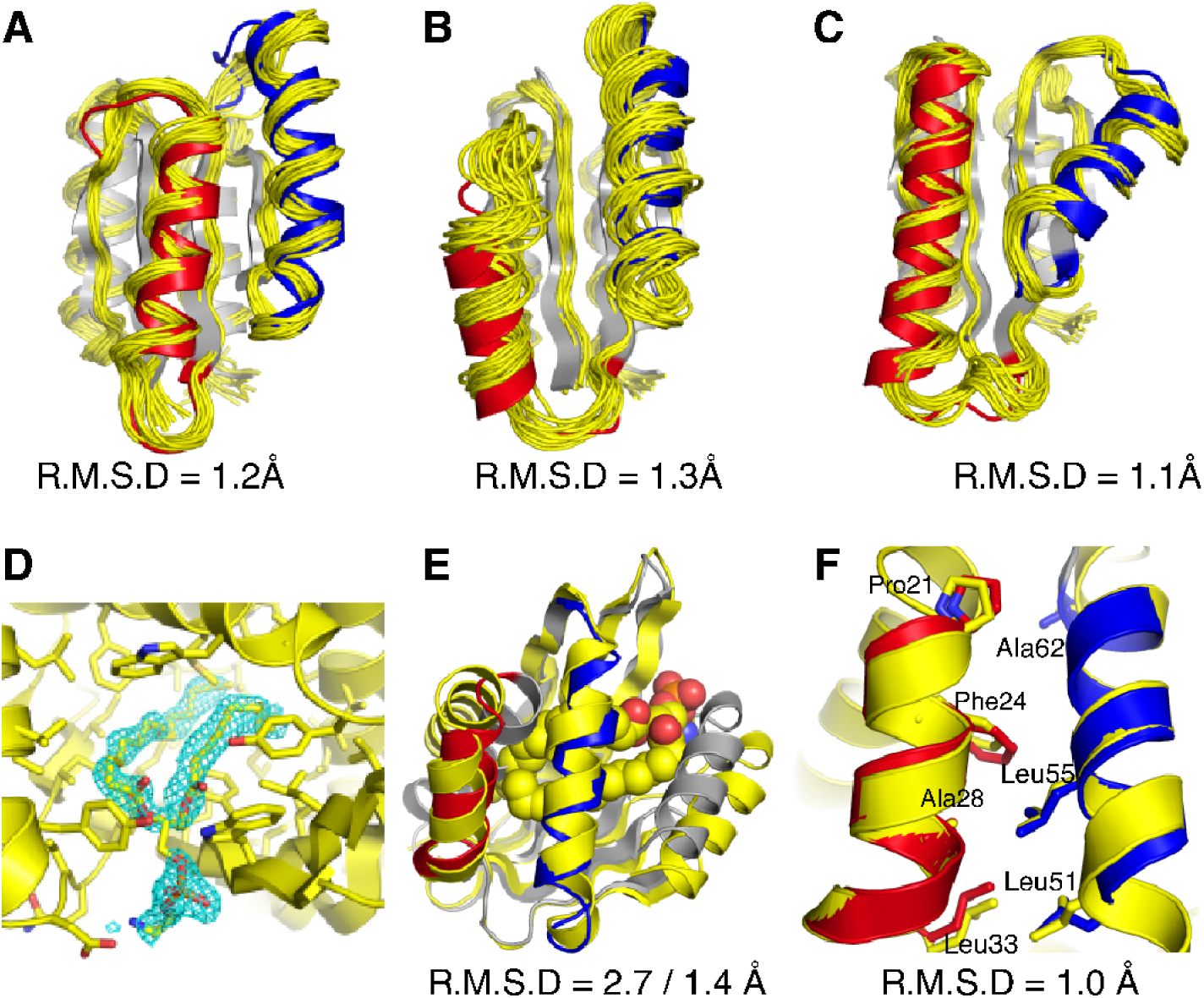
Close agreement between models and experimentally determined structures of designed proteins. **A-C**, designs for the Rossmann fold topology and **D-F**, designs for the NTF2 fold topology. Experimentally determined structures are shown in yellow and design models in grey with the reshaped LHL elements highlighted in red and blue. **A-C.** Comparison between computational models and NMR structures for designs RO2_1(**A**), RO2_20(**B**) and RO2_25(**C**). Also shown are the backbone heavy atom RMSDs calculated using the lowest energy structure from the NMR ensemble. **D**. The binding pocket of a phosphatidylethanolamine ligand. The 2Fo − Fc electron density map (cyan) for the ligand molecule is shown at 1.0 σ level. **E**. Comparison between computational model and X-ray crystal structure for the design NT_9. The phosphatidylethanolamine ligand is shown in spacefill representation (carbon atoms in yellow, oxygen atoms in red, phosphorus atoms in orange, and nitrogen atoms in blue). Also shown are the backbone heavy atom RMSDs calculated including or excluding the terminal helices, respectively. **F.** Alignment between the designed helices in the computational model and the experimentally solved structure. The hydrophobic residues at the packing interface are shown in stick representation. The RMSD shown includes the helix backbone heavy atoms and side chain heavy atoms displayed as sticks.

We next analyzed the magnitude of the geometric differences between our designs. We first compared the backbone heavy atom RMSDs between the reshaped helices of all well folded designs (**Fig. 1D**) after aligning the non-reshaped regions using both the design models and experimentally solved structures (**Fig. 3A, Supplementary Figure S8**). For the designs with one LHL unit reshaped, 18 out of 20 off-diagonal differences are more than 3Å (**Fig. 3A, left**). For the designs with two LHL units reshaped, 55 out of 68 off-diagonal differences are more than 4Å (**Fig. 3A, middle and right**). This scale of variation exceeds the backbone changes generated by existing flexible backbone design methods ^23,24^ that are typically smaller than 2Å RMSD. For each well-folded design, we also identified the closest existing structures in the protein data bank (PDB) using TM-align ^25^. Remarkably, 15 out of the 17 designed LHL units were significantly different (RMSD > 3Å for one LHL reshaped designs and RMSD > 4 Å for two LHL reshaped designs) from their closest match in the PDB (**Fig. 3A, Supplementary Figure S9**), indicating that the design protocol not only generates stable structures with considerable conformational divergence, but also geometries not observed in known structures.

**Figure 3.**
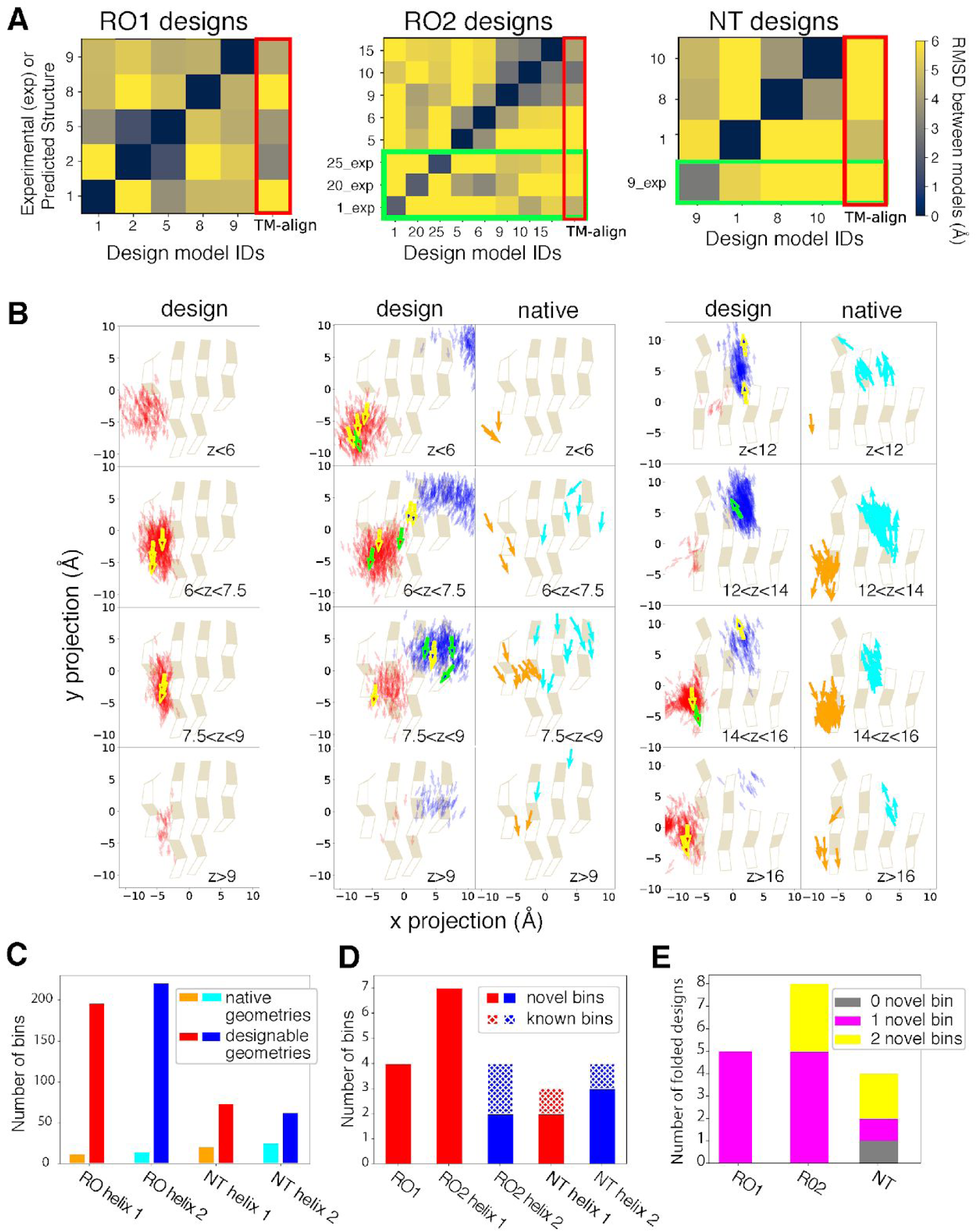
Geometry space sampled by d*e novo* designed fold families. In **A** and **B**, the columns show the 3 design problems: Left, Rossman fold with one designed LHL unit (RO1); middle, Rossmann fold with two designed LHL units (RO2); right: NTF2 fold with two designed LHL units (NT). **A.** Heatmaps showing backbone RMSDs between the reshaped LHL-regions of well-folded designs, comparing design models (x axis) with experimentally determined structures (_exp) or lowest-scoring models from Rosetta structure prediction (y axis). Green boxes show RMSDs calculated using experimentally solved structures. Red boxes (right columns) show the RMSDs between designs and the closest known structures found by TM-align. **B.** Projection of centers and directions of designed helices onto the underlying beta sheets. For the RO2 (middle) and NT (right) columns, left and right panels show distributions for designs and known structures, respectively. Sampled designable models (Fig. 1D) are represented by small arrows with reshaped helices colored in red and blue. The experimentally confirmed folded designs (Fig. 1D) are represented as bold arrows with yellow boundaries and the experimentally solved structures are represented as bold arrows with green boundaries. Helices are shown on 4 z-level planes based on their distances from the beta-sheet projection plane. For z-levels that have more than 1000 sampled structures, only 1000 randomly selected helices are shown. Projections for known Rossmann fold (middle) and NTF2 fold (right) protein structures are shown with the two helices corresponding to the designed regions colored in orange and cyan. The Rossmann fold structures are from the CATH superfamily 3.40.50.1980 and the NTF2 fold structures are from the CATH superfamily 3.10.450.50. **C.** Number of structure bins occupied by known structures (orange, cyan) and sampled by designable models generated by LUCS (red, blue). **D.** Structure bins occupied by well folded designs. **E.** Classification of the well folded structures by the number of novel structure bins they occupy.

We further analyzed the distribution of sampled geometries and their coverage of designable backbone structure space, where a structure is defined as designable if at least one sequence folds into that structure. As a computational approximation, we defined the models that passed the quality filters after the first iteration of sequence design (**Supplementary Methods**) as designable because they had good core packing, hydrogen bond satisfaction and local sequence structure compatibility with the designed sequence. We projected the center and directions of the helices onto the underlying beta sheets (**Fig. 3B**). The sampled helices from designable models at each position encompassed the distributions derived from native protein structures in the PDB (**Fig. 3B, right panels**). For the NTF2 fold, the distributions sampled in the designs were slightly shifted to the upper left when compared to the distributions in known structures (**Supplementary Figure S8**). This difference could be a result of the presence of a C-terminal helix in our designs occupying the region shown in the right of the space projection, whereas C terminal helices were often missing in the ensemble of known structures. Overall, since the number of known protein structures for a given topology is limited, the structure space covered by the known structures is much sparser than the space covered by the sampled structures. We quantified the size of structure space by dividing the 6-dimensional space of helix centers and orientations into bins (**Supplementary Methods**). For the geometries sampled in this work, the known structures covered between 12 and 26 bins, while LUCS generated structures covered between 63 and 221 bins (**Fig 3C**). The 17 well folded designs (**Fig. 1D**) sampled between 3 and 7 bins for each helix, respectively, and the majority (18/22) of these bins were not covered by known structures (**Fig 3D**). All but one of the well folded designs had at least one helix in a novel bin. Five well folded designs had both helices in novel bins (**Fig 3E**). Taken together, these results show that LUCS generates highly diverse geometries encompassing those found in nature but also exceeding known structure space.

We next sought to understand in more detail how the unique backbone geometries of the designed proteins were defined by the precise details of their non-covalent intramolecular interactions. The three experimentally solved Rossmann fold topology structures had distinct sequence patterns (**Fig. 4A**) resulting in distinct packing arrangements (**Fig. 4B, C**) in their hydrophobic cores. The beta sheets favored beta branched residues as expected, but the side chain sizes varied across different designs and resulted in differential hydrophobic packing. In particular, we observed previously described knob-socket type packing motifs ^26^ (**Fig. 4C, Supplementary Figure S10**) where nonpolar side chains fit into pockets formed by three residues on helices. These arrangements result in matched geometries between the side chains from sheets and helices that likely contribute to specifying the three-dimensional arrangement of the helices. We also applied tertiary motif analysis using MASTER ^27^. For all well-folded designs, we were able to match tertiary motifs to both the designed loops and interacting secondary structure elements (**Supplementary Figure S11**). Moreover, we identified side chains mediating helix-helix, helix-sheet and helix-loop interactions that are similar in our designs and the corresponding matched tertiary motifs (**Fig. 4D**). Despite the close match between the local structures in the design and the tertiary motifs, the source proteins of the motifs had overall structures very different from the designs (**Supplementary Figure S11**). Since no tertiary motif information was used in backbone generation or sidechain design, we conclude that our design protocol, which is guided solely by the LUCS sampling protocol and the Rosetta energy function ^21^, recapitulated tertiary structure motifs that were used recurrently by nature.

**Figure 4.**
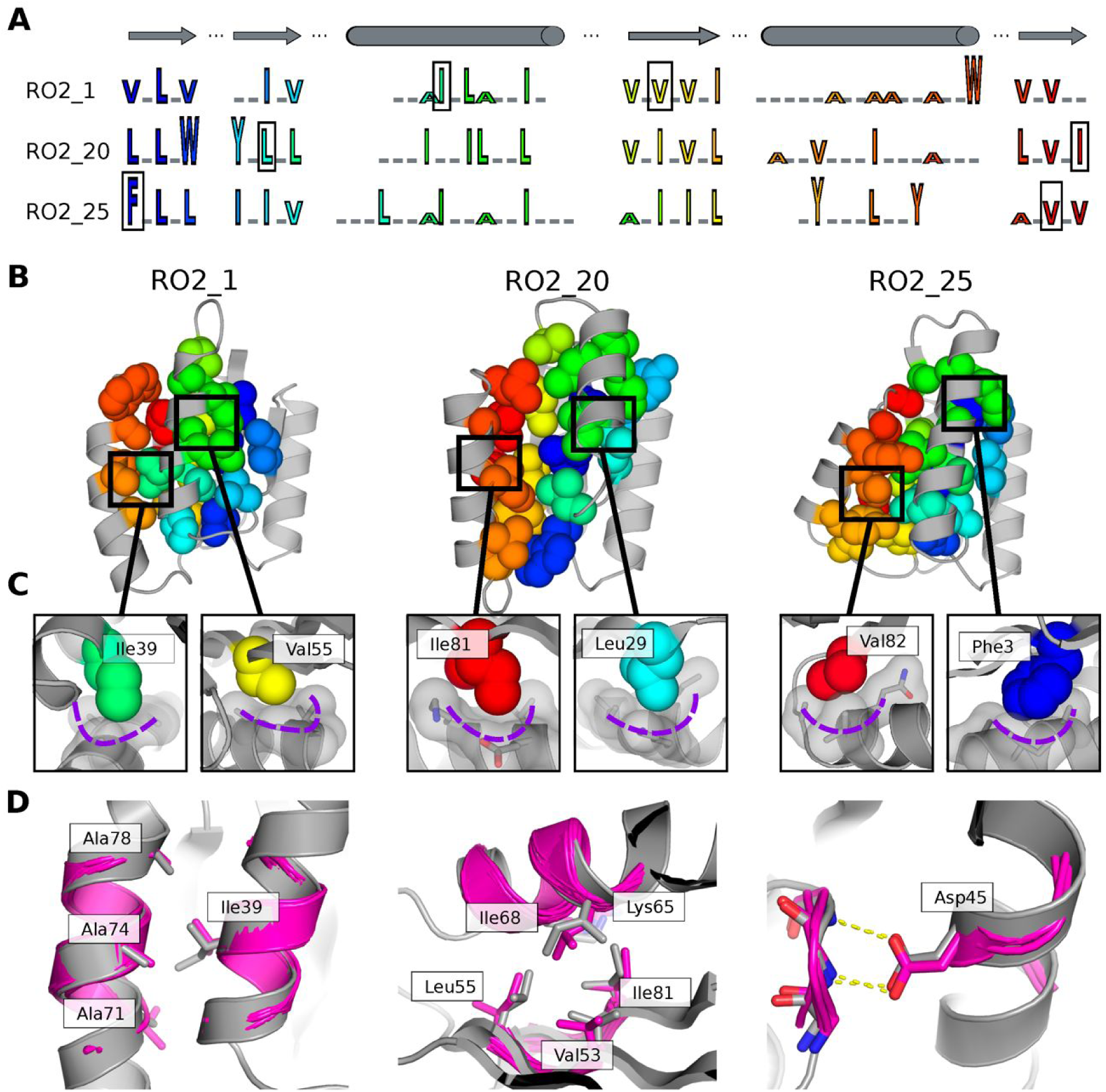
Structural features encoding distinct protein geometries. **A.** Sequence patterns of the hydrophobic cores in three designed models for the Rossman fold, aligned by corresponding secondary structure elements (top). Hydrophobic residues are shown as letters in rainbow colors ordered by position in the primary protein sequence and scaled by side chain size. Grey underlines indicate positions of surface exposed polar residues. The residues in the boxes are the knob residues shown in (**C). B.** Atomic packing of hydrophobic cores in the three experimentally determined structures for the Rossman fold (**Fig. 2**). The hydrophobic side chains in the designed cores are shown as spheres. **C.** Knob-socket packing motifs found in the designs. Three residues on a helix (grey sticks and surfaces) form a socket accommodating a knob residue shown as colored spheres. **D.** Examples of tertiary motifs matching the designed LHL structures. The designed structures are shown in grey and the matched motifs are shown in magenta. Sidechains of the best matched tertiary motifs and design models are shown as sticks.

Despite the more than 150,000 structures in the PDB, it is unknown how much backbone structure space is designable, and how much designable space is already covered by known structures. One way to probe the answers to these questions is by designing novel proteins that systematically explore the backbone space beyond known structures. Here we show that a large number of novel protein geometries can be sampled computationally. The experimentally validated, well-folded designs have geometries distinct from known structures. These results indicate that a large part of designable protein structure remains unexplored.

Previous key achievements in *de novo* design ^11-15,20^ focused on designing one or a few structures for diverse non-helical-bundle topologies by deriving design rules for specific topologies to identify the most favorable geometries. Proteins designed by this topology-centric strategy have pre-defined secondary structure sizes and loop torsions that are ideal to their topologies. In contrast, natural and LUCS generated structure families adopt non-ideal geometric features such as diverse helix positions, orientations, lengths and conformations of connector elements.

Exploring these non-ideal regions presents extra challenges ^28^. The topology-centric strategy typically finds deep energy minima and thereby succeeds in overcoming errors in energy functions. Sampling non-ideal geometric features can result in a smaller energy gap between the desired folded state and alternative states. Nevertheless, we show here that LUCS achieves a remarkably accurate atom-level control over diverse geometries. This success can at least partially be explained by the ability of LUCS to recover stable three-dimensional packing arrangements that are recurrent in nature (**Fig. 4D, Supplementary Figure S11**), but without using this information as input. Moreover, LUCS does not require prior definition of structural variation based on design rules identified in native structures ^20,29^ to generate diverse geometries that sample both known and new structural space. New protocols could exploit this ability to flexibly tune protein geometries during design simulations while simultaneously building new functional sites. The generalizable strategy underlying LUCS (**Fig. 1C**) could also be used for developing methods that sample other types of protein backbone geometries such as beta sheets.

We envision many applications for LUCS to precisely tune protein geometries for new protein functions that require atom-level control. By sampling LHL units, geometries of protein functional sites can be reshaped for ligand binding or protein-protein recognition. The systematic sampling of protein geometries should also enable designing dynamic proteins ^30^ that can switch between multiple distinct *de novo* designed conformations. Methods such as LUCS bring control over designable protein geometry space for arbitrary functions within reach.

## Acknowledgments

We would like to thank Muziyue Wu, Nicholas Hoppe, and members of the Kortemme lab for discussion.

## Author contributions

XP conceived the idea for the project. XP and TK conceived the computational and experimental approach. XP developed and performed the computational design. XP and YZ performed the majority of the experimental characterization. XP and MJSK determined the NMR structures. XP, MT, LL and JSF determined the crystal structure. JSF, MJSK and TK provided guidance, mentorship and resources. XP and TK wrote the manuscript with contributions from the other authors.

## Data and materials availability

Coordinates and structure files have been deposited to the Protein Data Bank (PDB) with accession codes 6VG7, 6VGA, 6VGB and 6W90. All other relevant data are available in the main text or the supplementary materials. Rosetta source code is available from rosettacommons.org. Upon publication, constructs will be made available via Addgene.

## Supplementary Materials

Materials and Methods

Fig S1 – S12

Table S1 – S5

Data S1 – S2

## References and Notes

1. Baker, D. An exciting but challenging road ahead for computational enzyme design. Protein Sci 19, 1817–9 (2010).

2. Kundert, K. & Kortemme, T. Computational design of structured loops for new protein functions. Biol Chem 400, 275–288 (2019).

3. Dawson, N.L., Lewis, T.E., Das, S., Lees, J.G., Lee, D., Ashford, P., Orengo, C.A. & Sillitoe, I. CATH: an expanded resource to predict protein function through structure and sequence. Nucleic Acids Res 45, D289–D295 (2017).

4. Fox, N.K., Brenner, S.E. & Chandonia, J.M. SCOPe: Structural Classification of Proteins--extended, integrating SCOP and ASTRAL data and classification of new structures. Nucleic Acids Res 42, D304–9 (2014).

5. Hou, J., Jun, S.R., Zhang, C. & Kim, S.H. Global mapping of the protein structure space and application in structure-based inference of protein function. Proc Natl Acad Sci U S A 102, 3651–6 (2005).

6. Huang, P.S., Oberdorfer, G., Xu, C., Pei, X.Y., Nannenga, B.L., Rogers, J.M., DiMaio, F., Gonen, T., Luisi, B. & Baker, D. High thermodynamic stability of parametrically designed helical bundles. Science 346, 481–485 (2014).

7. Jacobs, T.M., Williams, B., Williams, T., Xu, X., Eletsky, A., Federizon, J.F., Szyperski, T. & Kuhlman, B. Design of structurally distinct proteins using strategies inspired by evolution. Science 352, 687–90 (2016).

8. Thomson, A.R., Wood, C.W., Burton, A.J., Bartlett, G.J., Sessions, R.B., Brady, R.L. & Woolfson, D.N. Computational design of water-soluble alpha-helical barrels. Science 346, 485–8 (2014).

9. Hill, R.B., Raleigh, D.P., Lombardi, A. & DeGrado, W.F. De novo design of helical bundles as models for understanding protein folding and function. Acc Chem Res 33, 745–54 (2000).

10. Harbury, P.B., Plecs, J.J., Tidor, B., Alber, T. & Kim, P.S. High-resolution protein design with backbone freedom. Science 282, 1462–7 (1998).

11. Rocklin, G.J., Chidyausiku, T.M., Goreshnik, I., Ford, A., Houliston, S., Lemak, A., Carter, L., Ravichandran, R., Mulligan, V.K., Chevalier, A., Arrowsmith, C.H. & Baker, D. Global analysis of protein folding using massively parallel design, synthesis, and testing. Science 357, 168–175 (2017).

12. Koga, N., Tatsumi-Koga, R., Liu, G., Xiao, R., Acton, T.B., Montelione, G.T. & Baker, D. Principles for designing ideal protein structures. Nature 491, 222–7 (2012).

13. Huang, P.S., Feldmeier, K., Parmeggiani, F., Velasco, D.A.F., Hocker, B. & Baker, D. De novo design of a four-fold symmetric TIM-barrel protein with atomic-level accuracy. Nat Chem Biol 12, 29–34 (2016).

14. Dou, J., Vorobieva, A.A., Sheffler, W., Doyle, L.A., Park, H., Bick, M.J., Mao, B., Foight, G.W., Lee, M.Y., Gagnon, L.A., Carter, L., Sankaran, B., Ovchinnikov, S., Marcos, E., Huang, P.S., Vaughan, J.C., Stoddard, B.L. & Baker, D. De novo design of a fluorescence-activating beta-barrel. Nature 561, 485–491 (2018).

15. Marcos, E., Chidyausiku, T.M., McShan, A.C., Evangelidis, T., Nerli, S., Carter, L., Nivon, L.G., Davis, A., Oberdorfer, G., Tripsianes, K., Sgourakis, N.G. & Baker, D. De novo design of a non-local beta-sheet protein with high stability and accuracy. Nat Struct Mol Biol 25, 1028–1034 (2018).

16. Kuhlman, B., Dantas, G., Ireton, G.C., Varani, G., Stoddard, B.L. & Baker, D. Design of a novel globular protein fold with atomic-level accuracy. Science 302, 1364–8 (2003).

17. Crick, F. The packing of [alpha]-helices: simple coiled-coils. Acta Crystallographica 6, 689–697 (1953).

18. Brunette, T.J., Parmeggiani, F., Huang, P.S., Bhabha, G., Ekiert, D.C., Tsutakawa, S.E., Hura, G.L., Tainer, J.A. & Baker, D. Exploring the repeat protein universe through computational protein design. Nature 528, 580–4 (2015).

19. Gront, D., Kulp, D.W., Vernon, R.M., Strauss, C.E. & Baker, D. Generalized fragment picking in Rosetta: design, protocols and applications. PLoS One 6, e23294 (2011).

20. Marcos, E., Basanta, B., Chidyausiku, T.M., Tang, Y., Oberdorfer, G., Liu, G., Swapna, G.V., Guan, R., Silva, D.A., Dou, J., Pereira, J.H., Xiao, R., Sankaran, B., Zwart, P.H., Montelione, G.T. & Baker, D. Principles for designing proteins with cavities formed by curved beta sheets. Science 355, 201–206 (2017).

21. Park, H., Bradley, P., Greisen, P., Jr., Liu, Y., Mulligan, V.K., Kim, D.E., Baker, D. & DiMaio, F. Simultaneous Optimization of Biomolecular Energy Functions on Features from Small Molecules and Macromolecules. J Chem Theory Comput 12, 6201–6212 (2016).

22. Bradley, P., Misura, K.M. & Baker, D. Toward high-resolution de novo structure prediction for small proteins. Science 309, 1868–71 (2005).

23. Davey, J.A. & Chica, R.A. Multistate Computational Protein Design with Backbone Ensembles. Methods Mol Biol 1529, 161–179 (2017).

24. Ollikainen, N., Smith, C.A., Fraser, J.S. & Kortemme, T. Flexible backbone sampling methods to model and design protein alternative conformations. Methods Enzymol 523, 61–85 (2013).

25. Zhang, Y. & Skolnick, J. TM-align: a protein structure alignment algorithm based on the TM-score. Nucleic Acids Res 33, 2302–9 (2005).

26. Joo, H., Chavan, A.G., Phan, J., Day, R. & Tsai, J. An amino acid packing code for alpha-helical structure and protein design. J Mol Biol 419, 234–54 (2012).

27. Zhou, J. & Grigoryan, G. Rapid search for tertiary fragments reveals protein sequence-structure relationships. Protein Sci 24, 508–24 (2015).

28. Baker, D. What has de novo protein design taught us about protein folding and biophysics? Protein Sci 28, 678–683 (2019).

29. Lin, Y.R., Koga, N., Tatsumi-Koga, R., Liu, G., Clouser, A.F., Montelione, G.T. & Baker, D. Control over overall shape and size in de novo designed proteins. Proc Natl Acad Sci U S A 112, E5478–85 (2015).

30. Davey, J.A., Damry, A.M., Goto, N.K. & Chica, R.A. Rational design of proteins that exchange on functional timescales. Nat Chem Biol 13, 1280–1285 (2017).

