## Supplementary materials for "Expanding the space of protein geometries by computational design of *de novo* fold families"

**This PDF file includes:**

Materials and Methods
Figs. S1 to S12
Tables S1 to S5
Data S1 to S2

### Materials and Methods

Analysis of native loop-helix-loop (LHL) units in naturally occurring structures

Protein domain structures of all 2737 CATH superfamilies were downloaded from the CATH database (V4.1.0)[1]. Secondary structures for each protein were assigned using the DSSP algorithm[2] integrated in Rosetta[3] and used to identify LHL units in each structure. The number of LHL units in each superfamily was defined as the median number of LHL units in all structures from that superfamily. More than 83% of superfamilies had at least one LHL unit (**Figure S1A**). The LHL units in each superfamily were clustered. Two LHL units were clustered together if they connected the same types of secondary structure elements and the distances between the C$\alpha$ atoms of their starting and ending residues were within 3 Å. The helix RMSDs (see section: RMSD calculation between helices with different lengths) between all pairs of LHL units in the same clusters are shown in **Figure S1B**.

Generation of loop libraries (**Table S1**)

Each loop library contains loops with a defined length and connecting two secondary structure elements with defined types. Loop libraries were created by scanning the non-redundant protein structure database VALL[4]. All loops in VALL with a given length and connecting defined secondary structure types were selected. Loops were discarded if the helices they connected to had less than 6 residues or the strands they connected to had less than 3 residues. The number of loops in each library was recorded for subsequent redundancy calculation. If a loop had 2 Å or lower backbone heavy atom RMSD with other loops and the

RMSD of the C terminal residues was lower than 1.5 Å, the loop was considered redundant and removed. The redundancy of each loop library was defined as the number of all loops divided by the number of non-redundant loops.

Loop-helix-loop sampling

For each LHL unit to be sampled on a given scaffold structure, two insertion points were chosen as inputs. Residues between insertion points in the input structure were removed. Compatible loops with 2, 3, 4 or 5 residues were selected for each insertion point and inserted. All protein residues were mutated to alanine. Clashes between loop residues and the scaffold were detected. Two heavy atoms were defined to be clashing if the distance between atoms was smaller than the sum of their van der Waals radii times a scale factor of 0.6. Loops that did not clash with the scaffold were kept. All pairs of non-clashing loops were screened to test if a helix can be built to bridge the gap. For each pair of loops, 10-residue helices were grown from each of the ends of the loops using the ideal alpha helix dihedrals (phi=-57°, psi=-47°). The directions of half helices were calculated as the normalized average of all vectors pointing from atom N to atom C of each helix residue. If the dot product of directions of half helices was within 0.5, harmonic angle restraints were applied to the angle formed by the C$\alpha$ atoms of the helix start residue, front helix break residue and the helix end residue as well as the angle formed by the C$\alpha$ atoms of the helix start residue, back helix break residue and the helix end residue. Rosetta energy (only omega, rama_prepro and restraint terms were enabled) minimization was applied to minimize the restrained angles to align the two halves of the helix. During the minimization, phi, psi and omega torsions of the LHL unit residues were movable degrees of freedom. After aligning the helices, if there was a pair of residues on the half helices that had a backbone heavy atom RMSD within 3 Å, the excess residues were trimmed, and Rosetta energy minimization was applied to the LHL unit residue phi, psi and omega torsions to close the gap. During minimization, distances between the atom O and atom H in helix backbone hydrogen bonds as well as angles formed by atom N, atom H and atom O were restrained to maintain the helical structure. Quality filters were applied to the closed helix:

- Rosetta backbone hydrogen bond scores for all reshaped helix residues were lower than -0.7.
- There were no clashes (same definition as above) involving the reshaped helices after mutating all residues to valine (residues were mutated from ALA to VAL to check there was space between backbones for subsequent side chain design).
- The median contact degree (the number of C $\alpha$ atoms within 10 Å from the C $\alpha$ atom of a residue) of helix residues was greater than 1.
- The number of buried unsatisfied hydrogen bonds within the reshaped LHL units was less than 4.

If more than one pair of insertion points was sampled, two LHL units at different insertion points were deemed compatible if there was no clash (same definition as before) between them. A final ensemble of models was produced by applying a group of compatible LHL units at each insertion point. This step resulted in 13,421 models for RO1 designs, 290,836 models for RO2 designs, and 136,683 models for NT designs (**Table S2**). The protocol was developed using PyRosetta[5] and is available at:

https://github.com/Kortemme-Lab/loop_helix_loop_reshaping/releases/tag/1.0.0

The PyRosetta package used in this study was compiled from the Rosetta source code (commit: 3135d32229f5ebd35c8a716af00dcdfbbfa81805).

Sequence design

The spatial positions of residues were defined as the positions of their C$\beta$ atoms (C$\alpha$ for glycine). The sidechain directions were defined as the vector pointing from the C$\alpha$ atoms to the C$\beta$ atoms. A residue on the scaffold was defined as pointing toward the reshaped backbone region if there was a residue in the reshaped region such that the cosine of the angle between the vector from the surrounding residue C$\alpha$ atom to the reshaped residue C$\alpha$ atom and the sidechain direction vector of the surrounding residue was greater than 0.5. Residues within 10 Å (C$\alpha$-C$\alpha$ distance) from the backbone of the reshaped region and pointing toward the reshaped region were designed (i.e. allowed to change amino acid residue type and rotamer conformation). Residues within 8 Å from any designable residue and pointing toward the designable region were repackable (i.e. allowed to change rotamer conformation but keeping amino acid residue type). Residue types allowed for designable residues were determined by the extent of residue burial using the Rosetta LayerDesign task operation[6]. Cysteine and histidine were disallowed as designable residue types to avoid issues with disulfide bond formation and pH dependency. The Rosetta FastDesign [https://www.rosettacommons.org/docs/latest/scripting_documentation/RosettaScripts/Movers/movers_pages/FastDesignMover] and RotamerTrial [https://www.rosettacommons.org/docs/latest/scripting_documentation/RosettaScripts/Movers/movers_pages/RotamerTrialsMover] protocols were applied to design sequences that stabilize the backbone structures. The first iteration of design was done using the Dunbrack rotamer library without extra rotamers[7]. 58626, 432735 and 409101 sequences were designed for RO1, RO2 and NT. Designed structures that had less than 2 buried unsatisfied hydrogen bond donors or acceptors, had Rosetta hole scores[8] for the designed residues smaller than 0 and had fragment qualities[9] better than 2 Å were selected for the next iteration of designs. 1163, 9934 and 5715 designs passed the filters for RO1, RO2 and NT. For RO2 and NT, the selected designs were further optimized by 3 repeats of Rosetta FastDesign with extra rotamers enabled by the ex1 and ex2 options in the ExtraRotamersGeneric task operation. 49578 and 22831 sequences were designed for RO2 and NT. Final designs were filtered:

- Fragment qualities for the reshaped regions were better than 1 Å.
- Rosetta hole scores for the designable and repackable residues were lower than 0.
- Rosetta helix complementarity scores (Lawrence and Coleman shape complementarity)[10] for the reshaped helices were better than 0.6.
- There were no buried unsatisfied hydrogen bonds according to the custom buried unsatisfied hydrogen bond filter (next section).
- There were no oversaturated hydrogen bonds, i.e. hydrogen bond acceptors receiving hydrogen bonds from more than the allowed number of donors, for designable and repackable residues.
- The ratio of hydrophobic solvent accessible surface area (SASA) over the total SASA was lower than 58%.

722 and 98 designs passed the filters for RO2 and NT. The custom buried unsatisfied hydrogen bond filter sometimes underestimated the number of buried unsatisfied hydrogen bonds and the

holes filter allowed large hydrophobic holes which can exist in native proteins as binding sites. Therefore, designs that passed the filters with lowest residue average Rosetta scores were examined manually to check for possible issues such as hydrophobic voids and buried unsatisfied hydrogen bonds missed by the automatic filter. For the RO2 designs, we selected 50 top-scoring designs passing these manual criteria.

For the NT designs, we attempted to introduce cavities as potential binding pockets into the protein core. The 98 NT designs that passed the second iteration had hydrophobic cavities because the Rosetta layered design task operation assigned the residues surrounding the cavities to the core layer. We manually selected 10 low energy designs that passed iteration 2 (picking the lowest energy designs that were substantially different from each other) for further sequence design simulations to introduce pockets with polar residues into the proteins. We used the same sequence design protocol as in iteration 2, but with the restriction that 1-3 pocket residues had to be polar residues. For each of the 10 selected designs, 100 new sequences were designed. We then filtered the new designs with the same automatic filters followed by manual inspection as the previous round. 202 designs passed the automatic filters, of which 50 were selected by their Rosetta energy. For each of the 10 designs selected from iteration 2, at least one design derived from it was selected. The sequence design code is available at: https://github.com/Kortemme-Lab/local_protein_sequence_design/releases/tag/1.0.0

Custom buried unsatisfied hydrogen bond filter

The standard Rosetta buried unsatisfied hydrogen bond filter[11] overestimated the number of buried unsatisfied hydrogen bond donors and acceptors in many native structures (**Figure S12**) because it identified hydrogen bonds based on one static model. However, due to modeling errors and protein conformational flexibility, some of the identified buried unsatisfied hydrogen donors or acceptors present in the models may nevertheless form proper hydrogen bonds. In order to find only the buried unsatisfied hydrogen bond donors or acceptors that cannot be compensated by allowing some conformational flexibility, we developed a structure quality filter called the backrub ensemble consensus buried unsatisfied hydrogen bond (BECBUBH) filter. For each protein residue, except for the N- and C-terminal residues, the BECBUBH filter generated 5 structures by applying a local backrub[12] move on the residues and each of their two neighboring residues. The filter then determined the buried unsatisfied hydrogen bond donors or acceptors for each structure. Only buried unsatisfied donors or acceptors that were consistently present in all 5 structures were recorded. We used the BECBUBH filter to filter designed models.

*Ab initio* structure prediction

Rosetta *ab initio* structure prediction simulations were run for the top 50 selected designs for each design problem (**Table S2**). Fragments for structure predictions were generated using the make_fragments.pl script[4] distributed with Rosetta. The command for fragment generation was

fragments.pl -verbose -id design_id -frag_sizes 3,9 -n_frags 200 -n_candidates 1000 sequence.fasta

The structures of designs were predicted using the AbinitioRelax application in Rosetta. 20,000 models were generated for each designed sequence. The command was

AbinitioRelax.linuxgccrelease -abinitio:relax -use_filters true -abinitio::increase_cycles 10 -abinitio::rg_reweight 0.5 -abinitio::rsd_wt_helix 0.5 -abinitio::rsd_wt_loop 0.5 -relax::fast -in:file:fasta sequence.fasta -in:file:frag3 fragments_3mer_file -in:file:frag9 fragments_9mer_file -psipred_ss2 ss2_file_from_frag_generation -nstruct num_output -out:sf score_file_output -out:file:silent silent_file_output

For NT designs, standard Rosetta *ab initio* structure prediction simulations failed to sample models sufficiently close to the target model. The target model typically had a significantly lower Rosetta energy than any of the prediction models, indicating a sampling issue. To overcome this problem, we biased *ab initio* structure prediction simulations to favor low RMSD fragments by setting the weight of the score term FragmentCrmsd to be 5 and priority to be 800 in the fragment generation step. We then run standard *ab initio* structure prediction using the biased fragment set. We confirmed that designs folded into the desired target conformation within 1.5 Å when biasing the input fragments used during the structure prediction calculations.

Projecting helices to underlying beta sheets

Models with the Rossmann fold or NTF2 fold were aligned to PDB:2LV8 or PDB:5TPJ, respectively. The 3-dimensional (3D) helix centers were calculated by averaging the C$\alpha$ atom coordinates for all residues in a given helix. The 3D helix directions were defined as the average of C=O bond directions of the helix residues. The 3D centers and directions were projected onto a 2-dimensional (2D) plane by Cartesian projections. The beta sheet peptide bonds were shown as parallelograms where the C $\alpha$ atoms were located at the center of horizontal edges.

Binning the helix geometry space

Each helix was assigned a 6-dimensional vector whose first 3 coordinates were the helix center position and the last 3 coordinates were the direction of the helix. As above, the center of a helix was the average of the C $\alpha$ atom coordinates. The direction was the average of C=O bond directions normalized to a unit vector. The Cartesian space of helix centers was divided into 2Å cubes and the directions were divided into 8 octants. A 6-dimensional bin was assigned to a helix based on the cube and octant that the helix belonged to.

RMSD calculation between helices with different lengths

To calculate RMSD between two helices with different lengths, the longer helix was truncated. Only the middle part of the longer helix that had the same length as the shorter helix was kept. RMSDs between the corresponding backbone heavy atoms (N, C $\alpha$ and C) were calculated.

TM-align analysis

The designed models were submitted to the COFACTOR server[13], which used TM-align to find the 10 closest structures from the PDB. Except for the design RO2_5, the PDB structure 2KPO is the closest structure for all Rossmann fold designs and it ranked 2$^{nd}$ for the design

RO2_5. The PDB structure 5TPJ is the closest for all NTF2 fold designs. The difference between the designed geometries and known structures was then quantified by calculating the RMSD between the designed helices and the corresponding helices on the closest known structures with the same topologies found by TM-align (**Fig. 3A**).

Analysis of tertiary structure motifs

Tertiary structure motif analysis was performed using the MASTER[14] program. Small pieces of tertiary structures were specified manually from the well folded designs. The tertiary pieces included designed loops and fragments of interacting secondary structure elements involving the backbone reshaped regions (**Figure S11**). The helical element sizes were between 1 to 2 turns (5-9 residues) and the strand element sizes were between 3 to 5 residues. The tertiary pieces were extracted from the design models and saved as query pdb files. The query pdb files were converted to MASTER input query (pds) files by

```
createPDS --type query --pdb query.pdb
```

The query structures were searched against the standard MASTER database with

```
master --query query.pds --targetList MASTER/database/list --rmsdCut 0.5 --matchOut query.match --seqOut query.seq --bbRMSD --structOut query.struct
```

Protein expression tests

Plasmids (pET-28a(+)) encoding the designed proteins were ordered from Twist Bioscience. The DNA sequences of the designed proteins were inserted between the NdeI and XhoI restriction sites, which added the DNA coding sequence for an N-terminal MGSSHHHHHHSSGLVPRGSHM tag to the designed proteins. The plasmids were transformed into *Escherichia coli* BL21(DE3) cells. Proteins were expressed by overnight cell culture in 5mL autoinduction medium (ZY medium, 10 g/L tryptone, 5 g/L yeast extract) supplemented with the following stock mixtures: 20xNPS (1M $Na_2HPO_4$, 1 M $KH_2PO_4$, and 0.5 M $(NH_4)_2SO_4$), 50x 5052 (25% glycerol, 2.5% glucose, and 10% α-lactose monohydrate), 1000x trace metal mixture (50 mM $FeCl_3$, 20 mM $CaCl_2$, 10 mM each of $MnCl_2$ and $ZnSO_4$, and 2 mM each of $CoCl_2$, $CuCl_2$, $NiCl_2$, $Na_2MoO_4$, $Na_2SeO_3$, and $H_3BO_3$ in 60 mM HCl))[15] with 50 µg/ml kanamycin at 37°C. Cell cultures were aliquoted into 1mL aliquots. Cells were spun down by centrifuging at 20,000 g for 3 min. Cell pellets were resuspended in 1mL Phosphate-buffered saline (PBS) buffer (8g/L NaCl, 0.2g/L KCl, 1.44g/L $Na_2HPO_4$, 0.24g/L $KH_2PO_4$, pH=7.4) and lysed by sonication. Soluble and insoluble parts were separated by centrifuging at 21,000 g for 5 minutes. The solubility of a designed protein was assessed by Coomassie-stained SDS-PAGE (BIO-RAD Cat. #456-1095).

Protein expression and purification

Plasmids encoding the N-terminally His-tagged designed proteins were transformed into *Escherichia coli* BL21(DE3) cells. Colonies were inoculated into 5mL LB medium and cultured at 37°C for 12 hours. Seed cultures were inoculated into 1 L autoinduction medium and cultured at 225 RPM shaking speed at 37°C overnight. Cell cultures were centrifuged at 5,000 g for 5 minutes to spin down the cells. Cell pellets were resuspended in 30mL equilibration buffer (50mM Tris pH=7.5, 300mM NaCl, 10 mM imidazole) and lysed using a microfluidizer. Cell lysate was centrifuged at 39,000 g for 30 minutes to separate the soluble and insoluble fractions. The soluble fraction was mixed with 1mL Ni-resin beads (Thermo Scientific #88222) to pull down the His-tagged proteins. Ni-resin beads were washed 3 times with the wash buffer (50mM Tris pH=7.5, 300mM NaCl, 25mM imidazole). Proteins were eluted 3 times with 1mL elution buffer (50mM Tris pH=7.5, 300mM NaCl, 250mM imidazole). For crystallization, His-tags were cleaved by 10 NIH unit bovine thrombin (Sigma-Aldrich T4648-10KU) overnight at room temperature. Then the samples were further purified using a HiLoad® 16/600 Superdex® 75 pg size exclusion column (GE) with PBS buffer. The monomeric fraction was collected for subsequent characterization.

Analytical size exclusion chromatography

Protein samples purified by His-tag pull down (if sufficiently pure) or after additional purification with the HiLoad® 16/600 Superdex® 75 pg size exclusion column were analyzed using a Superdex® 75 10/300 GL size exclusion column from GE with PBS buffer. The relation between elution time and log molecular weight was fitted using a linear regression model with the BioRad Gel Filtration Standard (Catalog #151-1901).

Circular dichroism spectroscopy

Circular dichroism (CD) data were collected on a Jasco J-710 spectrometer. Purified RO1 proteins were diluted into 50mM phosphate buffer (pH 7.0). Purified RO2 and NT proteins were diluted into PBS buffer (8g/L NaCl, 0.2g/L KCl, 1.44g/L $Na_2HPO_4$, 0.24g/L $KH_2PO_4$, pH=7.4). The concentrations of diluted samples ranged from 2μM to 5μM. Protein concentrations were determined using the absorbance at 280 nm using a NanoDrop (Thermo Scientific). CD spectra were measured using a 1mm cuvette at 25°C. Melting curves at 220nm were measured by increasing temperature from 25°C to 95°C using a rate of 1°C/min.

1D-$^1$H NMR spectra

Purified proteins were exchanged into 50mM phosphate buffer (pH 7.0). Samples were concentrated to 15-200 μM. 56 mL $D_2O$ was added to 500 mL samples such that the final volume had 10% $D_2O$ and 90% $H_2O$. The 1D 1H spectra (pulse program: zgesgp) were measured at 297.9 K using a Bruker Avance I 800 MHz spectrometer with a Z-gradient TXI cryo-probe. The temperature was calibrated with 4% MeOD using the following coefficients of T (K) = (4.109 - D) * 0.008708 where D is the chemical shift difference between the $CH_3$ and OH protons in methanol. The spectra were processed with the program NMRPipe[16].

Protein expression for NMR structure determination

$^{15}N$ and $^{13}C$ labeled proteins were expressed by growing *E. coli* in M9 minimal medium that included $^{13}C$-glucose and $^{15}NH_4Cl$ (6g/L $Na_2HPO_4$, 3g/L $KH_2PO_4$, 0.5g/L NaCl, 0.5g/L $^{15}NH_4Cl$, 50mg/L EDTA, 8.3mg/L $FeCl_3$ x 6 $H_2O$, 0.84mg/L $ZnCl_2$, 0.13mg/L $CuCl_2$ x 2 $H_2O$, 0.1mg/L $CoCl_2$ x 6 $H_2O$, 0.1mg/L $H_3BO_3$, 0.016mg/L $MnCl_2$ x 6 $H_2O$, 0.2% (w/v) $^{13}C$-glucose, 1mM $MgSO_4$, 0.3mM $CaCl_2$, 1mg/L Biotin, 1mg/L Thiamine). Single bacterial colonies were first inoculated into 5mL seed culture and grown at 37 °C overnight. The seed culture was inoculated into 1L $^{15}N$ $^{13}C$ labeled M9 minimal medium and grown at 37 °C until $OD_{600}$ reached 0.5-0.7. Then 1mL 1M IPTG was added to induce protein expression at 18 °C overnight. The expressed proteins were purified following the Ni-resin pull down protocol described in the protein purification section. The His-tag purified proteins were further purified using the HiLoad® 16/600 Superdex® 75 pg size exclusion column from GE with 50mM phosphate buffer at pH 7.0. The monomeric fractions were collected and concentrated to 0.5-1 mM for NMR experiments.

Structure determination by NMR

Proteins labeled with $^{15}N$ and $^{13}C$ were exchanged into 50mM phosphate buffer (pH 7.0). 10% $D_2O$ was added to samples and the final protein concentrations were 0.74-0.8 mM. NMR spectra were measured at 297.9 K. Two dimensional (2D) $^{15}N$-HSQC (pulse program: fhsqcf3gpph), 2D $^{13}C$-HSQC (pulse program: hsqcetgpsisp2), 16 ms 3D HCCH-TOCSY (pulse program: hcchdigp3d) and 120 ms 3D simultaneous $^{13}C/^{15}N$-NOESY-HSQC (pulse program: noesyhsqcgpsismsp3d) spectra were measured using a Bruker Avance I 800 MHz spectrometer with a Z-gradient TXI cryo-probe. 3D CACB(CO)NH (pulse program: hncocacbgpwg3d) and 3D CACBNH (pulse program: hncacbgpwg3d) spectra were measured using a Bruker Avance DRX500 spectrometer with a Z-gradient QCI cryo-probe at 297.6 K (same temperature calibration as described above). NMR spectra were processed using the program NMRPipe[16] and indirect referencing to an external DSS standard was used[17]. Resonances were assigned using the program CCPN Analysis[18]. Backbone resonances were assigned using the 2D $^{15}N$-HSQC, 3D CACB(CO)NH and 3D CACBNH spectra. Sidechain resonances were assigned using the 2D

$^{13}$C-HSQC and 3D HCCH-TOCSY spectra. Distance restraints were generated using CCPN Analysis. Dihedral restraints were generated using the program DANGLE[19]. Hydrogen bond restraints included in the structure calculations were based on secondary structures predicted by DANGE and nOe patterns typical of alpha-helical secondary structure. The programs ARIA version 2.3.2[20] and CNS version 1.2 .1[21] were used to calculate the NMR structures. To solve the structures, 9 iterations of simulated annealing were performed using CNS. For the first 8 rounds of simulated annealing, the n_structures parameter was set to 100 and the n_best_structures parameter was set to 35. For the 9th round, the n_best_structures parameter was set to 20. Finally, a refinement in water was performed on the 20 structures from the 9th iteration. Otherwise, the default values were used for the remaining ARIA parameters. The ensemble of the refined structures was validated using the PDB validation server[22] and the Protein Structure Validation Suite (PSVS) server[23]. The agreement between structures and NMR data were assessed using the program PyRPF[24] integrated into the CCPN suite.

Protein crystallization

We concentrated the NT_9 protein to 25.2 mg/mL in buffer containing 20 mM Tris Buffer (pH=7.5) and 150 mM sodium chloride, and carried out initial crystallization trials using the JCSG I-IV commercial crystallization screen (Qiagen). Crystallization drops were prepared in 96-well sitting drop format by mixing 100 nl of protein solution with 100 nl of the mother liquor using a Mosquito liquid handling robot (TTP Labtech). Drops were sealed inside a reservoir containing an additional 100 µl of the mother liquor solution. Crystals were obtained from mother liquor containing 0.1M MES Buffer at pH=6, 30% PEG-600, 5% PEG-1000, and

10% Glycerol. Crystals from the initial screen were used for data collection without further optimization.

X-ray data collection and processing

Prior to X-ray data collection, crystals were flash-cooled by rapid plunging into liquid nitrogen. The high concentrations of polyethylene glycols and glycerol in the crystallization mother liquor allowed the crystals to be harvested and frozen directly without additional cryoprotection. We collected single-crystal X-ray diffraction data on beamline 8.3.1 at the Advanced Light Source. The beamline was equipped with a Pilatus3 S 6M detector (Dectris), the X-ray energy was set to 11111 keV, and the crystals were maintained at a cryogenic temperature (100 K) throughout the course of data collection.

We processed the X-ray data using the Xia2 system[25], which performed indexing, integration, and scaling with XDS and XSCALE[26], followed by merging with Pointless[27]. A resolution cutoff (1.50 Å) was taken where the completeness of the data fell to a value of approximately 90%. Although other metrics of data quality (such as CC1/2 and <I/σI>) suggest that a more aggressive resolution cutoff would be acceptable, we were limited by the data completeness that could be obtained with the minimum accessible sample-to-detector distance. Further information regarding data collection and processing is presented in Table S5. The reduced diffraction data were analyzed with phenix.xtriage (http://www.ccp4.ac.uk/newsletters/newsletter43/articles/PHZ_RWGK_PDA.pdf) to check for common crystal pathologies, none of which were identified.

Structure determination

We obtained initial phase information for calculation of electron density maps by molecular replacement using the program Phaser[28], as implemented in the PHENIX suite[29]. We identified a single copy of the protein in the asymmetric unit using the coordinates from our design model, consistent with an analysis of Matthews probabilities for the observed unit cell and molecular weight of the protein[30,31].

Next, we attempted to rebuild our model of the protein using the electron-density maps calculated using the model phases derived from molecular replacement. We immediately noticed that a cavity in the protein surface, inherent to the designed topology, was occupied with a large electron density feature present in both 2mFo-DFc and mFo-DFc electron density maps. This feature was interpreted as phosphatidylethanolamine, an *E. coli* phospholipid that we suspect binds to the protein during recombinant expression and remains bound throughout the purification. In addition to modeling the bound phospholipid, we rebuilt parts of the design model using the initial electron-density maps calculated from molecular replacement. We then performed additional, iterative refinement of atomic positions, individual atomic displacement parameters (B-factors) with a TLS model, and occupancies, using riding hydrogen atoms and automatic weight optimization, until the model reached convergence. Our tentative modeling of phosphatidylethanolamine bound to the protein was supported by the overall flatness of mFo-DFc electron density maps in the vicinity of the ligand, and by the reduction in R-free obtained when adding the ligand to the model. All model building was performed using Coot[32] and refinement steps were performed with phenix.refine (v1.16-3549) within the PHENIX suite[29,32]. Restraints for the phosphatidylethanolamine (PEE) ligand were calculated using phenix.elbow[33]. The final model coordinates were deposited in the Protein Data Bank (PDB[34]) under accession code 6W90. Further information regarding model building and refinement is presented in **Table S5**.

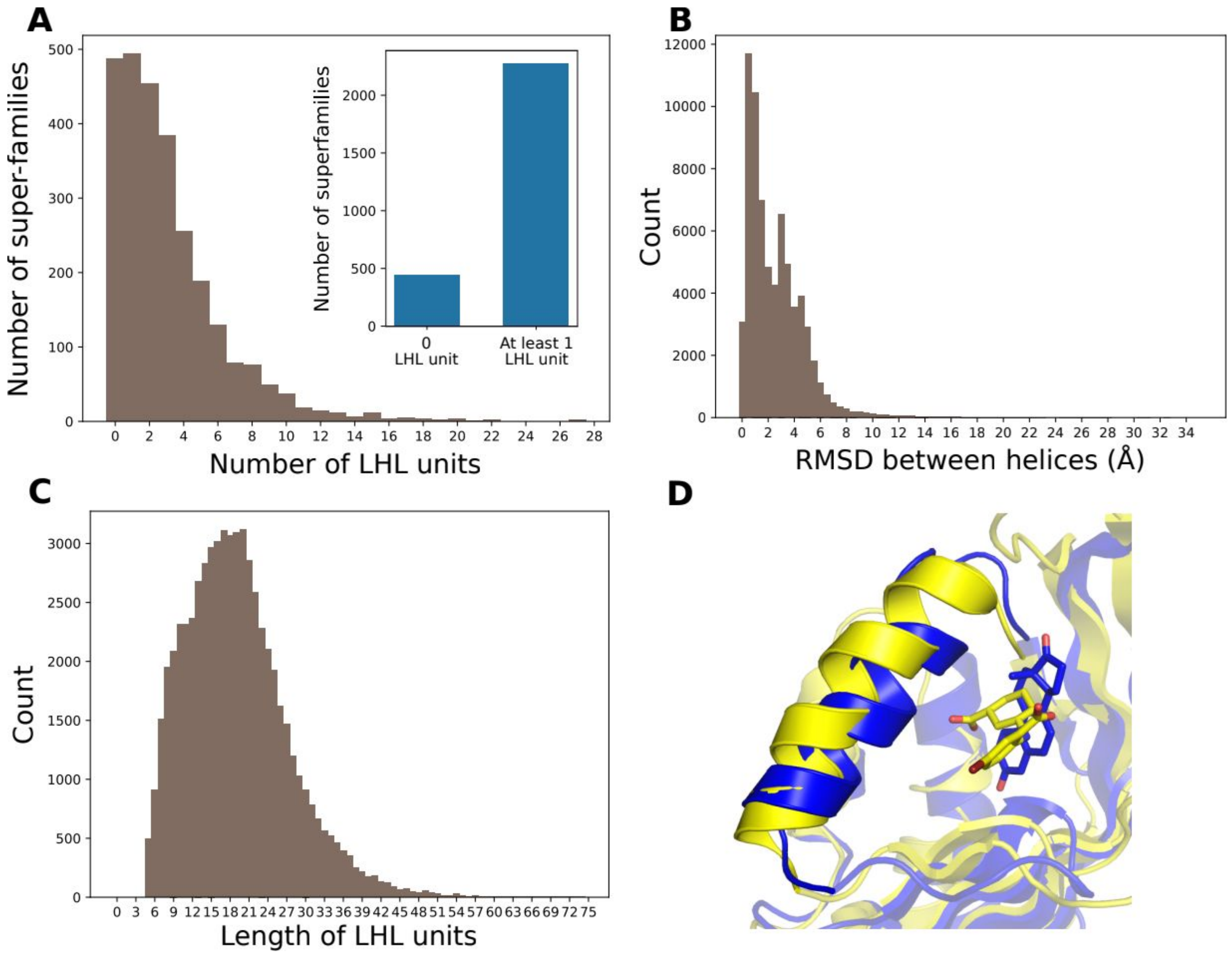


**Fig. S1. Diversity of geometries in naturally occurring fold topologies to enable distinct functions**.

**A.** Distribution of the number of LHL units contained in each CATH protein superfamily. Inset: numbers of superfamilies that have no or at least one LHL units. 83.7% of all CATH superfamilies have at least one LHL unit. **B.** Diversity of the geometries of LHL units in CATH superfamilies. Shown are the helix RMSDs between all pairs of LHL units in the same clusters computed as described in Supplementary Methods. **C.** Distribution of LHL lengths in all CATH structure superfamilies. **D.** Example where a change of the LHL geometry at the active site alters ligand specificity. The blue LHL element is from ketosteroid isomerase (PDB:1OH0) that binds a equilenin and the yellow LHL element is from Phenazine biosynthesis protein A/B that binds a 5-bromo-2-{[(1S,3R)-3-carboxycyclohexyl]amino}benzoic acid (PDB:3JUM).

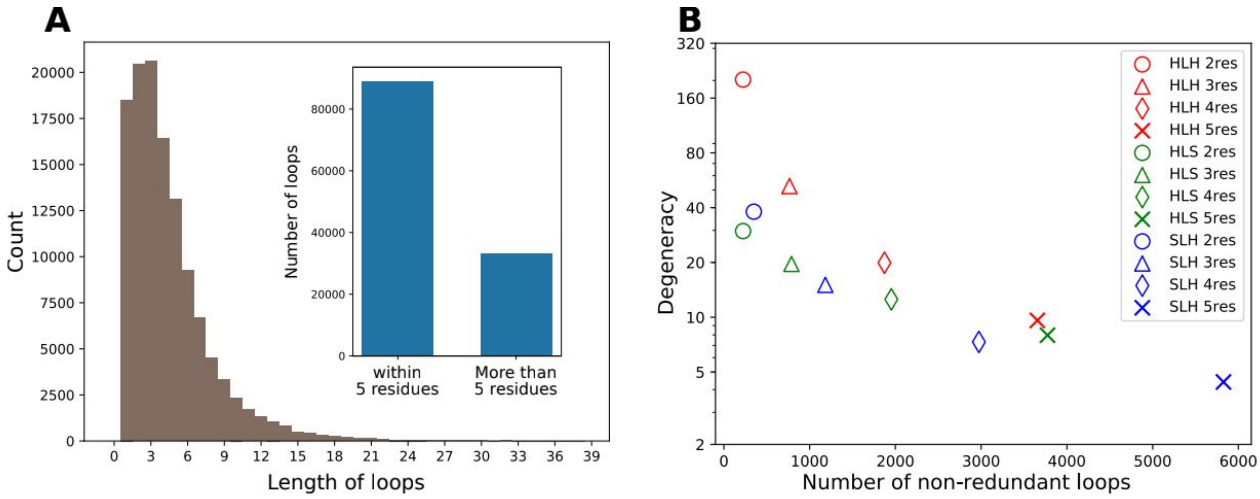


**Fig. S2. Common loop connector elements in naturally occurring LHL units.**

**A.** Distribution of loop lengths in LHL units. Inset: numbers of loops that have at most 5 residues or more than 5 residues. **B.** Sizes of non-redundant loop libraries versus the degeneracy defined as the total number of loops divided by the number of non-redundant loops. Loops connecting two helices are shown in red. Loops connecting a helix and a strand are shown in green. Loops connecting a strand and a helix are shown in blue. Loops with different numbers of residues are indicated by different markers.

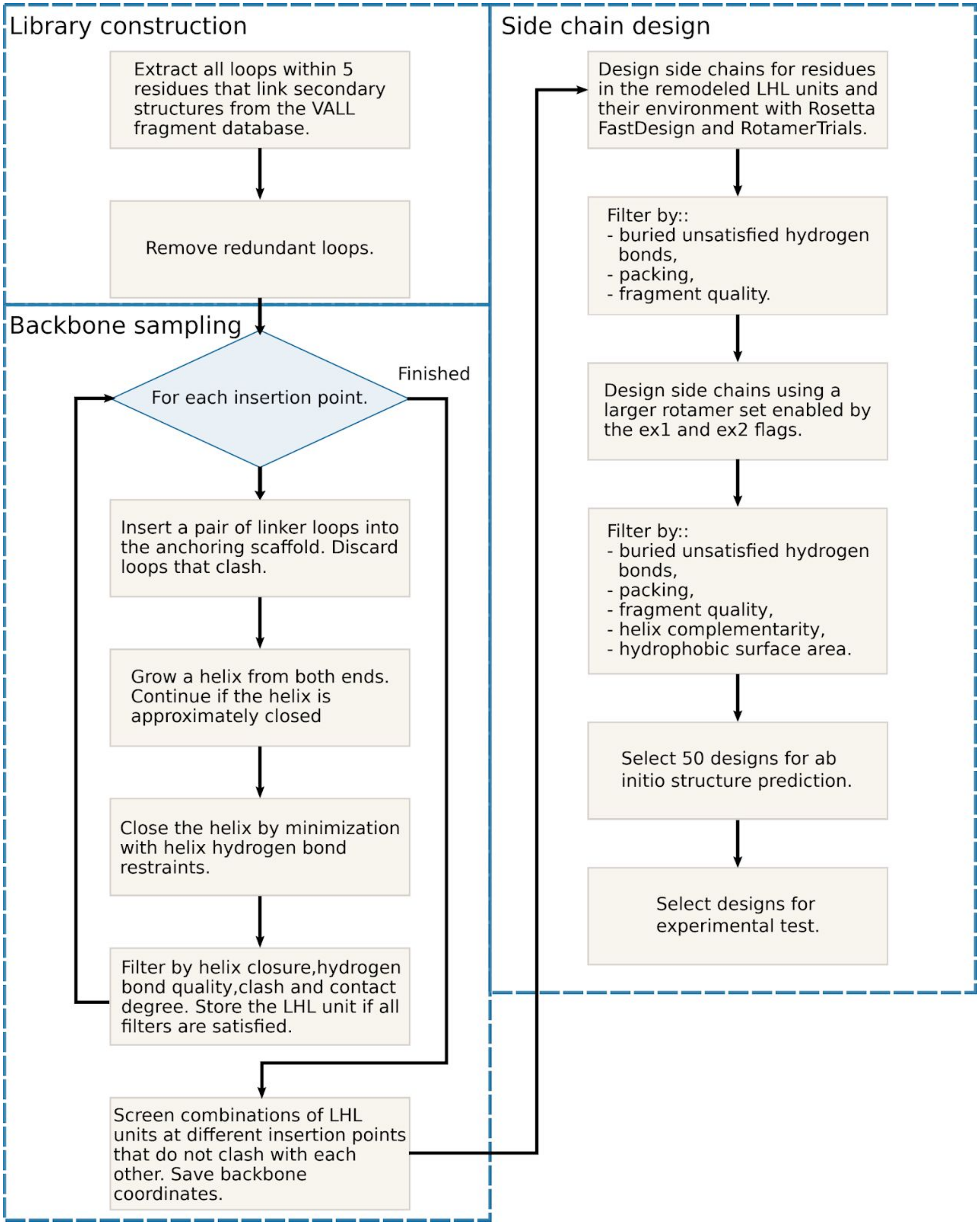


**Fig. S3.**
**Detailed flowchart of the LUCS design protocol.**

RO1_1

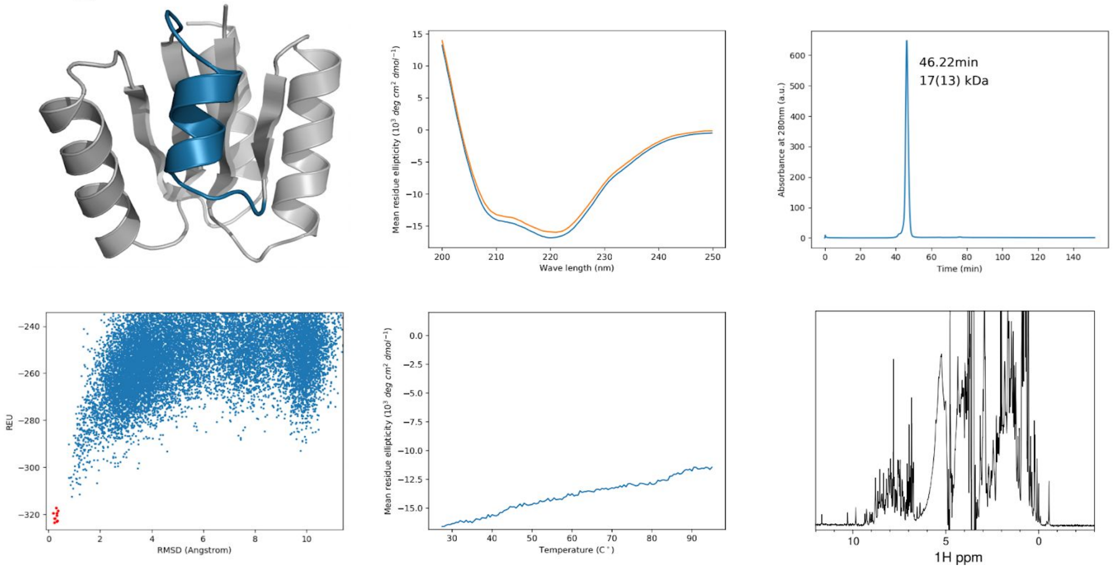


RO1_2

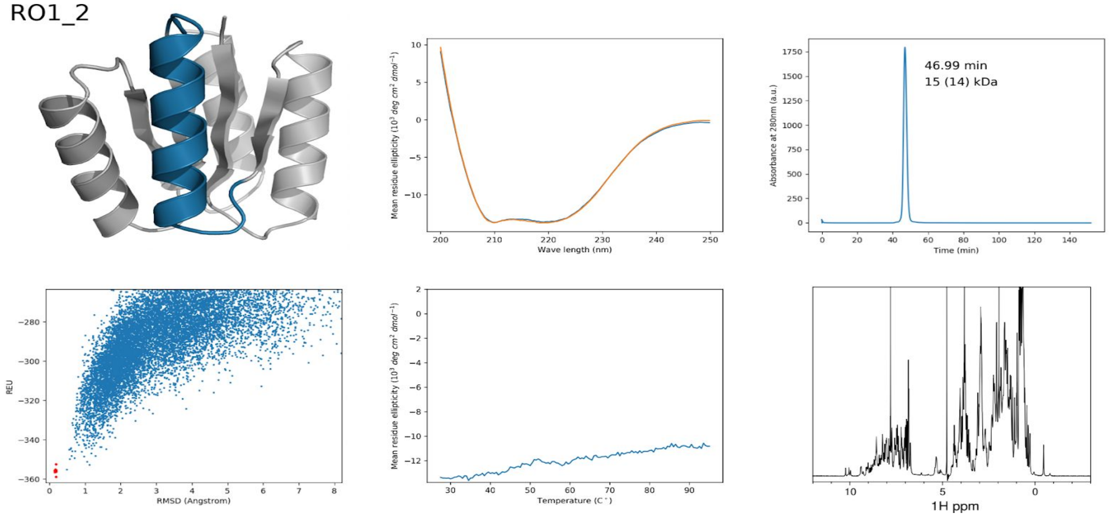


**Fig. S4. Characterization of well-folded designs.** For each design, the design model is shown in the upper left panel with the reshaped LHL units shown in blue. The lower left panels show the results of Rosetta *ab initio* structure prediction simulations. Each blue point represents a model from the prediction simulations and each red point represents a relaxed design model. For the NTF2 fold designs, the *ab initio* structure prediction simulations used a biased fragment set (see Supplementary Methods). The upper middle panels are the CD spectra at 25°C before (blue) and after measuring a melting curve (orange). The lower panels are CD melting curves measured at 220nm. The upper right panels show the size exclusion chromatograms. The peak positions and molecular weights calculated from peak positions (molecular weights calculated from amino acid sequences) are shown next to the monomer peak. For designs RO1_8, RO1_9, RO2_5, RO2_6, RO2_9, RO2_10, RO2_20 and RO2_25, the chromatograms were measured using samples purified by gel filtration. The chromatograms for the remainder of the designs were measured directly after His-tag purification. The lower right panels show the 1D-1H NMR spectra.

RO1_5

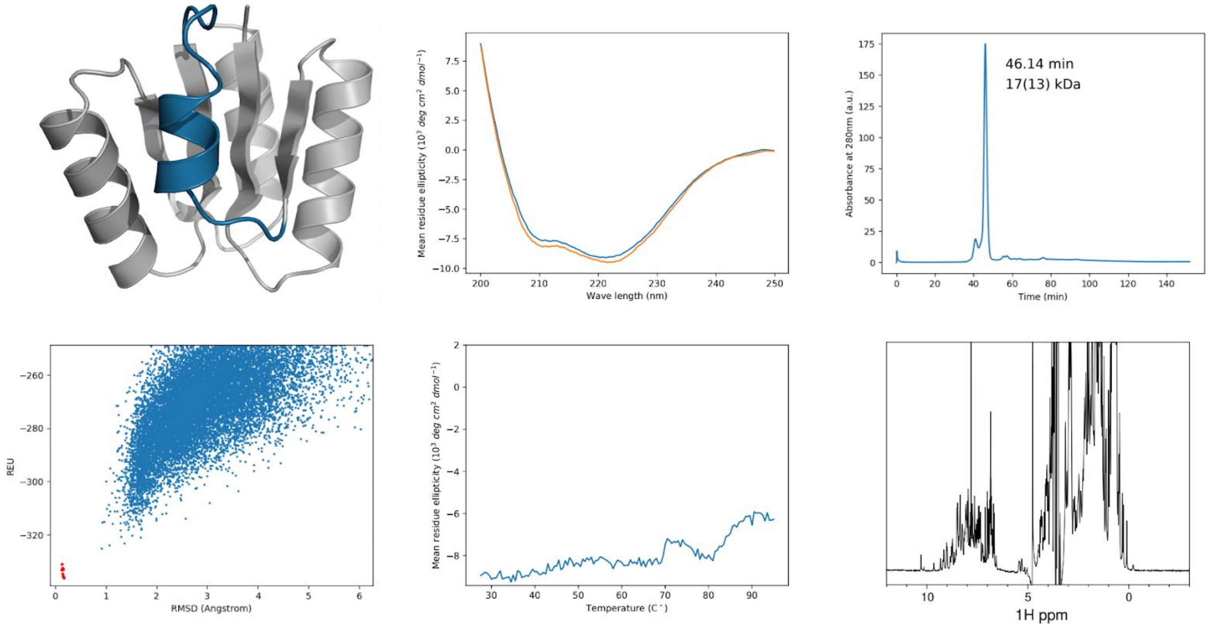


RO1_8

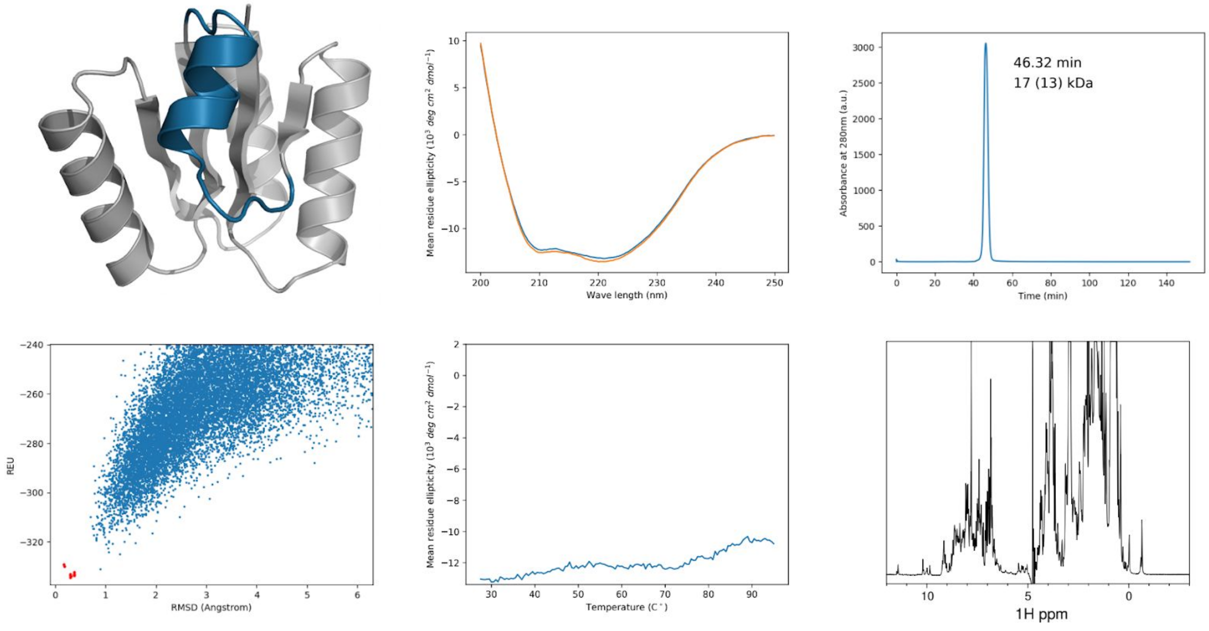


**Fig. S4. Characterization of well folded designs cont.**

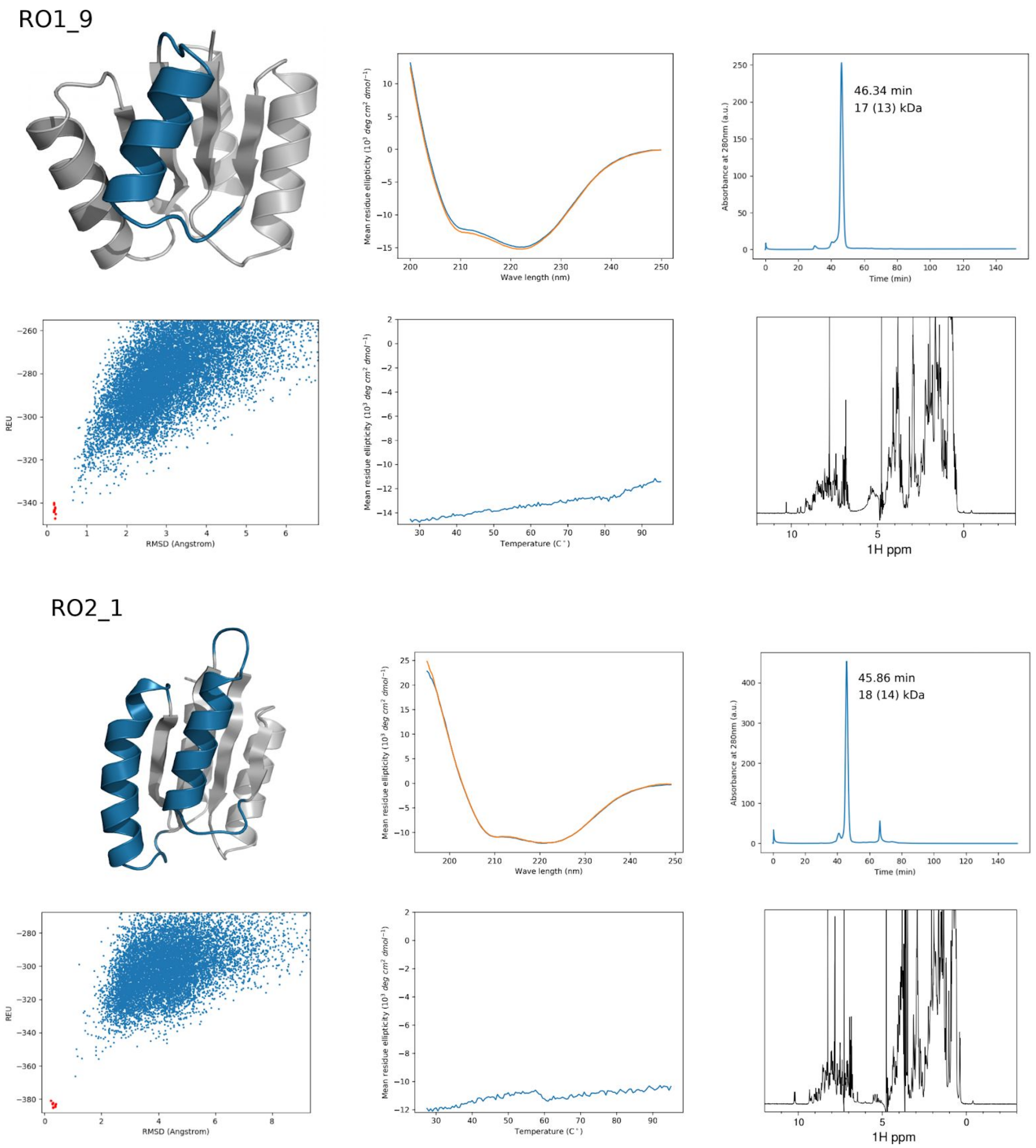


**Fig. S4. Characterization of well folded designs cont.**

RO2_5

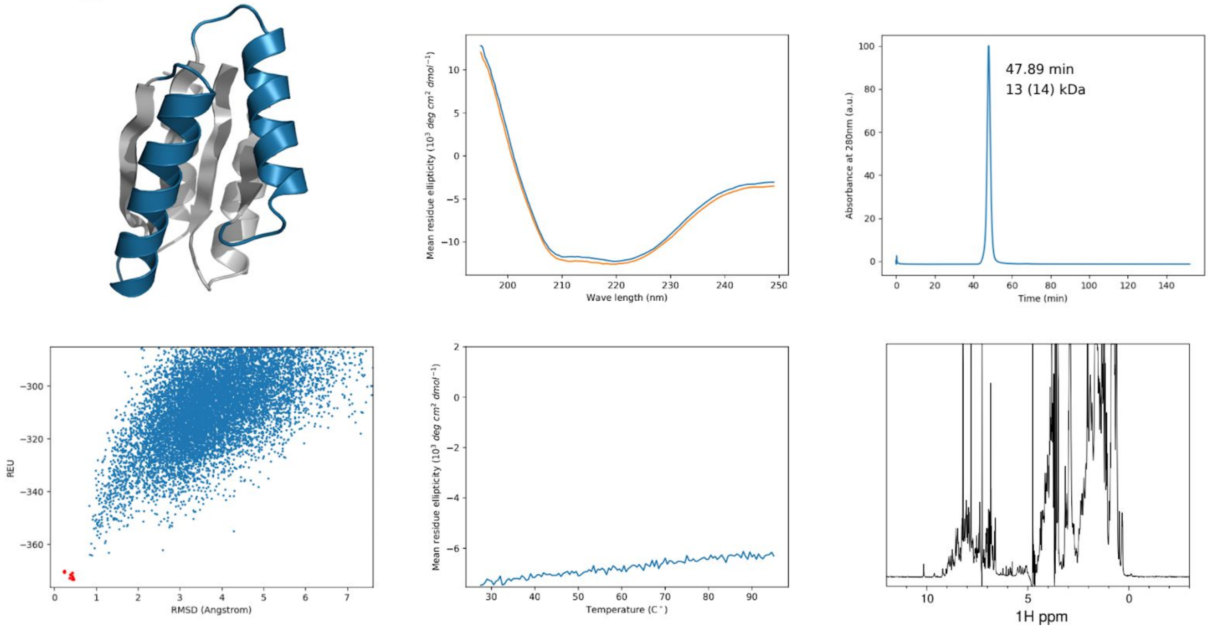


RO2_6

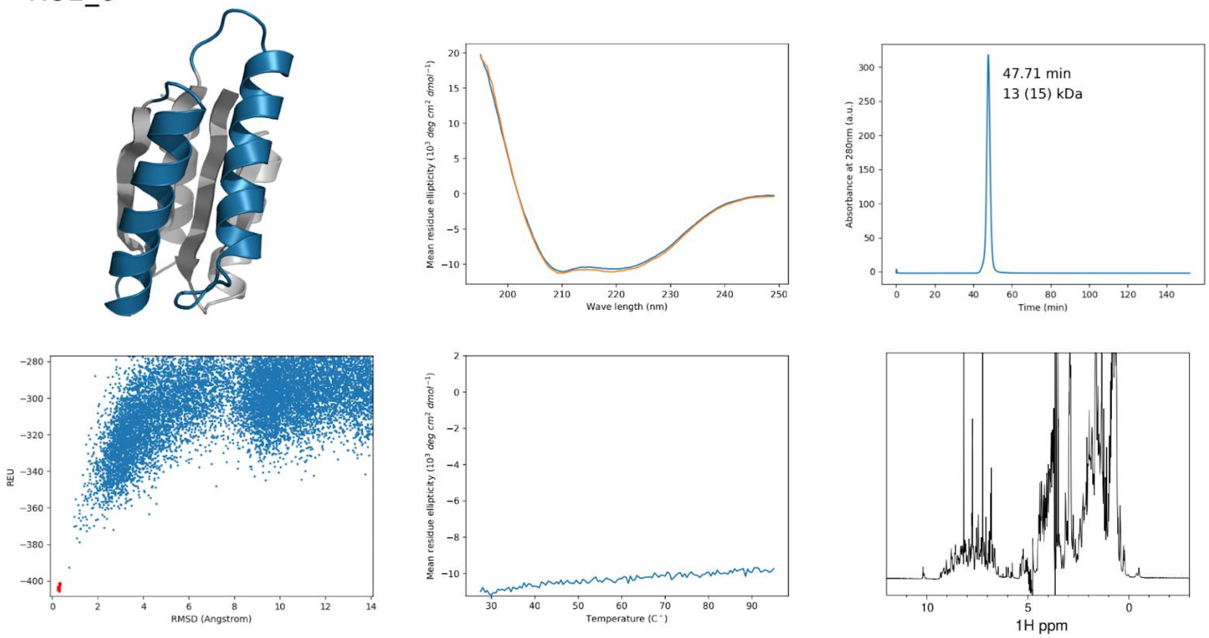


**Fig. S4. Characterization of well folded designs cont.**

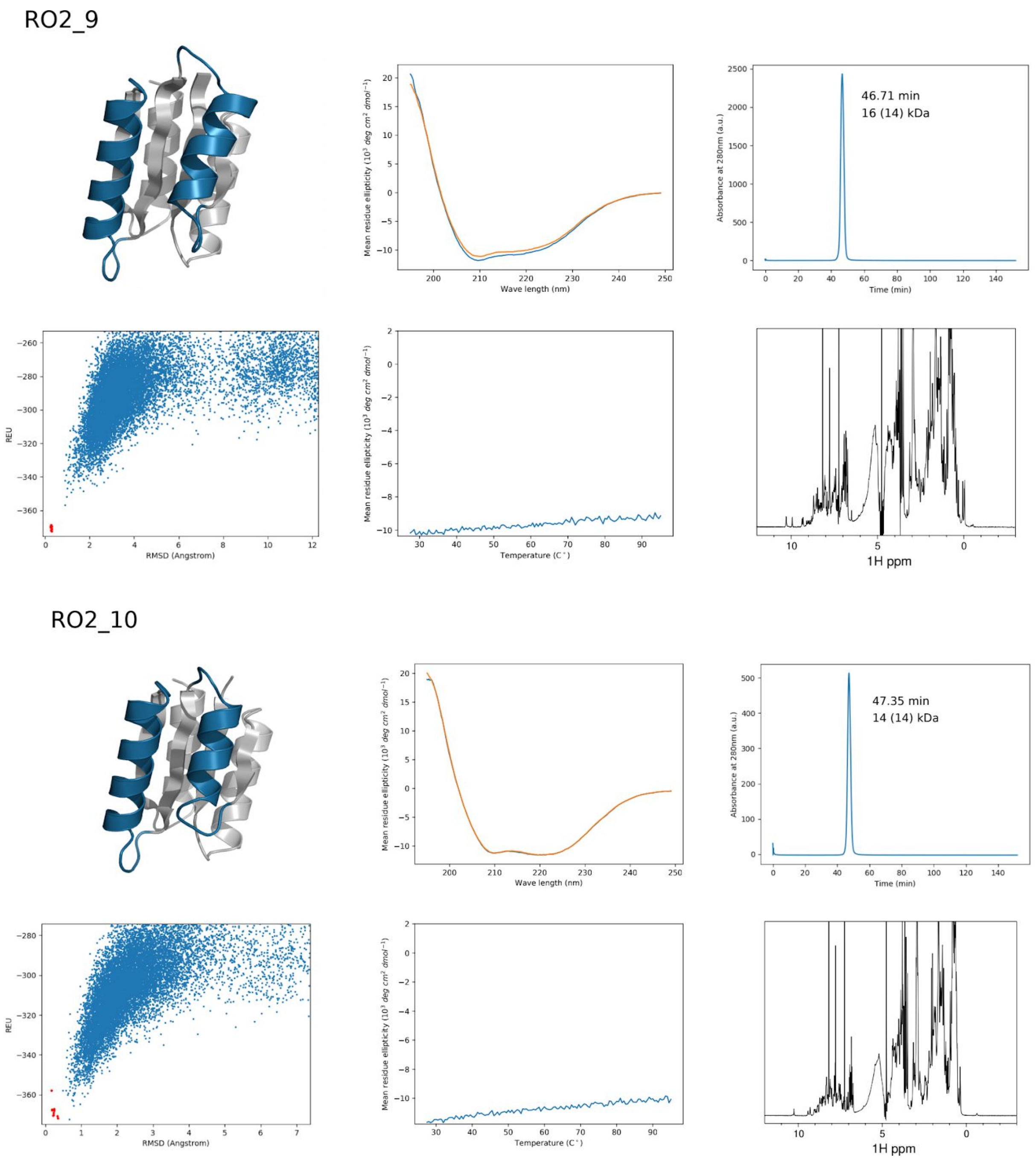


**Fig. S4. Characterization of well folded designs cont.**

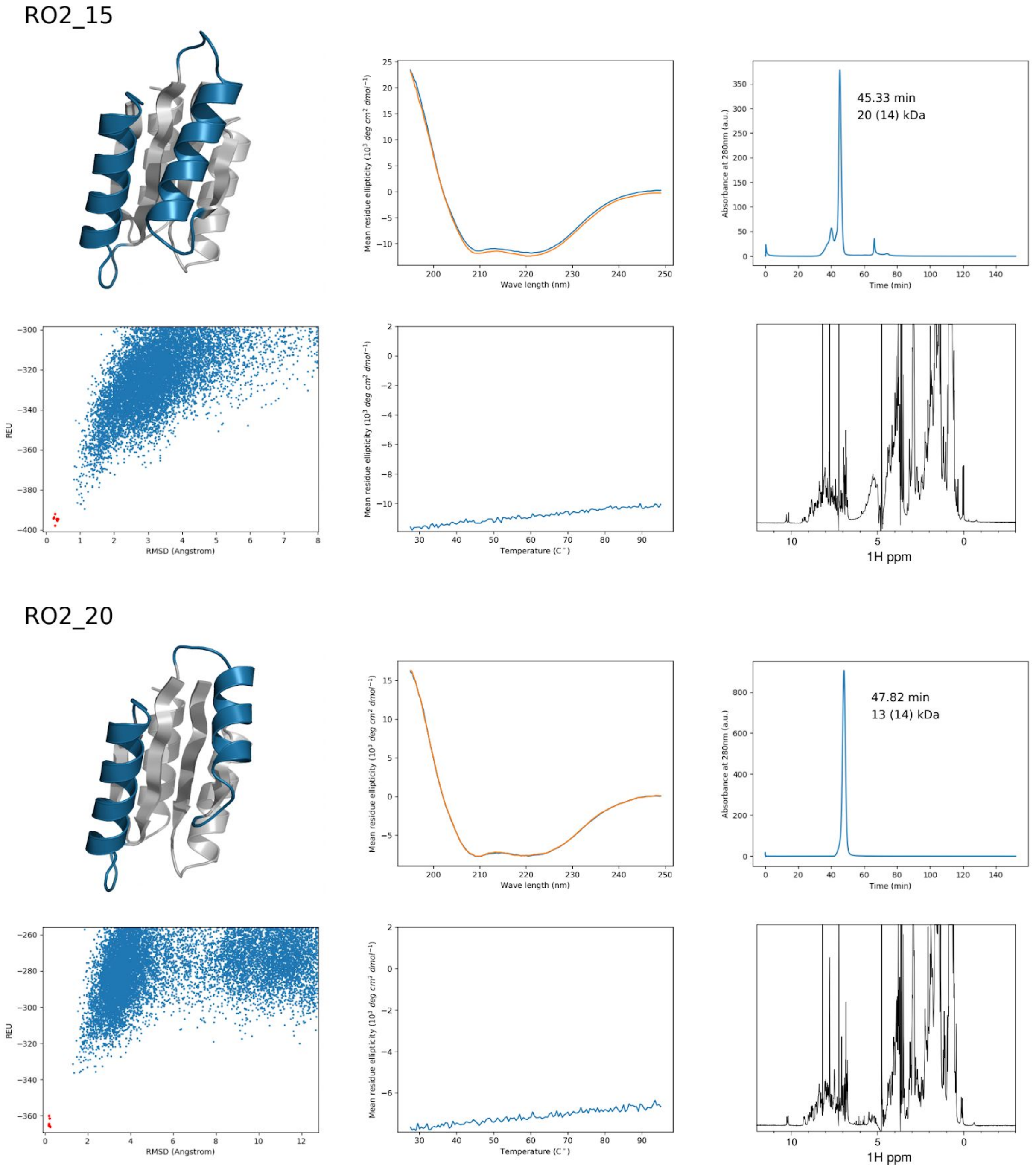


**Fig. S4. Characterization of well folded designs cont.**

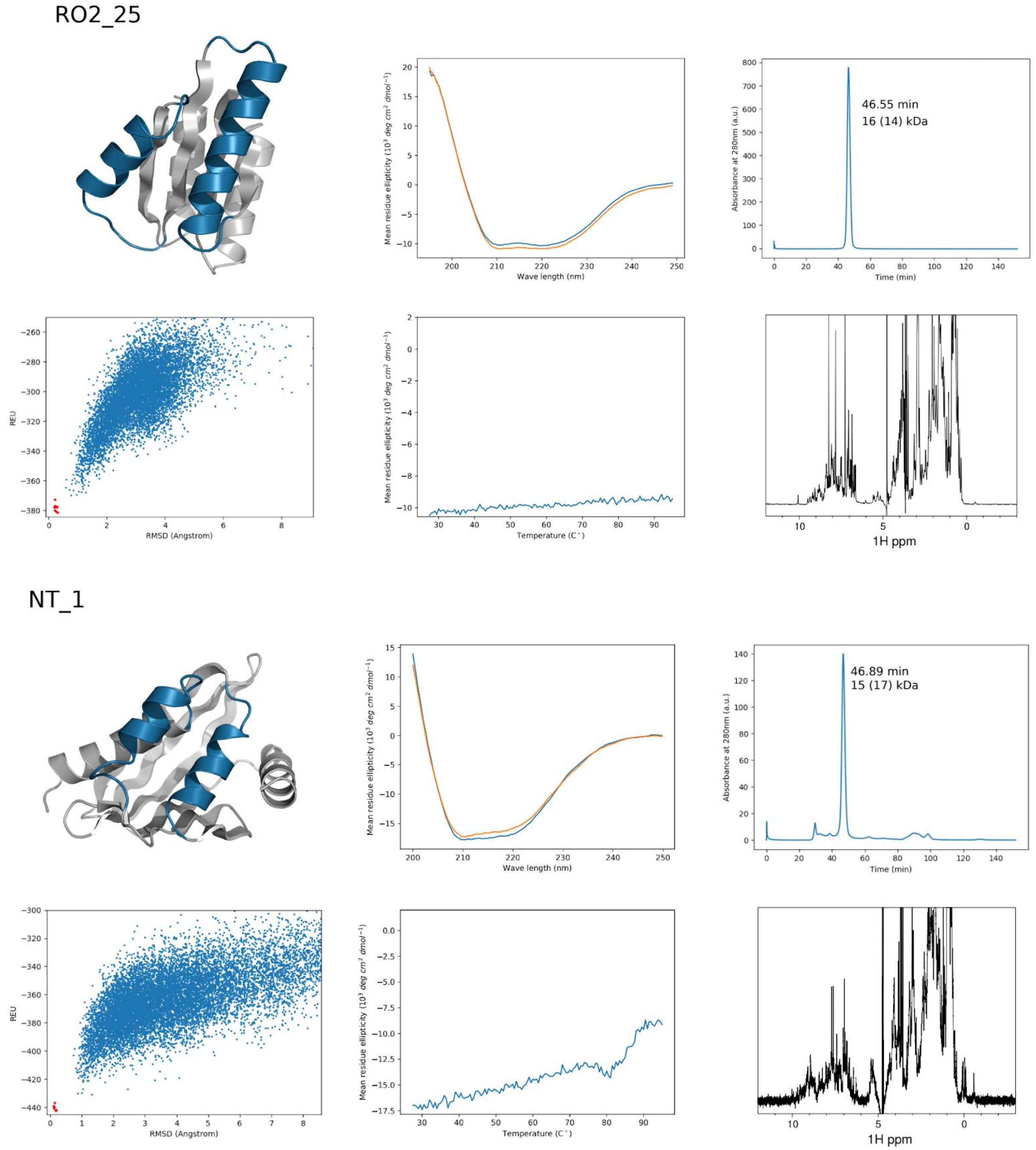


**Fig. S4. Characterization of well folded designs cont.**

NT_8

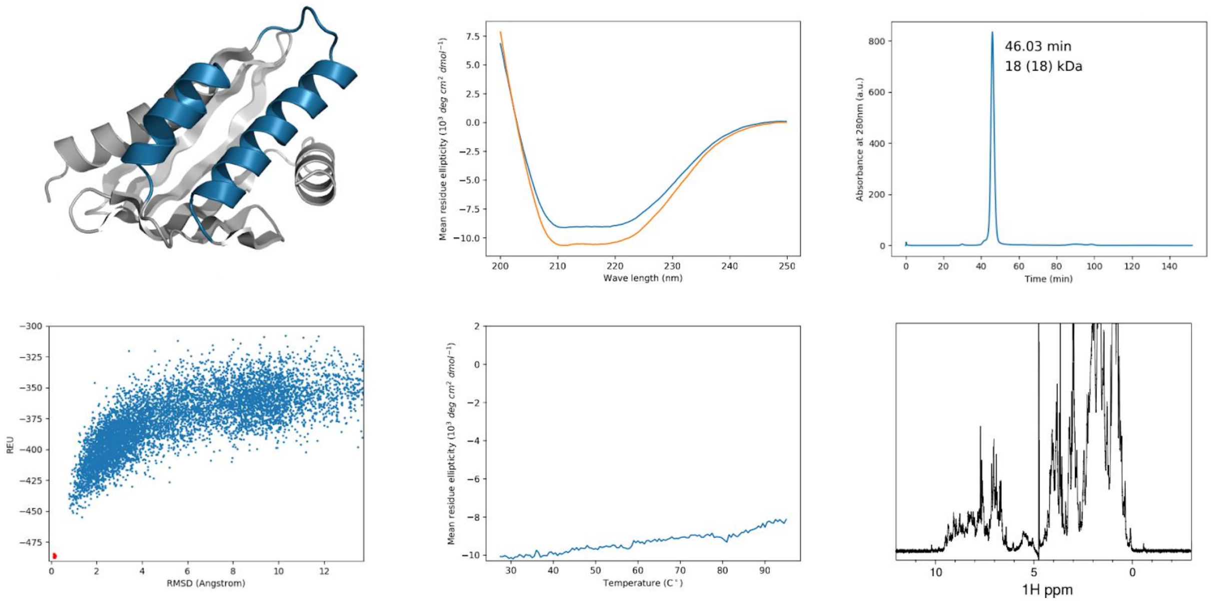


NT_9

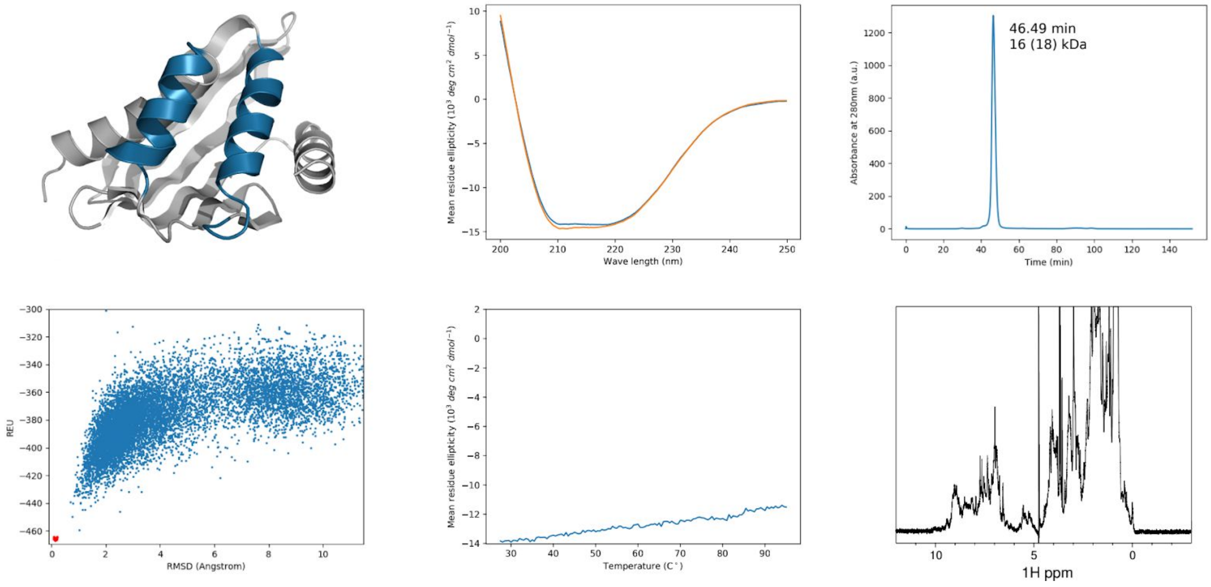


**Fig. S4. Characterization of well folded designs cont.**

NT_10

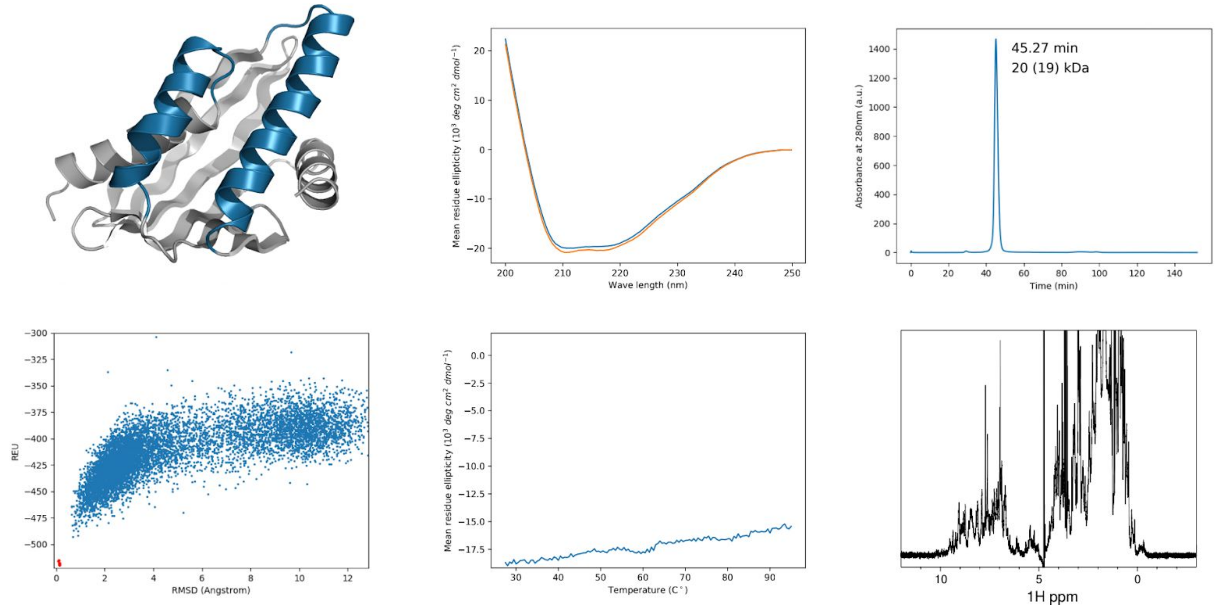


**Fig. S4. Characterization of well folded designs cont.**

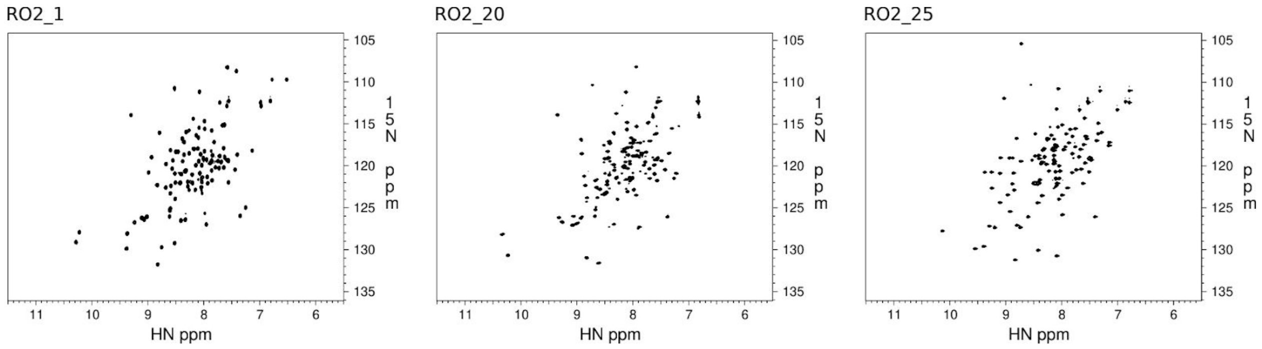


**Fig. S5.** $^{15}N$-$^{1}H$ HSQC spectra of designs whose structures were solved by NMR.

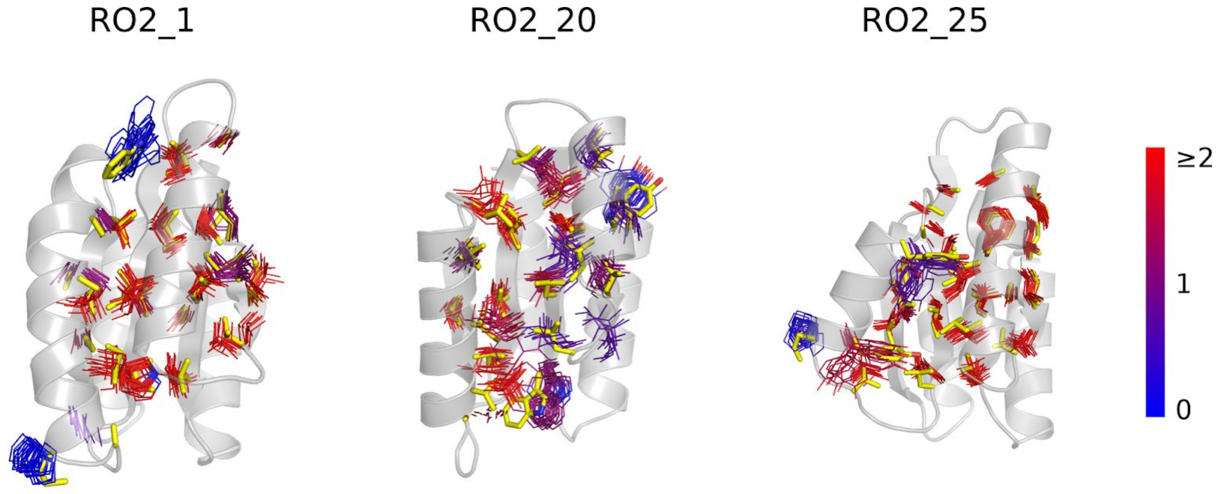


**Fig. S6. Conformations of core side chains overall agree between designed models and structures solved by NMR.** The backbones of the designed models are shown as cartoons. The hydrophobic core side chains of models are shown as yellow sticks. The conformations of the side chain in the NMR models are shown as lines and colored according to the ratio of the number of long range (distance in primary sequence larger than 3) NOEs involving its side chain atoms over the number of its side chain atoms.

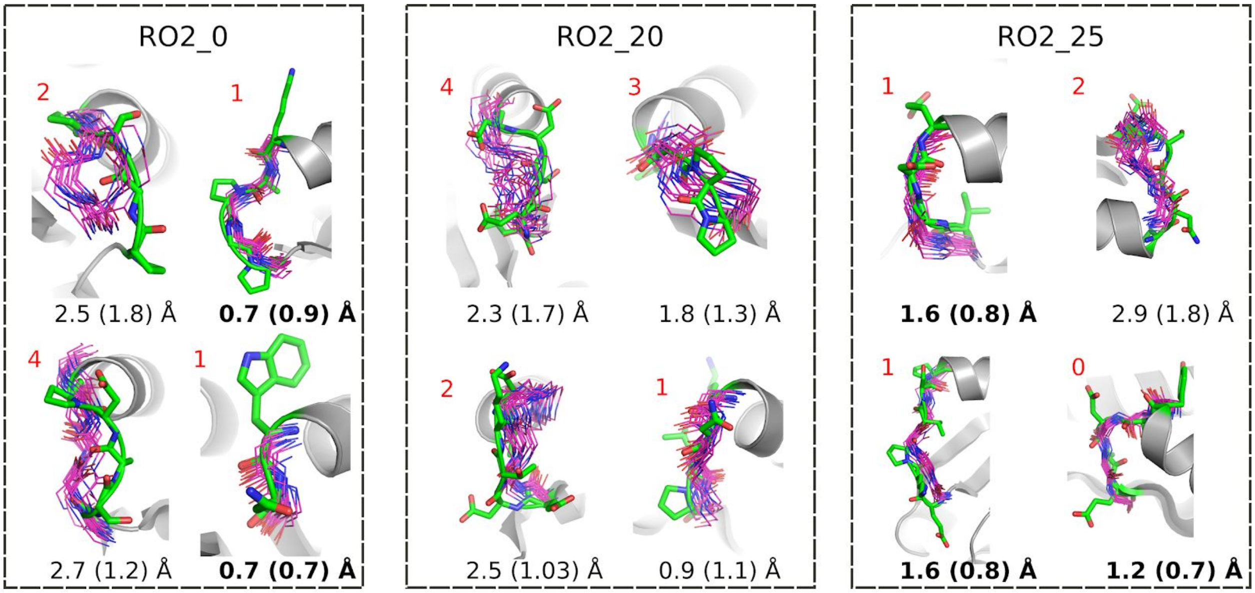


**Fig. S7. Comparison of loop conformations between designed models and structures solved by NMR.** The loops of designed models are shown as green sticks. The backbone heavy atoms of experimentally solved loops are shown in magenta. The backbone heavy atom RMSDs between the designs and lowest energy NMR models are shown below each loop and the max loop residue ensemble backbone RMSDs calculated by CCPN analysis are shown in parentheses. The 5 converged loops (pairwise backbone RMSD within the ensemble of NMR models within 1 Å) have RMSDs within 1.6Å (shown in bold). The number of residues that lack long range (distance in primary sequence larger than 3) NOE restraints are shown at the top left of each loop (red).

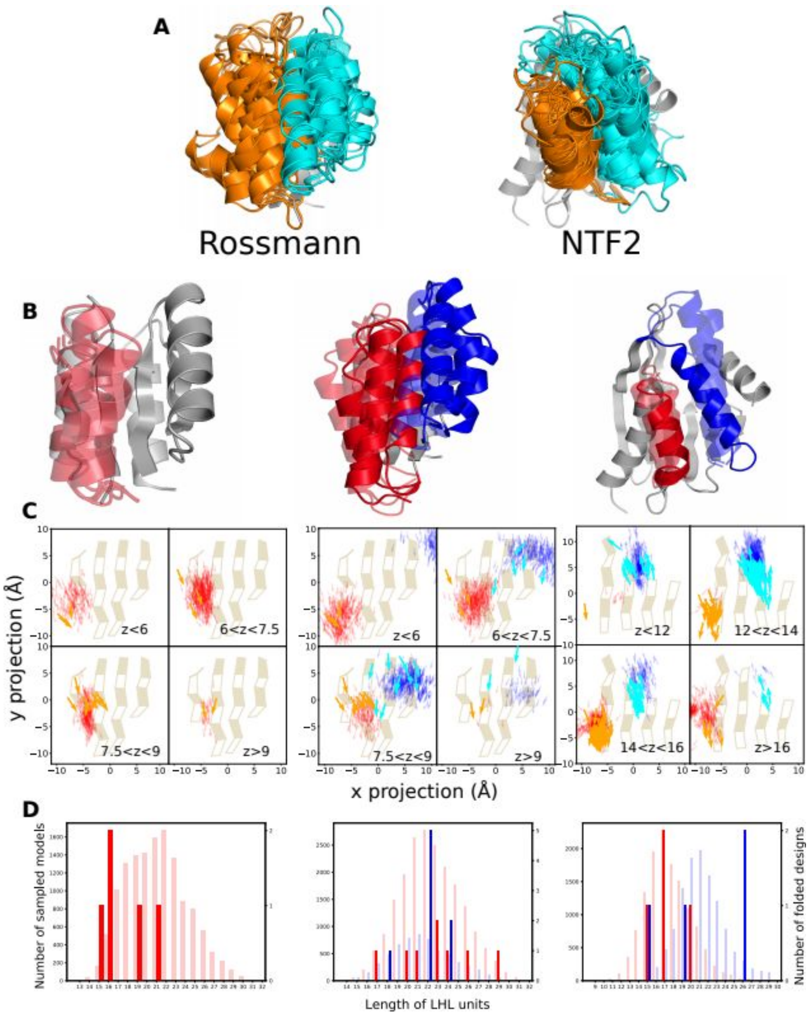


**Fig. S8. Distributions of LHL geometries in designed *de novo* fold families**. **A.** Structures of native Rossmann fold (left) and NTF2 fold (right) proteins (the two helices corresponding to the designed regions are shown in orange and cyan). The Rossmann fold structures are from the CATH superfamily 3.40.50.1980 and the NTF2 fold structures are from the CATH superfamily 3.10.450.50. In **B**,**C** and **D**, columns show the 3 design problems: Left, Rossman fold with one designed LHL unit; middle, Rossmann fold with two designed LHL units; right: NTF2 fold with two designed LHL units. **B.** Superimposition of well-folded designs. The designed LHL units are colored in red or blue. Designs with experimentally solved structures are shown in solid colors and all other well-folded designs are shown in transparent colors. **C.** Projection of centers and directions of helices from designable LUCS models (red, blue) and known structures (orange, cyan) onto the underlying beta sheets, as shown in **Fig.3** in the main text but overlaid. **D.** The distributions of LHL lengths of the sampled models (light colors) and the well-folded designs (dark colors).

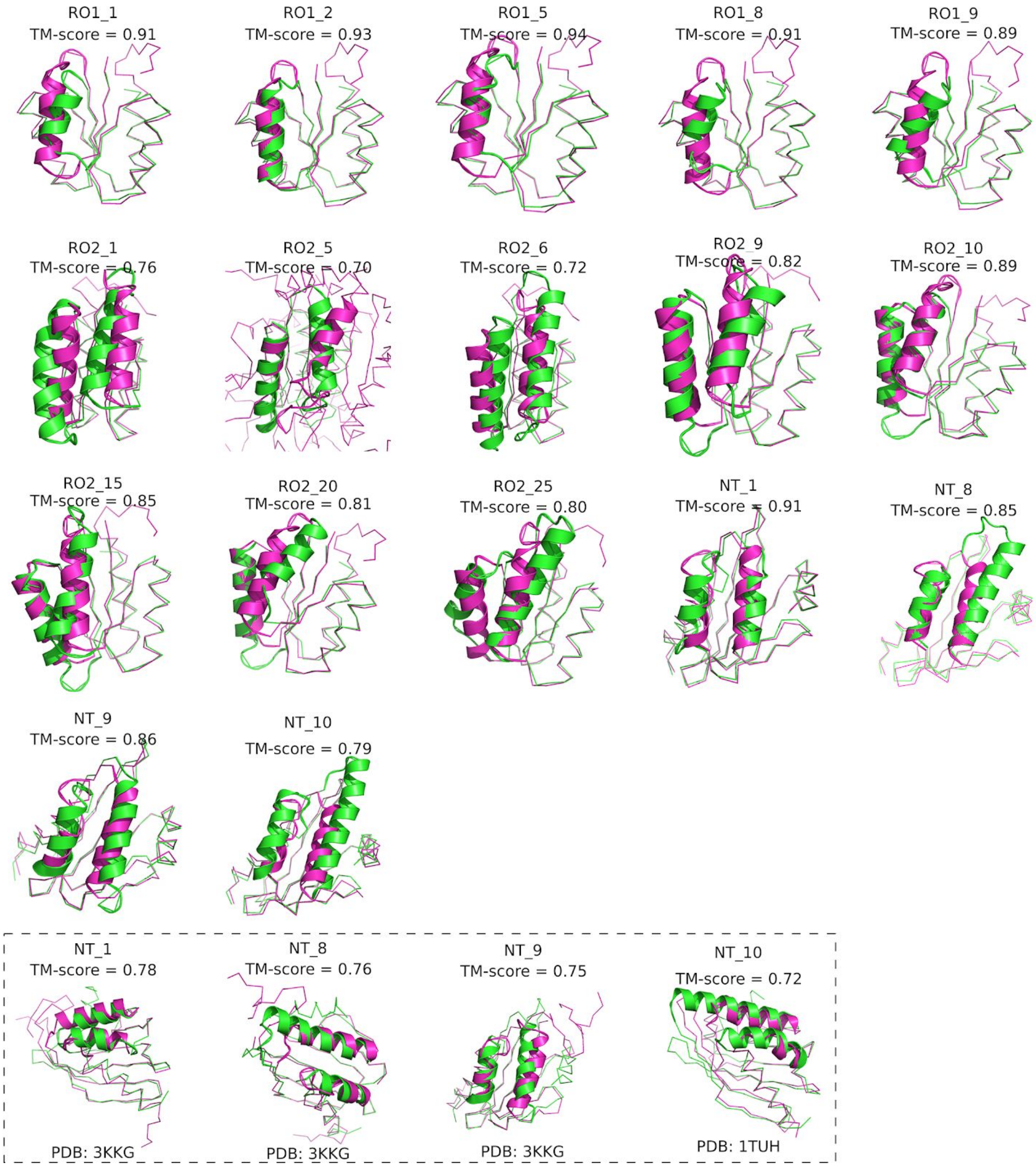


**Fig. S9. Models of well-folded designs show considerable geometry differences to their best matches from TM-align**. Design models are shown in green and the matched structures are shown in magenta. Except for RO2_5, all Rossmann fold designs matched best to PDB:2KPO which is the *de novo* designed input scaffold PDB:2LV8. All natural proteins in the top 10 TM-align matches to Rossmann fold designs have different topologies from the designed topology. All NTF2 fold designs matched best to the *de novo* designed input scaffold PDB:5TPJ. The last row shows the best natural protein matches to the NTF2 fold designs.

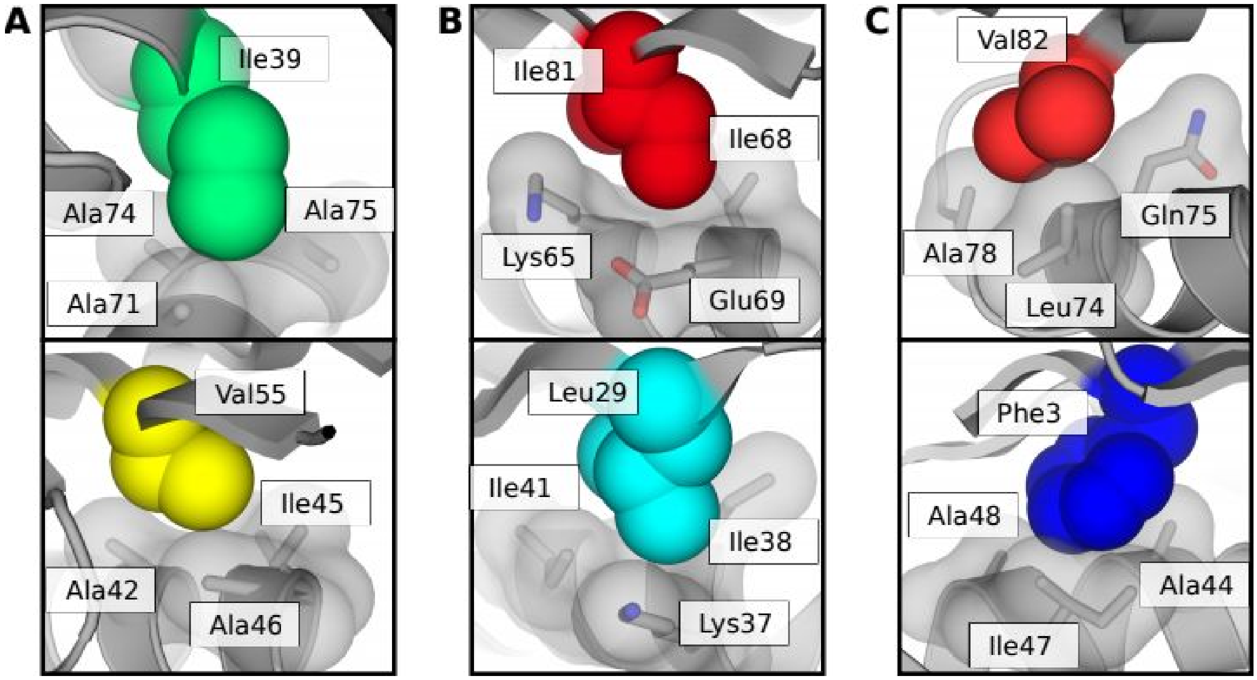


**Fig. S10. Residues that form knobs-into-sockets packing.** The knob residues shown as colored spheres and the socket residues are shown as grey sticks with transparent surfaces. **A.** RO2_1, **B.** RO2_20 **C.** RO2_25.

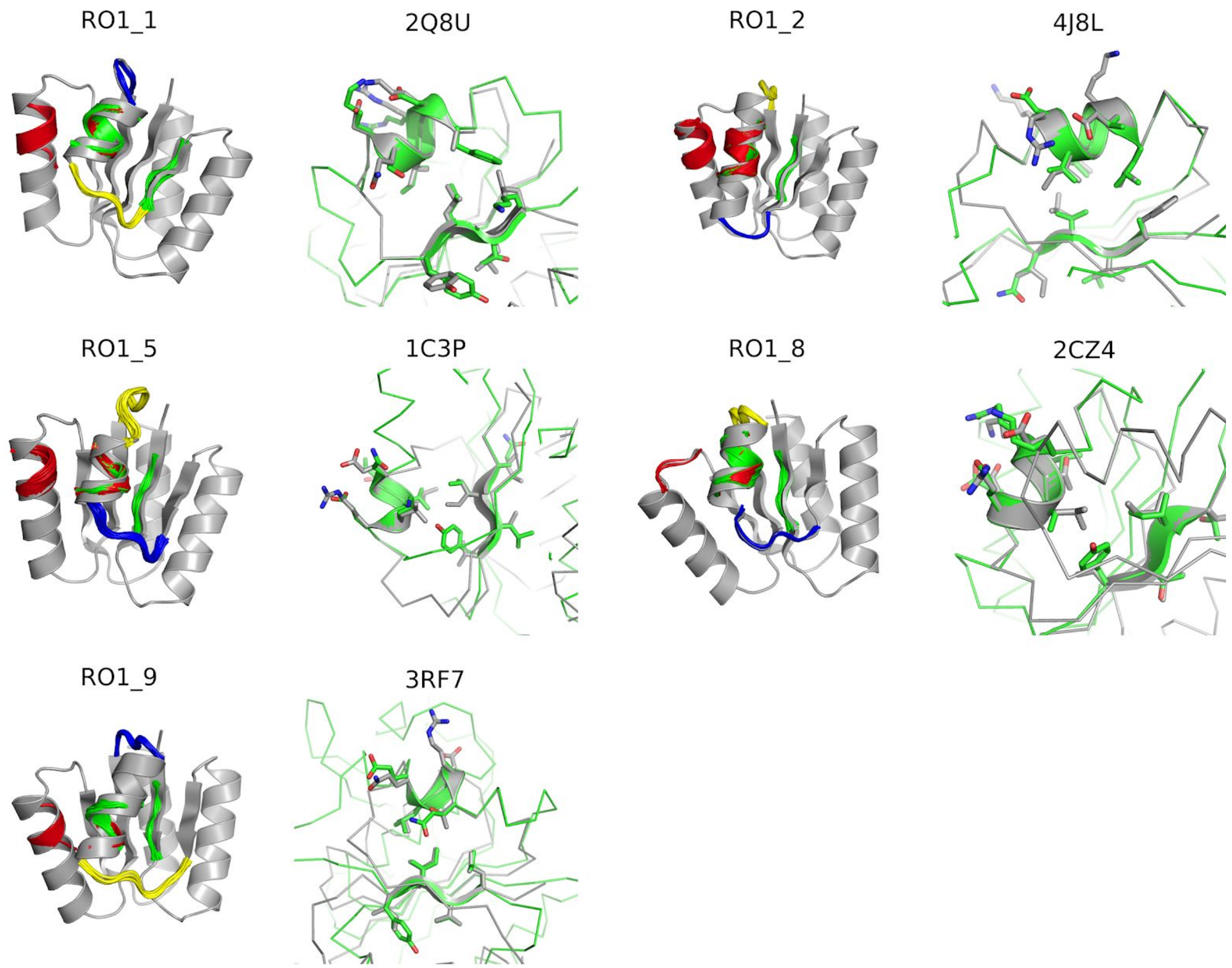


**Fig. S11. Identification of tertiary motifs for designed models.** For each design, the left panel shows the designed model in grey and the matched tertiary motifs in color. The right panel shows the top match of the green motif. The designs are colored in grey and the source structure of one of the motifs in green. The two structures are superimposed at the motif region. The backbones of the matched region are shown as cartoons and the side chains as sticks. The PDB code of the motif source structures are shown at the top of the right panels.

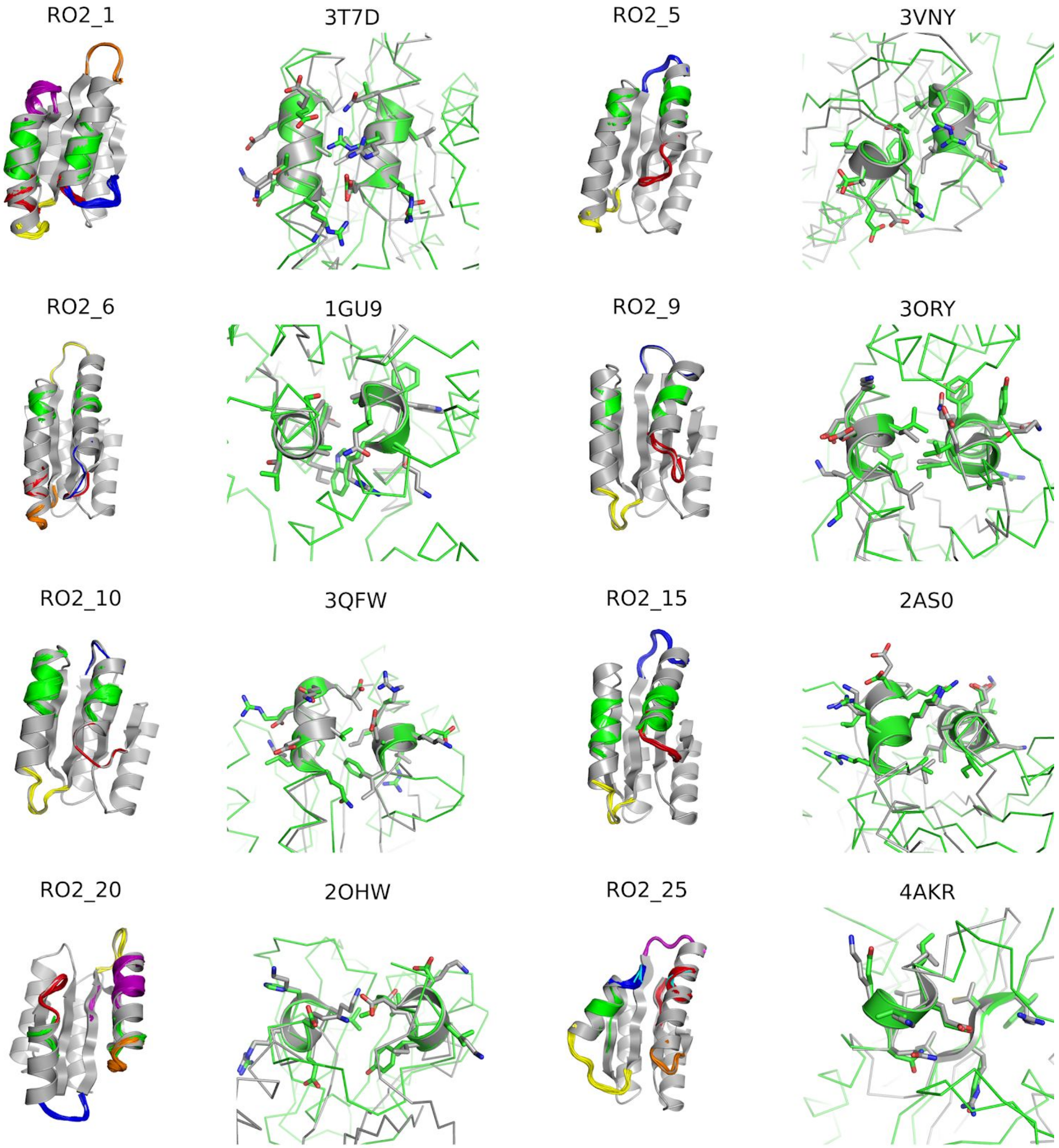


**Fig. S11. Identification of tertiary motifs for designed models cont.**

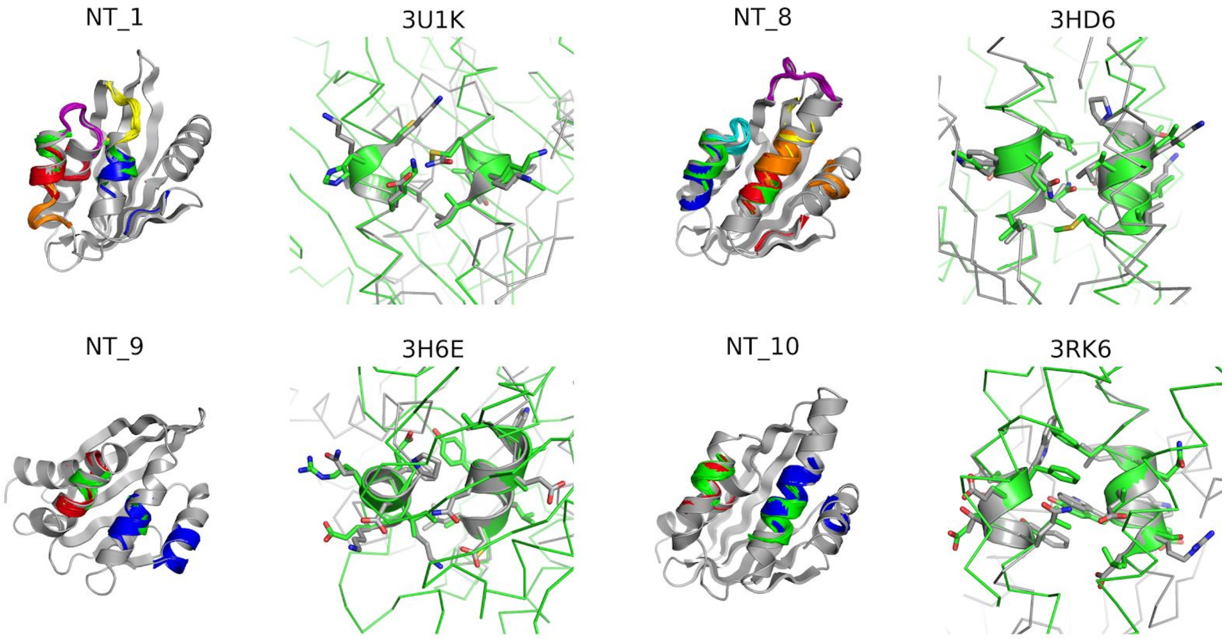


**Fig. S11. Identification of tertiary motifs for designed models cont.**

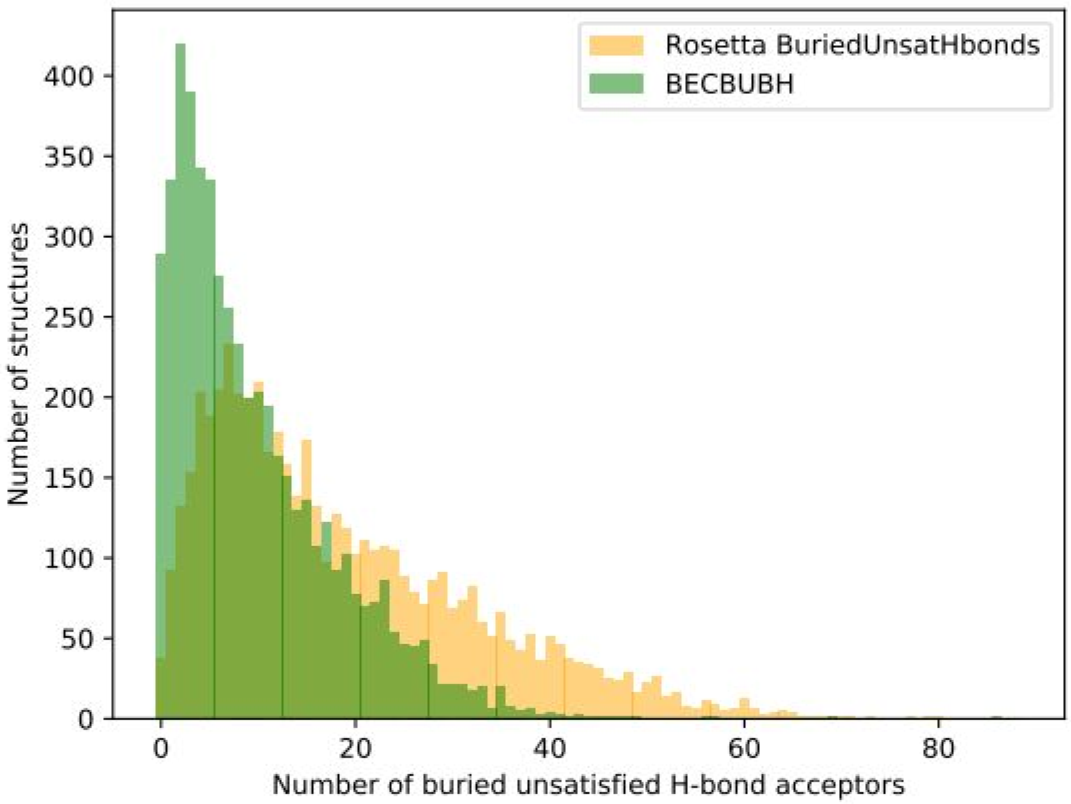


**Fig. S12. Number of buried unsatisfied hydrogen bond acceptors found in the top8000 protein structure data set** (http://kinemage.biochem.duke.edu/databases/top8000.php). The unsatisfied acceptors found by the Rosetta BuriedUnsatHbonds filter are shown in yellow and the unsatisfied acceptors found by the BECBUBH filter are shown in green.

**Table S1. Loop libraries.**

| Type | Residue number | Number of all loops | Number of non-redundant loops | Degeneracy |
|---|---|---|---|---|
| Helix-loop-helix | 2 | 45273 | 224 | 202 |
| | 3 | 40194 | 765 | 52.5 |
| | 4 | 37295 | 1873 | 19.9 |
| | 5 | 35118 | 3656 | 9.61 |
| Helix-loop-strand | 2 | 6663 | 224 | 29.7 |
| | 3 | 15465 | 788 | 19.6 |
| | 4 | 24487 | 1953 | 12.5 |
| | 5 | 30061 | 3774 | 7.97 |
| Strand-loop-helix | 2 | 13263 | 349 | 38.0 |
| | 3 | 17819 | 1183 | 15.1 |
| | 4 | 21750 | 2974 | 7.31 |
| | 5 | 25684 | 5826 | 4.41 |

**Table S2. Number of models at each design stage.**

| Design stage | RO1 | RO2 | NT |
|---|---|---|---|
| Backbones generated | 13421 | 290836 | 136683 |
| Sequences designed (iteration 1) | 58626 | 432735 | 409101 |
| Designs passed filters (iteration 1) | 1163 | 9934 | 5715 |
| Designs selected for *ab initio* folding | 50 | n/a | n/a |
| Sequences designed (iteration 2) | | 49578 | 22831 |
| Designs passed filters (iteration 2) | | 722 | 98 |
| Designs selected for *ab initio* folding | | 50 | n/a |
| Designs selected for refinement (iteration 3) | | | 10 (manual) |
| Sequences designed (iteration 3) | | | 1000 |
| Designs passed filters (iteration 3) | | | 202 |
| Designs selected for *ab initio* folding | | | 50 |
| Designs passed *ab initio* folding | 22 | 25 | 10 |
| Designs experimentally tested | 10 | 25 | 10 |

**Table S3. Experimental characterization of designs**

| Design ID | Soluble | Relative dimer peak size in SEC* | Alpha-beta protein CD spectrum | Well resolved 1D-NMR | 15N HSQC with good dispersion |
|---|---|---|---|---|---|
| RO1_1 | Y | no dimer peak | Y | Y | |
| RO1_2 | Y | no dimer peak | Y | Y | |
| RO1_3 | N | | | | |
| RO1_4 | N | | | | |
| RO1_5 | Y | 0.11 | Y | Y | |
| RO1_6 | Y | | | | |
| RO1_7 | N | | | | |
| RO1_8 | Y | no dimer peak | Y | Y | |
| RO1_9 | Y | <0.05 | Y | Y | |
| RO1_10 | N | | | | |
| RO2_1 | Y | 0.06 | Y | Y | Y |
| RO2_2 | Y | | | | |
| RO2_3 | N | | | | |
| RO2_4 | N | | | | |
| RO2_5 | Y | no dimer peak | Y | Y | |
| RO2_6 | Y | no dimer peak | Y | Y | |
| RO2_7 | N | | | | |
| RO2_8 | N | | | | |
| RO2_9 | Y | no dimer peak | Y | Y | |
| RO2_10 | Y | no dimer peak | Y | Y | |
| RO2_11 | N | | | | |
| RO2_12 | Y | | | | |
| RO2_13 | Y | | | | |
| RO2_14 | Y | | | | |
| RO2_15 | Y | 0.15 | Y | Y | |
| RO2_16 | N | | | | |
| RO2_17 | N | | | | |
| RO2_18 | N | | | | |
| RO2_19 | N | | | | |
| RO2_20 | Y | no dimer peak | Y | Y | Y |
| RO2_21 | N | | | | |
| RO2_22 | N | | | | |
| RO2_23 | Y | | | | |
| RO2_24 | Y | | | | |

| | | | | | |
|---|---|---|---|---|---|
| RO2_25 | Y | no dimer peak | Y | Y | Y |
| NT_1 | Y | 0.09 | Y | Y | |
| NT_2 | N | | | | |
| NT_3 | N | | | | |
| NT_4 | N | | | | |
| NT_5 | N | | | | |
| NT_6 | N | | | | |
| NT_7 | N | | | | |
| NT_8 | Y | <0.05 | Y | Y | |
| NT_9 | Y | <0.05 | Y | Y | |
| NT_10 | Y | <0.05 | Y | Y | |

### Table S4. NMR statistics

| | RO2_1 | RO2_20 | RO2_25 |
|---|---|---|---|
| Number of residues | 105 | 100 | 104 |
| **Distance restraints** | | | |
| Total NOE | 1288 | 1542 | 2326 |
| intra-residue [i = j] | 385 | 455 | 577 |
| sequential [\| i - j \| = 1] | 375 | 426 | 595 |
| medium range [1 < \| i - j \| < 5] | 203 | 305 | 492 |
| long range [\| i - j \| ≥ 5] | 325 | 356 | 662 |
| Hydrogen bonds | 36 | 96 | 36 |
| Dihedral | 226 | 222 | 230 |
| **Violations** | | | |
| Distance constraints (Å) | 0.008±0.001 | 0.02±0.002 | 0.014±0.0009 |
| Dihedral angle constraints (°) | 1.1±0.7 | 1.3±0.1 | 2.2±0.17 |
| Max. distance constraint violation (Å) | 0.38 | 0.37 | 0.46 |
| Max. dihedral angle restraint violation (°) | 14.4 | 14.2 | 15.8 |
| Num. Distance violations > 0.3 Å | 0.1±0.4 | 0.6±0.7 | 1.1±0.7 |
| Num dihedral violations between 5-10° | 2.3±2 | 3.6±1 | 5.6±2 |
| Num dihedral violations > 10° | 0.85±1 | 0.25±0.5 | 3.8±1 |
| **Validate peaks vs structures (PyRPF)** | | | |
| Recall | 0.985 | 0.95 | 0.944 |
| Precision | 0.746 | 0.687 | 0.793 |
| F-score | 0.849 | 0.798 | 0.862 |
| DP-score | 0.87 | 0.855 | 0.9 |
| **Structure validation (PSVS)** | | | |
| Deviations from idealized geometry | | | |
| Bond lengths (Å) | 0.004 | 0.004 | 0.005 |
| Bond angles (°) | 0.6 | 0.7 | 0.8 |
| Ramachandran plot | | | |
| Most favored regions | 95.20% | 90.90% | 86.80% |
| Additionally allowed regions | 4.80% | 9.10% | 12.40% |
| Generously allowed regions | 0 | 0.10% | 0.60% |
| Disallowed regions | 0 | 0 | 0.20% |
| Average pairwise r.m.s.d. (Å) | | | |
| Heavy | 1.4 | 1.4 | 1.1 |
| Backbone | 0.8 | 0.8 | 0.6 |
| Structure Quality Factors (raw/Z-scores) | | | |
| Procheck G-factor (phi/psi) | 0.13/0.83 | -0.09/-0.04 | -0.17/-0.35 |
| Procheck G-factor (all) | 0.01/0.06 | -0.15/-0.89 | -0.18/-1.06 |
| Verify3D | 0.29/-2.73 | 0.27/-3.05 | 0.26/-3.21 |
| MolProbity clashscore | 8.27/0.11 | 9.32/-0.07 | 12.46/-0.61 |

**Table S5. X-ray data reduction and model refinement.**

| | |
|---|---|
| Wavelength | 1.116Å |
| Resolution Range | 40.94-1.50 Å (1.53-1.50 Å) |
| Unit Cell | a=33.50Å , b=51.99Å , c=66.41Å<br>**α**=**β**=***γ***=90° |
| Space Group | $P2_12_12_1$ |
| Unique Reflections | 19203 (895) |
| Multiplicity | 22.7 (12.4) |
| Completeness | 99.9% (92.9%) |
| <I/σI> | 18.5 (1.8) |
| $CC_{1/2}$ | 0.999 (0.719) |
| $R_{pim}$ | 0.019 (0.283) |
| $R_{work}$ | 0.1864 |
| $R_{free}$ | 0.2104 |
| Total Refined Atoms | 1344 |
| Protein Residues | 128 |
| Solvent Molecules | 81 |
| Refined Ligand Atoms | 48 |
| Average B-factor | 33.81Å$^2$ |
| $RMSD_{bonds}$ | 0.013Å |
| $RMSD_{angles}$ | 1.31° |
| Rama. Plot: | |
| Favored | 98.44% |
| Allowed | 1.56% |
| Outliers | 0.0% |
| Molprobity Clashscore[d] | 3.84 |
| PDB ID | 6W90 |

**Data S1. Protein sequences of designs selected for experimental testing.**

| Design ID | Protein Sequence |
|---|---|
| RO1_1 | KLVVVIDSNDKKLIEEAKKMAEKANLLLLYDVDEDQVRKAAGNARILVLVSNDEQLDKWKEWAQRLELDVRTRKVTSPDEAKRWIKEFSEE |
| RO1_2 | KLFVLILSNDKKLIEEAKKMAEKANLELYVSSSEEDAKRILKELKDRNADSVLVLVSNDEQLDTAKEWAQRLELNVRTRKVTSPDEAKRWIKEFSEE |
| RO1_3 | TLVVVIISNDKKLIEEAKKMAEKANLQLYTDLDPDQAVKLAKKLNADKVLVLVSNDEDLDKAKEAAQRAELDVRIRKVTSPDEAKRWIKEFSEE |
| RO1_4 | KLVVIILSNDKKLIEEAQKMAEKANLELYVVDSEELKKLLKKIADENPNTKVLILVSNDEQLDLAKEIAQRLELDVRTRKVTSPDEAKRWIKEFSEE |
| RO1_5 | TLVVIIDSNDKKLIEEAKKMAEKANLELYYERDIEDLLRKLKDADRILILVSNDEQLDKAKEIAQRLEVPVRTRKVTSPDEAKRWIKEFSEE |
| RO1_6 | KLSVIILSNDKKLVEEAKKMAEKANLELYVVTDPDQAEKIIRKLIKEDPTIRILVLVSNDEALDWVKELAQKLEVDLRTRKVTSPDEAKRWIKEFSEE |
| RO1_7 | TLVVIILSNDKKLIEEAKKMAEKANLYLFEVTSDEDWKKAIKTAKEIAKKEQRPLRILVLVSNDEQLDKAKEIAQRQELDVRTRKVTSPDEAKRWIKEFSEE |
| RO1_8 | TLVVIIDSNDKKLIEEAKKMAEKANLILIESSPDPEKTLRDLNADRVLVLVSNDEQLDTWKEWAQRWELPIRTRKVTSPDEAKRWIKEFSEE |
| RO1_9 | TLLVIILSNDKKLVEEAKKMAEKANLILINSPLSPEQLERTVKSVNADRVLILVSNDEQLDQAKETAQRAELPIRTRKVTSPDEAKRWIKEFSEE |
| RO1_10 | TLVVIIISNDKKLIEEAKKMAEKANLILLVVDNPEEALERAYRLNADKILVLVSNDEQLDWAKEAAQRWELPVRVRKVTSPDEAKRWIKEFSEE |
| RO2_1 | RLVVLIVSNDKKLIEEARKMAEKANLELITVPGSPEEAIRLAQEIAEKAPGPVKVLVLITGSADPDEKTKAKKAAEEARKWNVRVRTVTSPDEAKRWIKEFSEE |
| RO2_2 | QLYVIISSNDKKLIEEARKMAEKANLNLLTADVDEAYELAKKLIDKAGSAKVLILITGSADPSQKKKIKELAEKARSLNVRIRIVTSPDEAKRWIKEFSEE |
| RO2_3 | TLVVIIVSKDKKLIEEARKMAEKANLLLVVYEPGEDEEAAKEASRRLKESLNNNQPAKVLVLISSSLSPSLAETAAKQLAPDAEVRIRTVTSPDEAKRWIKEFSEE |
| RO2_4 | TLVVFIVSKDKKLIEEARKMAEKANLKLYTAPVSPSIAEKVAKEAKKKNQPAKFLFLVDGTDPTAREIATKLAKYASTVANAEVRIREVTSPDLAKRWIKEFSEE |
| RO2_5 | GLLVIIVSKDKKLIEEARKMAEKANLLLITAPTDPRELETAIKLLQKSNTPIKILILSDGTDPTAEKIAKKLAKEAATKANAEYRIRKVTSPDQAKRWIKEFSEE |

| | |
|---|---|
| RO2_6 | GLVVIIVSKDKKLIEEARKMAEKANLYLFTLEPNADPSQLDTLRKWAQEILKRDGP SKLKVLVLSDGTDPTAQKLAKLIAKIVATAANAEVRIRSVTSPDQAKRWIKEFSEE |
| RO2_7 | TLIVVIVSKDKKLIEEARKMAEKANLLLFTGDLTNEQEKTAKEAADRDGSAKILILSD GTDPDARDKATKAAKKLATKLNAEFRIREVTSPDQAKRWIKEFSEE |
| RO2_8 | TLVVIIISDDKKLIEEARKMAEKANLILVTKSEIDDAIREIKKKAKDRPAKILILSDGTN PEAEKIAKKIAEKIAKILNAEVRIRKVTSPDQAKRWIKEFSEE |
| RO2_9 | NLIVFIWSNDKKLIEEARKMAEKANLYLFTLGDNAEKVLQEAVEKVAGDNVKILVLI EDTKDADKLAKKLKEIADKKNWDIRIRKVTSPDEAKRWIKEFSEE |
| RO2_10 | RLIVIILSNDKKLIEEARKMAEKANLELITVRSDEDIEKVLRKAGNAKVLLLIEDTKDA DKLAKKAKEAADKLNVDLRIRKVTSPDEAKRWIKEFSEE |
| RO2_11 | SLFVIIFSNDKKLIEEARKMAEKANLILITVEGSPSAVQEAIKIAVEIARKQNAESIKIL LLVENTKDAEKVKKLAKEAADKLNVDIRIRTVTSPDEAKRWIKEFSEE |
| RO2_12 | SLFVIIYSNDKKLIEEARKMAEKANLNLYTVSGDWREVKKLIEELIKRAKDKNPSEE VKVLLLVKDPRATEAAKKLEKNAPPNVRIRTVTSPDEAKRWIKEFSEE |
| RO2_13 | ELIVLILSNDKKLIEEARKMAEKANLELYTLEGDDEQIKKWIKKLAKTALSRNPSEA KILVLVEDTKDADKKIKIIKKAADEANIEIRIRKVTSPDEAKRWIKEFSEE |
| RO2_14 | QLFVIIVSNDKKLIEEARKMAEKANLELYTADLDTAVKIAKELLKKAEGPAKVLILVS GSASPDQKTKLDKIAKKLRSYNIRLREVTSPDEAKRWIKEFSEE |
| RO2_15 | KLVVLILSNDKKLIEEARKMAEKANLELLTLDGSPEQLKKILKTLLDKAGDRPLKILV LIEDTKDADKWAKAIKEAAKELNIDVRIRKVTSPDEAKRWIKEFSEE |
| RO2_16 | TLIVIIISNDKKLIEEARKMAEKANLNLYTWDDEDKAKKALKDATKYENVKLLFLIEN TKDAEKIEKKIKDTAKKLNLDVRVRLVTSPDEAKRWIKEFSEE |
| RO2_17 | TLIVVIWSNDKKLIEEARKMAEKANLLLLTVTSDEDLKKAAKIAQSAPGEVKVLLLV EDTKDADKIADKAKKIFKKANVDIRIRKVTSPDEAKRWIKEFSEE |
| RO2_18 | TLVVLIWSNDKKLIEEARKMAEKANLYLITVGDDKALEKAIRTAEKIAKDNNADSFK ILILIEDTKDADKISKKAKDIASKLNIEIRVRKVTSPDEAKRWIKEFSEE |
| RO2_19 | TLIVLIISNDKKLIEEARKMAEKANLLLYTLEPNQDPSIEKEIKTIQKRADPRDLKILVL IENTKDAEKIATEIKRKAEKNNLNVRIRLVTSPDEAKRWIKEFSEE |
| RO2_20 | GLLVLIWSNDKKLIEEARKMAEKANLYLLTLETDDKKIEDILKSLGPPVKILVLLEDT KDADKVKKEIEKKARKKNLPVRIRKVTSPDEAKRWIKEFSEE |
| RO2_21 | TLIVIIISNDKKLIEEARKMAEKANLVLITDEGSPSAEEKLKKTITDAKRKDPTDPVKI LVLIEDTKDADKIAEEIKRKADKANWDVRIRKVTSPDEAKRWIKEFSEE |
| RO2_22 | TLVVLIFSNDKKLIEEARKMAEKANLELYTRSELDPNIVTKLRDNAENAKLLVLIEDT KDADKLAEKIKKALDKNNIDVRIRKVTSPDEAKRWIKEFSEE |

| | |
|---|---|
| RO2_23 | SLVVFIWSNDKKLIEEARKMAEKANLELITVSSIDQAIKLAREIAKKQKRPAKFLILVSGSLDPSQKKKVDEIAKEARKDNIRVRTVTSPDEAKRWIKEFSEE |
| RO2_24 | KLLVVILSNDKKLIEEARKMAEKANLELITVTSLEEAKKAAEKALKEANGNAKVLVLITGSADPTQKKKATEWAKKAKDYNIRVRTVTSPDEAKRWIKEFSEE |
| RO2_25 | TLFVLILSNDKKLIEEARKMAEKANLILITVGDEEELKKAIKKADDIAKKQNSSEAKILILLEKPVSPEYEKKLQKYADAEVRVRTVTSPDEAKRWIKEFSEE |
| NT_1 | SREEIRKVVETFLRAANSQDKKKLEEAAKNILSPDVRLEVGNYTWTSIEQMLKFYQLSEIDRVEIRKVQVDGNHVRVEIEVERNGKKWTWEVEVEVRNGLIKRIRNQVDPEYKKDVQNIWNNT |
| NT_2 | SREEIRKVVEEFIRAQEDPDKLEKVASKALSPDVRVEIGNFTLEDKKQVIKWQKAFYKVLQEKAGKDASFRYEIRKVQVDGNHVRVEVEVETNGKKWTYEIELEVRNGKIKRIRLQVDPEYKEIVQLAWNRT |
| NT_3 | SREEIRKVVETWVRLFNSGDPRDRKKYEKAQKELLSPDVRTEIGNYTIEPGTLERFVQAYWKVLDELWPNVPIRVEIRKVQVDGNHVRIEVEVEINGKKYTFEIEVEVRNGKIKRIRIQRDPEMKELIQIAWNRT |
| NT_4 | SREEIRKVVETYVRISLSSSEETKKILRDLLSPDARLEFGNYTIESGDIWKFMQLFWKYYAGDAPLRLEIRKVQVDGNHVRIEVEVETKGKKWTYEIEVEVRNGKIKRVRTQVDPEYKKALQYAWNAT |
| NT_5 | SREEIRKIVELFVKAWDNPDAREKFEKNKDKVLSPDVRLEIGNFTLENKDKLESFYRVLIKLWQEKAGPNVRIEIRKVQVDGNHVRVEVEVETNGKKWTYEIEVEVRNGTIKRIRTQYDPEYKKDIQQAWNLS |
| NT_6 | SREEIRKIVETIVRANRDTSLFEKLAKELNLFSPDTRIEIGNYTFEGDAIKVIKAYIEANLQFAKKVSKDAPVRIEIRKVQVDGNHVRVEVEIELAGKKFTTEIEVEVRNGVVKRIRIQVDPEFKKLVQYAWNKT |
| NT_7 | SREEIRKVVEIFIRLQSLDPSQLEKALKDLNILSPDVRLEVGNITLNSADKLIRFLALITEILIRLWTGKPAPLRVEIRKVQVDGNHVRVEIEQEINGRKWTYEIEFEVRNGVIKRIRVQLDPSTKEAVQRAWNLT |
| NT_8 | SREEIRKVVETFVRAKQDPREFTKALSLLSPDVRMEIGNYTLTSIRDIKRFFEALVEIWKRKNLTDWRYEIRKVQVDGNHVRIEVETQTDGKKWTWEIEIEVRNGKIKRIREQYDPEYKKDVQLAWNLT |
| NT_9 | SREEIRKVVEEYIRLLYTDPDQFKKAARDKLLSPDVRIEIGNYTFDSRNLDRFLDAMQEWASRYDRVEIRKVQVDGNHVRVEIELESNGKKWTFEIEVEVRNGKIKRIRQQVDPEYKKVVQNLWNNT |
| NT_10 | SREEIRKVVETWIRLFYSSDPNDWETFQKAKKDLLSPDVRVEIGNYTLNSEQVDRWWEAWVKIIQKELEEKNEPLRTEIRKVQVDGNHVRVEIEQEKNGKKWTFEVEVEVRNGKIKRIRQQVDPEYKKEVQAAWNNT |

**Data S2. DNA sequences of designs selected for experimental testing**

| Design ID | DNA Sequence |
|---|---|
| RO1_1 | AAACTGGTGGTGGTGATTGATAGCAACGATAAAAAACTGATTGAAGAAGCGAAAAAAATGGCGGAAAAAGC<br>GAACCTGCTGCTGCTGTATGATGTGGATGAAGATCAGGTGCGTAAAGCGGCGGGCAACGCGCGTATTCTGGT<br>GCTGGTGAGCAACGATGAACAGCTGGATAAATGGAAAGAATGGGCGCAGCGTCTGGAACTGGATGTGCGTA<br>CCCGTAAAGTGACCAGCCCGGATGAAGCGAAACGTTGGATTAAAGAATTTAGCGAAGAATAATAAGGCAGCT<br>AAGGCAGCTAAGGC |
| RO1_2 | AAACTGTTTGTGCTGATTCTGAGCAACGATAAAAAACTGATTGAAGAAGCGAAAAAAATGGCGGAAAAAGCG<br>AACCTGGAACTGTATGTGAGCAGCAGCGAAGAAGATGCGAAACGTATTCTGAAAGAACTGAAAGATCGTAAC<br>GCGGATAGCGTGCTGGTGCTGGTGAGCAACGATGAACAGCTGGATACCGCGAAAGAATGGGCGCAGCGTCT<br>GGAACTGAACGTGCGTACCCGTAAAGTGACCAGCCCGGATGAAGCGAAACGTTGGATTAAAGAATTTAGCGA<br>AGAATAATAAGGC |
| RO1_3 | ACCCTGGTGGTGGTGATTATTAGCAACGATAAAAAACTGATTGAAGAAGCGAAAAAAATGGCGGAAAAAGCG<br>AACCTGCAGCTGTATACCGATCTGGATCCGGATCAGGCGGTGAAACTGGCGAAAAAACTGAACGCGGATAAA<br>GTGCTGGTGCTGGTGAGCAACGATGAAGATCTGGATAAAGCGAAAGAAGCGGCGCAGCGTGCGGAACTGGA<br>TGTGCGTATTCGTAAAGTGACCAGCCCGGATGAAGCGAAACGTTGGATTAAAGAATTTAGCGAAGAATAATA<br>AGGCAGCTAAGGC |
| RO1_4 | AAACTGGTGGTGATTATTCTGAGCAACGATAAAAAACTGATTGAAGAAGCGCAGAAAATGGCGGAAAAAGCG<br>AACCTGGAACTGTATGTGGTGGATAGCGAAGAACTGAAAAAACTGCTGAAAAAAATTGCGGATGAAAACCCG<br>AACACCAAAGTGCTGATTCTGGTGAGCAACGATGAACAGCTGGATCTGGCGAAAGAAATTGCGCAGCGTCTG<br>GAACTGGATGTGCGTACCCGTAAAGTGACCAGCCCGGATGAAGCGAAACGTTGGATTAAAGAATTTAGCGAA<br>GAATAATAAGGC |
| RO1_5 | ACCCTGGTGGTGATTATTGATAGCAACGATAAAAAACTGATTGAAGAAGCGAAAAAAATGGCGGAAAAAGCG<br>AACCTGGAACTGTATTATGAACGTGATATTGAAGATCTGCTGCGTAAACTGAAAGATGCGGATCGTATTCTGA<br>TTCTGGTGAGCAACGATGAACAGCTGGATAAAGCGAAAGAAATTGCGCAGCGTCTGGAAGTGCCGGTGCGTA<br>CCCGTAAAGTGACCAGCCCGGATGAAGCGAAACGTTGGATTAAAGAATTTAGCGAAGAATAATAAGGCAGCT<br>AAGGCAGCTAA |
| RO1_6 | AAACTGAGCGTGATTATTCTGAGCAACGATAAAAAACTGGTGGAAGAAGCGAAAAAAATGGCGGAAAAAGC<br>GAACCTGGAACTGTATGTGGTGACCGATCCGGATCAGGCGGAAAAAATTATTCGTAAACTGATTAAAGAAGAT<br>CCGACCATTCGTATTCTGGTGCTGGTGAGCAACGATGAAGCGCTGGATTGGGTGAAAGAACTGGCGCAGAAA<br>CTGGAAGTGGATCTGCGTACCCGTAAAGTGACCAGCCCGGATGAAGCGAAACGTTGGATTAAAGAATTTAGC<br>GAAGAATAATAA |
| RO1_7 | ACCCTGGTGGTGATTATTCTGAGCAACGATAAAAAACTGATTGAAGAAGCGAAAAAAATGGCGGAAAAAGCG<br>AACCTGTATCTGTTTGAAGTGACCAGCGATGAAGATTGGAAAAAAGCGATTAAAACCGCGAAAGAAATTGCG<br>AAAAAAGAACAGCGTCCGCTGCGTATTCTGGTGCTGGTGAGCAACGATGAACAGCTGGATAAAGCGAAAGAA<br>ATTGCGCAGCGTCAGGAACTGGATGTGCGTACCCGTAAAGTGACCAGCCCGGATGAAGCGAAACGTTGGATT<br>AAAGAATTTAGCGAAGAATAA |
| RO1_8 | ACCCTGGTGGTGATTATTGATAGCAACGATAAAAAACTGATTGAAGAAGCGAAAAAAATGGCGGAAAAAGCG<br>AACCTGATTCTGATTGAAAGCAGCCCGGATCCGGAAAAAACCCTGCGTGATCTGAACGCGGATCGTGTGCTGG<br>TGCTGGTGAGCAACGATGAACAGCTGGATACCTGGAAAGAATGGGCGCAGCGTTGGGAACTGCCGATTCGTA |

| | |
|---|---|
| | CCCGTAAAGTGACCAGCCCGGATGAAGCGAAACGTTGGATTAAAGAATTTAGCGAAGAATAATAAGGCAGCT<br>AAGGCAGCTAA |
| RO1_9 | ACCCTGCTGGTGATTATTCTGAGCAACGATAAAAAACTGGTGGAAGAAGCGAAAAAAATGGCGGAAAAAGCG<br>AACCTGATTCTGATTAACAGCCCGCTGAGCCCGGAACAGCTGGAACGTACCGTGAAAAGCGTGAACGCGGAT<br>CGTGTGCTGATTCTGGTGAGCAACGATGAACAGCTGGATCAGGCGAAAGAAACCGCGCAGCGTGCGGAACTG<br>CCGATTCGTACCCGTAAAGTGACCAGCCCGGATGAAGCGAAACGTTGGATTAAAGAATTTAGCGAAGAATAAT<br>AAGGCAGCTAA |
| RO1_10 | ACCCTGGTGGTGATTATTATTAGCAACGATAAAAAACTGATTGAAGAAGCGAAAAAAATGGCGGAAAAAGCG<br>AACCTGATTCTGCTGGTGGTGGATAACCCGGAAGAAGCGCTGGAACGTGCGTATCGTCTGAACGCGGATAAA<br>ATTCTGGTGCTGGTGAGCAACGATGAACAGCTGGATTGGGCGAAAGAAGCGGCGCAGCGTTGGGAACTGCC<br>GGTGCGTGTGCGTAAAGTGACCAGCCCGGATGAAGCGAAACGTTGGATTAAAGAATTTAGCGAAGAATAATA<br>AGGCAGCTAAGGC |
| RO2_1 | CGCCTTGTTGTATTGATCGTAAGTAATGACAAGAAGTTGATCGAAGAGGCCCGCAAGATGGCTGAGAAGGCT<br>AATTTGGAGTTGATCACGGTTCCAGGTAGTCCTGAGGAAGCCATCCGCTTGGCTCAAGAGATCGCCGAGAAG<br>GCTCCTGGGCCCGTTAAGGTATTGGTCTTAATCACAGGTTCAGCCGACCCCGACGAGAAGACGAAGGCCAAG<br>AAGGCAGCAGAGGAAGCTCGCAAGTGGAATGTCCGCGTTCGCACGGTTACATCTCCTGACGAGGCAAAGCGC<br>TGGATCAAGGAGTTCTCAGAGGAGTGA |
| RO2_2 | CAGCTGTATGTGATTATTTCCAGTAACGATAAGAAACTGATTGAAGAGGCGCGTAAAATGGCGGAAAAGGCA<br>AACCTGAACCTGCTTACAGCCGATGTGGATGAAGCGTATGAACTGGCGAAGAAATTGATTGATAAAGCAGGG<br>AGCGCGAAAGTACTTATTCTGATTACCGGCAGTGCGGATCCCTCTCAGAAGAAGAAAATTAAAGAACTCGCGG<br>AAAAGGCACGTAGCTTAAACGTACGTATTCGTATTGTTACCAGCCCGGATGAAGCGAAACGTTGGATTAAAGA<br>ATTTTCGGAAGAATAA |
| RO2_3 | ACACTCGTTGTCATCATCGTATCGAAGGACAAGAAGCTTATCGAAGAGGCCCGCAAGATGGCTGAGAAGGCT<br>AATTTGCTCTTGGTCGTTTACGAGCCAGGTGAGGACGAAGAGGCAGCCAAAGAGGCCAGTCGCCGCTTAAAG<br>GAGTCTCTCAATAATAATCAACCTGCAAAGGTCTTGGTTCTCATCTCAAGTTCCCTCTCACCTAGTTTAGCCGAG<br>ACGGCAGCCAAGCAATTGGCCCCAGACGCTGAGGTACGCATCCGCACGGTAACATCTCCAGACGAGGCAAAG<br>CGCTGGATCAAGGAGTTCTCTGAGGAGTGA |
| RO2_4 | ACTTTAGTCGTTTTCATCGTTAGTAAGGACAAGAAGCTCATCGAGGAAGCCCGCAAGATGGCCGAGAAGGCCA<br>ATCTTAAGCTCTACACAGCTCCAGTTTCCCCCTCTATCGCCGAGAAGGTCGCTAAAGAGGCAAAGAAGAAGAA<br>TCAACCAGCAAAGTTCCTCTTCCTCGTTGACGGTACGGACCCCACAGCACGCGAGATCGCCACAAAGTTAGCC<br>AAGTACGCATCAACTGTCGCCAATGCCGAGGTCCGCATCCGCGAGGTTACTTCTCCTGACCTTGCAAAGCGCT<br>GGATCAAGGAGTTCTCAGAGGAGTGA |
| RO2_5 | GGTCTCCTCGTAATCATCGTTAGCAAGGACAAGAAGTTAATTGAAGAAGCTCGCAAGATGGCCGAGAAGGCA<br>AATTTATTGTTAATCACAGCACCTACGGACCCACGCGAGTTGGAGACTGCAATCAAGTTACTTCAAAAGTCAAA<br>TACGCCAATCAAGATCTTAATCTTATCAGACGGAACGGACCCCACGGCTGAGAAGATCGCAAAGAAGTTGGCC<br>AAGGAAGCAGCAACGAAGGCCAATGCCGAGTACCGCATCCGCAAGGTTACTTCACCCGACCAAGCAAAGCGC<br>TGGATCAAGGAGTTCTCTGAGGAGTGA |
| RO2_6 | GGGCTCGTAGTCATCATCGTTTCTAAGGACAAGAAGTTAATCGAAGAGGCCCGCAAGATGGCCGAGAAGGCC<br>AATTTATACTTATTCACGTTAGAGCCAAATGCCGACCCAAGTCAATTGGACACGCTCCGCAAGTGGGCCCAAG<br>AGATCCTTAAGCGCGACGGGCCCTCAAAGCTCAAGGTATTGGTTCTCTCGGACGGTACGGACCCCACAGCACA<br>AAAGTTGGCCAAGCTTATCGCAAAGATCGTAGCAACGGCAGCCAATGCCGAGGTTCGCATCCGCTCCGTAACT<br>TCACCTGACCAAGCCAAGCGCTGGATCAAGGAGTTCTCCGAGGAGTGA |

| | |
|---|---|
| RO2_7 | ACTCTCATCGTCGTTATCGTATCTAAGGACAAGAAGTTAATCGAGGAAGCCCGCAAGATGGCCGAGAAGGCCA<br>ATTTACTCTTATTCACTGGAGACCTCACGAATGAGCAAGAGAAGACAGCAAAAGAGGCTGCCGACCGCGACG<br>GATCAGCTAAGATCCTTATCCTTAGTGACGGTACGGACCCTGACGCTCGCGACAAGGCTACGAAGGCCGCCAA<br>GAAGCTCGCAACAAAGCTTAATGCTGAGTTCCGCATCCGCGAGGTAACGTCACCTGACCAAGCCAAGCGCTGG<br>ATCAAGGAGTTCAGTGAGGAGTGA |
| RO2_8 | ACATTGGTTGTTATCATCATCTCAGACGACAAGAAGTTGATCGAGGAAGCACGCAAGATGGCTGAGAAGGCA<br>AATTTAATCTTGGTTACGAAGAGTGAGATCGACGACGCAATCCGCGAGATCAAGAAGAAGGCCAAGGACCGC<br>CCTGCCAAGATCTTAATCTTAAGTGACGGGACGAATCCAGAGGCAGAGAAGATCGCTAAGAAGATCGCAGAG<br>AAGATCGCCAAGATCCTCAATGCCGAGGTACGCATCCGCAAGGTAACGAGTCCTGACCAAGCAAAGCGCTGG<br>ATCAAGGAGTTCAGTGAGGAGTGA |
| RO2_9 | AATTTGATCGTTTTCATCTGGTCCAATGACAAGAAGCTTATCGAAGAGGCCCGCAAGATGGCAGAGAAGGCAA<br>ATTTGTACCTCTTCACGTTGGGAGACAATGCTGAGAAGGTTTTACAAGAGGCCGTTGAGAAGGTTGCCGGTGA<br>CAATGTAAAGATCTTGGTTTTGATCGAGGACACGAAGGACGCTGACAAGCTTGCAAAGAAGTTAAAGGAGAT<br>CGCAGACAAGAAGAATTGGGACATCCGCATCCGCAAGGTAACTTCGCCAGACGAGGCTAAGCGCTGGATCAA<br>GGAGTTCTCTGAGGAGTGA |
| RO2_10 | CGCTTGATCGTAATCATTCTGTCCAATGACAAGAAGCTTATCGAAGAGGCCCGCAAGATGGCCGAGAAGGCAA<br>ATCTTGAGCTTATCACAGTTCGCAGTGACGAGGACATCGAGAAGGTATTACGCAAGGCCGGGAATGCAAAGG<br>TCCTTTTGCTTATCGAGGACACGAAGGACGCCGACAAGTTGGCCAAGAAGGCCAAGGAAGCCGCCGACAAGT<br>TGAATGTAGACCTTCGCATCCGCAAGGTAACGTCTCCTGACGAGGCTAAGCGCTGGATCAAGGAGTTCAGTGA<br>GGAGTGATAA |
| RO2_11 | TCGTTGTTCGTCATCATCTTCTCAAATGACAAGAAGCTTATCGAGGAAGCCCGCAAGATGGCAGAGAAGGCCA<br>ATTTGATCTTGATCACAGTCGAGGGTTCTCCTTCTGCAGTCCAAGAGGCAATCAAGATCGCCGTTGAGATCGCA<br>CGCAAGCAAAATGCAGAGTCGATCAAGATCTTGCTTTTAGTCGAGAATACAAAGGACGCAGAGAAGGTTAAG<br>AAGCTCGCCAAAGAGGCCGCCGACAAGTTAAATGTTGACATCCGCATCCGCACTGTTACGTCTCCAGACGAGG<br>CTAAGCGCTGGATCAAGGAGTTCTCCGAGGAGTGA |
| RO2_12 | AGTCTTTTCGTTATCATCTACTCGAATGACAAGAAGCTCATCGAGGAAGCACGCAAGATGGCAGAGAAGGCTA<br>ATCTTAATTTATACACGGTTTCGGGAGACTGGCGCGAGGTTAAGAAGCTTATCGAGGAGTTGATCAAGCGCGC<br>CAAGGACAAGAATCCATCTGAGGAAGTTAAGGTCTTACTTTTAGTCAAGGACCCTCGCGCCACTGAGGCTGCC<br>AAGAAGTTAGAGAAGAATGCACCCCCAAATGTTCGCATCCGCACTGTTACGTCTCCCGACGAGGCCAAGCGCT<br>GGATCAAGGAGTTCTCAGAGGAGTGA |
| RO2_13 | GAGTTGATCGTTTTGATCCTTTCGAATGACAAGAAGCTCATCGAGGAAGCACGCAAGATGGCAGAGAAGGCC<br>AATCTCGAATTGTACACGTTGGAAGGTGACGACGAGCAAATCAAGAAGTGGATCAAGAAGCTCGCCAAGACT<br>GCCTTGTCTCGCAATCCCAGTGAGGCAAAGATCTTAGTCCTTGTTGAGGACACTAAGGACGCAGACAAGAAGA<br>TCAAGATCATCAAGAAGGCCGCCGACGAGGCAAATATCGAGATCCGCATCCGCAAGGTTACATCACCCGACG<br>AGGCAAAGCGCTGGATCAAGGAGTTCTCAGAGGAGTGA |
| RO2_14 | CAACTCTTCGTCATCATCGTATCAAATGACAAGAAGCTTATTGAAGAGGCACGCAAGATGGCAGAGAAGGCCA<br>ATCTCGAACTTTACACGGCAGACTTAGACACTGCAGTAAAGATCGCCAAGGAGTTGTTGAAGAAGGCCGAGG<br>GGCCAGCTAAGGTCCTCATCTTAGTCTCAGGGTCAGCATCGCCAGACCAAAAGACAAAGTTAGACAAGATCGC<br>CAAGAAGCTTCGCTCGTACAATATCCGCTTACGCGAGGTTACGTCTCCAGACGAGGCAAAGCGCTGGATCAAG<br>GAGTTCAGCGAGGAGTGA |
| RO2_15 | AAGCTGGTCGTCTTGATCTTGAGCAATGACAAGAAGTTGATTGAAGAGGCCCGCAAGATGGCCGAGAAGGCA<br>AATTTGGAGTTGCTTACGTTAGACGGTTCACCTGAGCAACTCAAGAAGATCCTTAAGACGCTTTTAGACAAGG |

| | |
|---|---|
| | CCGGAGACCGCCCATTGAAGATCTTGGTCTTGATCGAGGACACGAAGGACGCCGACAAGTGGGCCAAGGCAA<br>TCAAAGAGGCCGCCAAGGAGCTCAATATCGACGTCCGCATCCGCAAGGTCACATCCCCTGACGAGGCCAAGC<br>GCTGGATCAAGGAGTTCTCGGAAGAGTGA |
| RO2_16 | ACACTTATCGTCATCATCATCTCGAATGACAAGAAGCTCATCGAGGAAGCCCGCAAGATGGCCGAGAAGGCCA<br>ATTTAAATCTCTACACGTGGGACGACGAGGACAAGGCTAAGAAGGCATTAAAGGACGCAACTAAGTACGAGA<br>ATGTTAAGTTATTATTCCTCATCGAGAATACTAAGGACGCCGAGAAGATCGAGAAGAAGATCAAGGACACGG<br>CAAAGAAGCTCAATTTAGACGTCCGCGTTCGCTTGGTTACGTCGCCAGACGAGGCCAAGCGCTGGATCAAGG<br>AGTTCTCAGAGGAGTGA |
| RO2_17 | ACATTAATCGTCGTAATCTGGTCCAATGACAAGAAGTTAATCGAAGAGGCACGCAAGATGGCTGAGAAGGCA<br>AATCTCCTCTTATTGACAGTTACTTCCGACGAGGACTTGAAGAAGGCAGCAAAGATCGCACAAAGTGCCCCAG<br>GAGAGGTAAAGGTCCTTCTCCTTGTTGAGGACACAAAGGACGCTGACAAGATCGCCGACAAGGCCAAGAAGA<br>TCTTCAAGAAGGCCAATGTAGACATCCGCATCCGCAAGGTTACGTCGCCTGACGAGGCCAAGCGCTGGATCAA<br>GGAGTTCTCCGAGGAGTGA |
| RO2_18 | ACACTTGTAGTCTTAATCTGGTCGAATGACAAGAAGCTTATCGAAGAGGCACGCAAGATGGCCGAGAAGGCC<br>AATCTCTACCTTATCACGGTCGGTGACGACAAGGCCTTGGAGAAGGCTATCCGCACGGCAGAGAAGATCGCA<br>AAGGACAATAATGCCGACTCTTTCAAGATCTTAATCCTTATCGAGGACACAAAGGACGCCGACAAGATCAGTA<br>AGAAGGCAAAGGACATCGCCAGCAAGCTCAATATCGAGATCCGCGTTCGCAAGGTAACATCTCCTGACGAGG<br>CTAAGCGCTGGATCAAGGAGTTCTCTGAGGAGTGA |
| RO2_19 | ACTTTAATCGTTTTGATCATCTCGAATGACAAGAAGTTGATCGAGGAAGCCCGCAAGATGGCCGAGAAGGCTA<br>ATTTATTACTTTACACTCTTGAGCCTAATCAAGACCCATCCATCGAGAAGGAGATCAAGACGATCCAAAAGCGC<br>GCTGACCCACGCGACTTAAAGATCCTTGTTCTTATCGAGAATACTAAGGACGCAGAGAAGATCGCCACGGAGA<br>TCAAGCGCAAGGCAGAGAAGAATAATTTAAATGTACGCATCCGCCTTGTCACGTCGCCCGACGAGGCCAAGC<br>GCTGGATCAAGGAGTTCAGTGAGGAGTGA |
| RO2_20 | GGATTATTGGTCCTCATTTGGTCGAATGACAAGAAGCTCATCGAGGAAGCTCGCAAGATGGCTGAGAAGGCC<br>AATCTCTACCTCTTGACGCTCGAAACTGACGACAAGAAGATCGAGGACATCTTAAAGTCGCTCGGGCCGCCCG<br>TTAAGATCCTCGTTCTCTTAGAGGACACAAAGGACGCCGACAAGGTCAAGAAGGAGATCGAGAAGAAGGCCC<br>GCAAGAAGAATTTACCCGTACGCATCCGCAAGGTAACTTCGCCAGACGAGGCAAAGCGCTGGATCAAGGAGT<br>TCAGTGAGGAGTGA |
| RO2_21 | ACTTTGATTGTCATCATCATCAGTAACGACAAGAAGTTGATCGAGGAAGCTCGCAAGATGGCTGAGAAGGCCA<br>ATCTCGTCTTGATCACTGACGAGGGAAGTCCCAGTGCAGAGGAGAAGCTCAAGAAGACAATCACGGACGCCA<br>AGCGCAAGGACCCAACGGACCCAGTAAAGATCTTAGTATTAATCGAGGACACTAAGGACGCCGACAAGATCG<br>CCGAGGAGATCAAGCGCAAGGCTGACAAGGCAAATTGGGACGTCCGCATCCGCAAGGTAACATCCCCAGACG<br>AGGCAAAGCGCTGGATCAAGGAGTTCTCCGAGGAGTGA |
| RO2_22 | ACTTTGGTTGTTTTGATCTTCTCGAATGACAAGAAGTTGATCGAAGAGGCACGCAAGATGGCAGAGAAGGCA<br>AATCTCGAACTTTACACTCGCAGTGAGTTGGACCCAAATATCGTAACAAAGCTCCGCGACAATGCAGAGAATG<br>CCAAGCTTCTTGTATTGATCGAGGACACGAAGGACGCAGACAAGCTCGCCGAGAAGATCAAGAAGGCCCTCG<br>ACAAGAATAATATCGACGTACGCATCCGCAAGGTAACGTCGCCTGACGAGGCCAAGCGCTGGATCAAGGAGT<br>TCTCTGAGGAGTGA |
| RO2_23 | TCATTAGTTGTCTTCATCTGGTCCAATGACAAGAAGTTGATCGAGGAAGCTCGCAAGATGGCAGAGAAGGCTA<br>ATTTGGAGTTGATCACGGTCAGTTCAATCGACCAAGCAATCAAGTTGGCCCGCGAGATCGCAAAGAAGCAAAA<br>GCGCCCTGCCAAGTTCTTGATCTTGGTCTCAGGGAGTTTAGACCCCTCTCAAAAGAAGAAGGTTGACGAGATC |

| | |
|---|---|
| | GCTAAAGAGGCCCGCAAGGACAATATCCGCGTTCGCACAGTCACTTCGCCTGACGAGGCCAAGCGCTGGATC<br>AAGGAGTTCAGTGAGGAGTGA |
| RO2_24 | AAGTTGCTCGTTGTTATCCTTAGTAATGACAAGAAGTTGATCGAGGAAGCACGCAAGATGGCAGAGAAGGCA<br>AATTTAGAGTTAATCACGGTTACATCGCTTGAAGAGGCCAAGAAGGCCGCAGAGAAGGCCTTGAAAGAGGCA<br>AATGGTAATGCCAAGGTACTCGTTTTAATCACTGGGTCCGCCGACCCAACGCAAAAGAAGAAGGCAACTGAGT<br>GGGCAAAGAAGGCTAAGGACTACAATATCCGCGTACGCACAGTAACATCTCCAGACGAGGCCAAGCGCTGGA<br>TCAAGGAGTTCAGCGAGGAGTGA |
| RO2_25 | ACGCTTTTCGTACTCATCTTATCAAATGACAAGAAGCTTATCGAGGAAGCCCGCAAGATGGCAGAGAAGGCCA<br>ATTTGATCCTCATCACGGTCGGGGACGAAGAGGAGTTAAAGAAGGCCATCAAGAAGGCCGACGACATCGCTA<br>AGAAGCAAAATTCGTCAGAGGCCAAGATCCTCATCTTGCTTGAGAAGCCAGTCTCGCCTGAGTACGAGAAGAA<br>GTTACAAAAGTACGCAGACGCAGAGGTTCGCGTTCGCACAGTTACGTCACCAGACGAGGCCAAGCGCTGGAT<br>CAAGGAGTTCTCTGAGGAGTGA |
| NT_1 | TCGCGCGAGGAGATCCGTAAAGTAGTCGAAACCTTTCTGCGTGCTGCGAATAGCCAGGACAAAAAAAAACTC<br>GAAGAGGCGGCGAAAAATATTCTGTCACCTGATGTTCGTCTGGAAGTCGGCAACTATACCTGGACCAGCATTG<br>AACAAATGCTTAAGTTTTATCAACTGTCTGAGATTGATCGCGTCGAAATTCGCAAAGTGCAGGTCGATGGCAA<br>CCACGTGCGTGTGGAAATTGAAGTAGAACGTAATGGCAAAAAGTGGACCTGGGAAGTTGAGGTGGAAGTAC<br>GTAATGGTTTGATTAAACGCATTCGTAATCAGGTCGATCCGGAATATAAAAAAGATGTGCAAAATATCTGGAA<br>TAATACCTAA |
| NT_2 | AGCCGCGAAGAAATCCGTAAAGTAGTGGAAGAATTTATCCGTGCGCAAGAAGATCCCGACAAACTCGAGAAA<br>GTAGCGTCAAAGGCGCTGTCCCCGGACGTCCGCGTCGAAATTGGCAATTTCACATTGGAAGATAAGAAGCAG<br>GTCATCAAATGGCAAAAGGCCTTTTATAAAGTACTGCAGGAAAAAGCGGGGAAGGATGCCTCGTTTCGCTATG<br>AAATTCGCAAAGTACAAGTAGACGGTAATCATGTTCGGGTCGAAGTAGAAGTGGAAACGAACGGGAAAAAAT<br>GGACCTATGAAATCGAACTGGAGGTCCGTAATGGTAAAATCAAACGTATTCGCCTGCAAGTGGATCCGGAATA<br>TAAAGAAATCGTGCAACTGGCATGGAATCGTACATAA |
| NT_3 | TCTCGCGAGGAAATCCGCAAAGTCGTCGAAACGTGGGTTCGCTTATTCAACAGCGGCGACCCGCGCGACCGC<br>AAAAAATATGAAAAAGCTCAGAAGGAGCTGCTGTCCCCTGACGTTCGCACCGAAATTGGAAATTATACGATCG<br>AACCCGGAACCCTGGAGCGTTTTGTGCAGGCATACTGGAAAGTACTTGATGAGCTTTGGCCGAACGTCCCTAT<br>CCGCGTAGAGATCCGCAAAGTGCAAGTGGATGGCAACCATGTGCGTATTGAGGTCGAAGTAGAAATCAACGG<br>TAAAAAATATACGTTTGAAATTGAAGTCGAGGTGCGTAATGGAAAGATCAAGCGTATTCGTATTCAGCGCGAT<br>CCCGAAATGAAGGAGCTGATCCAGATCGCGTGGAACCGTACCTGA |
| NT_4 | AGTCGTGAAGAAATCCGCAAGGTTGTTGAAACGTACGTTCGCATCTCACTGAGCTCTAGCGAAGAAACCAAAA<br>AAATTCTGCGTGACTTGCTGTCGCCTGACGCACGCTTAGAATTTGGTAACTATACGATCGAATCCGGTGATATT<br>TGGAAATTCATGCAGTTGTTTTGGAAATACTACGCCGGCGACGCACCGCTCCGCCTGGAAATCCGTAAAGTGC<br>AGGTTGATGGTAATCATGTTCGTATCGAAGTCGAAGTCGAGACCAAGGGAAAAAAATGGACCTATGAAATTG<br>AAGTGGAAGTCCGCAACGGTAAAATTAAACGTGTTCGCACCCAAGTTGATCCGGAATATAAGAAAGCATTGCA<br>ATATGCCTGGAACGCAACCTAA |
| NT_5 | TCTCGTGAAGAGATTCGCAAAATCGTGGAGCTCTTTGTAAAAGCATGGGATAATCCTGACGCTCGCGAAAAGT<br>TTGAAAAGAATAAAGACAAGGTATTATCTCCCGATGTCCGTCTGGAAATTGGGAATTTTACCCTGGAAAATAA<br>AGATAAACTTGAATCTTTCTACCGTGTCCTGATCAAACTTTGGCAGGAGAAAGCTGGTCCCAACGTCCGTATCG<br>AGATCCGCAAGGTGCAAGTCGATGGTAACCATGTGCGTGTCGAAGTTGAAGTGGAAACGAATGGCAAAAAAT<br>GGACCTACGAAATTGAAGTAGAAGTTCGTAACGGTACGATTAAACGCATCCGCACTCAGTATGACCCTGAATA<br>TAAAAAGGACATCCAGCAGGCTTGGAACCTCTCATAA |

| | |
|---|---|
| NT_6 | AGCCGGGAAGAAATCCGCAAAATTGTGGAAACGATCGTACGTGCGAATCGCGACACGAGTCTCTTTGAGAAG<br>TTAGCCAAAGAACTGAATCTCTTTAGTCCGGACACCCGTATTGAAATTGGCAATTATACTTTTGAAGGGGATGC<br>GATCAAAGTTATCAAAGCGTACATTGAAGCTAATCTGCAGTTCGCGAAAAAGGTTTCGAAAGACGCGCCGGTG<br>CGTATTGAAATCCGTAAAGTGCAAGTGGATGGTAACCACGTTCGTGTTGAGGTTGAAATCGAACTGGCTGGAA<br>AGAAATTTACAACAGAGATCGAGGTTGAAGTGCGCAACGGGGTGGTTAAGCGCATTCGTATTCAAGTCGACC<br>CCGAGTTCAAAAAACTGGTTCAGTACGCTTGGAATAAAACGTAA |
| NT_7 | AGCCGCGAGGAAATTCGGAAGGTGGTAGAAATTTTCATTCGTCTTCAAAGCCTGGACCCGAGCCAGTTAGAAA<br>AAGCTTTAAAAGATTTGAACATTCTTTCGCCGGATGTGCGCCTGGAAGTCGGGAATATCACGCTGAACTCCGC<br>GGATAAACTCATTCGTTTCTTAGCGCTCATTACGGAGATCCTGATTCGCCTGTGGACGGGCAAACCCGCGCCTC<br>TGCGTGTAGAAATCCGTAAAGTGCAGGTAGATGGTAATCACGTGCGCGTGGAGATTGAACAAGAAATTAACG<br>GGCGTAAGTGGACGTACGAAATTGAATTTGAAGTACGCAATGGCGTTATTAAACGTATTCGTGTGCAGCTGGA<br>CCCATCCACCAAAGAGGCTGTCCAGCGTGCATGGAACCTTACCTAA |
| NT_8 | AGTCGCGAGGAAATTCGTAAAGTTGTGGAGACCTTCGTCCGTGCAAAGCAGGATCCGCGTGAATTCACCAAA<br>GCGTTGTCCCTGCTCAGCCCTGATGTTCGGATGGAAATCGGCAACTACACTCTGACCTCCATCCGGGACATTAA<br>ACGCTTTTTCGAGGCCTTAGTAGAAATTTGGAAACGTAAAAACTTGACTGACTGGCGTTACGAAATTCGCAAG<br>GTGCAGGTCGATGGCAATCACGTCCGCATTGAGGTGGAAACACAGACCGACGGCAAAAAATGGACTTGGGAA<br>ATTGAAATCGAAGTACGCAATGGGAAGATCAAACGTATCCGCGAGCAATATGACCCTGAATACAAAAAGGAT<br>GTACAGCTGGCATGGAACCTGACTTGA |
| NT_9 | AGCCGCGAAGAGATCCGCAAAGTTGTCGAGGAGTATATCCGTCTGCTGTATACAGACCCTGATCAGTTCAAGA<br>AAGCGGCCCGCGATAAATTGCTGAGTCCGGATGTGCGTATTGAAATCGGTAATTATACGTTTGATTCCCGCAA<br>CCTCGATCGCTTTCTGGACGCAATGCAGGAATGGGCGAGCCGTTACGATCGTGTGGAAATTCGCAAGGTTCAG<br>GTTGACGGGAACCATGTGCGTGTTGAGATTGAGTTGGAATCGAACGGTAAAAAATGGACGTTTGAAATCGAG<br>GTTGAGGTTCGCAATGGCAAAATTAAGCGCATCCGTCAACAGGTGGATCCGGAATATAAAAAAGTTGTGCAG<br>AATCTGTGGAATAACACGTGA |
| NT_10 | TCCCGGGAAGAGATTCGGAAGGTCGTGGAGACTTGGATCCGCCTTTTTTATAGCTCGGATCCGAACGACTGGG<br>AAACGTTCCAGAAAGCGAAGAAAGATCTGCTCTCACCAGATGTGCGCGTAGAAATCGGAAATTACACGCTTAA<br>TAGTGAACAAGTCGATCGTTGGTGGGAAGCCTGGGTGAAAATCATCCAGAAAGAATTGGAAGAGAAAAACG<br>AACCGCTCCGCACGGAAATTCGCAAGGTTCAAGTGGACGGCAATCACGTACGCGTGGAAATCGAGCAGGAAA<br>AAAACGGCAAGAAGTGGACCTTTGAAGTTGAAGTGGAAGTTCGTAATGGAAAAATCAAGCGTATTCGTCAAC<br>AGGTAGACCCAGAGTACAAAAAGGAGGTCCAGGCGGCTTGGAACAACACCTAA |

**References**

1. Dawson, N.L., Lewis, T.E., Das, S., Lees, J.G., Lee, D., Ashford, P., Orengo, C.A. & Sillitoe, I. CATH: an expanded resource to predict protein function through structure and sequence. *Nucleic Acids Res* **45**, D289-D295 (2017).
2. Kabsch, W. & Sander, C. Dictionary of protein secondary structure: pattern recognition of hydrogen-bonded and geometrical features. *Biopolymers* **22**, 2577-637 (1983).
3. Leaver-Fay, A., Tyka, M., Lewis, S.M., Lange, O.F., Thompson, J., Jacak, R., Kaufman, K., Renfrew, P.D., Smith, C.A., Sheffler, W., Davis, I.W., Cooper, S., Treuille, A., Mandell, D.J., Richter, F., Ban, Y.E., Fleishman, S.J., Corn, J.E., Kim, D.E., Lyskov, S., Berrondo, M., Mentzer, S., Popovic, Z., Havranek, J.J., Karanicolas, J., Das, R., Meiler,

J., Kortemme, T., Gray, J.J., Kuhlman, B., Baker, D. & Bradley, P. ROSETTA3: an object-oriented software suite for the simulation and design of macromolecules. *Methods Enzymol* **487**, 545-74 (2011).
4. Gront, D., Kulp, D.W., Vernon, R.M., Strauss, C.E. & Baker, D. Generalized fragment picking in Rosetta: design, protocols and applications. *PLoS One* **6**, e23294 (2011).
5. Chaudhury, S., Lyskov, S. & Gray, J.J. PyRosetta: a script-based interface for implementing molecular modeling algorithms using Rosetta. *Bioinformatics* **26**, 689-91 (2010).
6. Koga, N., Tatsumi-Koga, R., Liu, G., Xiao, R., Acton, T.B., Montelione, G.T. & Baker, D. Principles for designing ideal protein structures. *Nature* **491**, 222-7 (2012).
7. Shapovalov, M.V. & Dunbrack, R.L., Jr. A smoothed backbone-dependent rotamer library for proteins derived from adaptive kernel density estimates and regressions. *Structure* **19**, 844-58 (2011).
8. Sheffler, W. & Baker, D. RosettaHoles2: a volumetric packing measure for protein structure refinement and validation. *Protein Sci* **19**, 1991-5 (2010).
9. Marcos, E., Basanta, B., Chidyausiku, T.M., Tang, Y., Oberdorfer, G., Liu, G., Swapna, G.V., Guan, R., Silva, D.A., Dou, J., Pereira, J.H., Xiao, R., Sankaran, B., Zwart, P.H., Montelione, G.T. & Baker, D. Principles for designing proteins with cavities formed by curved beta sheets. *Science* **355**, 201-206 (2017).
10. Lawrence, M.C. & Colman, P.M. Shape complementarity at protein/protein interfaces. *J Mol Biol* **234**, 946-50 (1993).
11. Maguire, J.B., Boyken, S.E., Baker, D. & Kuhlman, B. Rapid Sampling of Hydrogen Bond Networks for Computational Protein Design. *J Chem Theory Comput* **14**, 2751-2760 (2018).
12. Smith, C.A. & Kortemme, T. Backrub-like backbone simulation recapitulates natural protein conformational variability and improves mutant side-chain prediction. *J Mol Biol* **380**, 742-56 (2008).
13. Zhang, C., Freddolino, P.L. & Zhang, Y. COFACTOR: improved protein function prediction by combining structure, sequence and protein-protein interaction information. *Nucleic Acids Res* **45**, W291-W299 (2017).
14. Zhou, J. & Grigoryan, G. Rapid search for tertiary fragments reveals protein sequence-structure relationships. *Protein Sci* **24**, 508-24 (2015).
15. Studier, F.W. Protein production by auto-induction in high density shaking cultures. *Protein Expr Purif* **41**, 207-34 (2005).
16. Delaglio, F., Grzesiek, S., Vuister, G.W., Zhu, G., Pfeifer, J. & Bax, A. NMRPipe: a multidimensional spectral processing system based on UNIX pipes. *J Biomol NMR* **6**, 277-93 (1995).
17. Wishart, D.S., Bigam, C.G., Yao, J., Abildgaard, F., Dyson, H.J., Oldfield, E., Markley, J.L. & Sykes, B.D. 1H, 13C and 15N chemical shift referencing in biomolecular NMR. *J Biomol NMR* **6**, 135-40 (1995).
18. Vranken, W.F., Boucher, W., Stevens, T.J., Fogh, R.H., Pajon, A., Llinas, M., Ulrich, E.L., Markley, J.L., Ionides, J. & Laue, E.D. The CCPN data model for NMR spectroscopy: development of a software pipeline. *Proteins* **59**, 687-96 (2005).

19. Cheung, M.S., Maguire, M.L., Stevens, T.J. & Broadhurst, R.W. DANGLE: A Bayesian inferential method for predicting protein backbone dihedral angles and secondary structure. *J Magn Reson* **202**, 223-33 (2010).
20. Rieping, W., Habeck, M., Bardiaux, B., Bernard, A., Malliavin, T.E. & Nilges, M. ARIA2: automated NOE assignment and data integration in NMR structure calculation. *Bioinformatics* **23**, 381-2 (2007).
21. Brunger, A.T., Adams, P.D., Clore, G.M., DeLano, W.L., Gros, P., Grosse-Kunstleve, R.W., Jiang, J.S., Kuszewski, J., Nilges, M., Pannu, N.S., Read, R.J., Rice, L.M., Simonson, T. & Warren, G.L. Crystallography & NMR system: A new software suite for macromolecular structure determination. *Acta Crystallogr D Biol Crystallogr* **54**, 905-21 (1998).
22. Berman, H., Henrick, K. & Nakamura, H. Announcing the worldwide Protein Data Bank. *Nat Struct Biol* **10**, 980 (2003).
23. Bhattacharya, A., Tejero, R. & Montelione, G.T. Evaluating protein structures determined by structural genomics consortia. *Proteins* **66**, 778-95 (2007).
24. Skinner, S.P., Goult, B.T., Fogh, R.H., Boucher, W., Stevens, T.J., Laue, E.D. & Vuister, G.W. Structure calculation, refinement and validation using CcpNmr Analysis. *Acta Crystallogr D Biol Crystallogr* **71**, 154-61 (2015).
25. Winter, G. xia2: an expert system for macromolecular crystallography data reduction. *Journal of Applied Crystallography* **43**, 186-190 (2010).
26. Kabsch, W. Xds. *Acta Crystallogr D Biol Crystallogr* **66**, 125-32 (2010).
27. Evans, P. Scaling and assessment of data quality. *Acta Crystallogr D Biol Crystallogr* **62**, 72-82 (2006).
28. McCoy, A.J., Grosse-Kunstleve, R.W., Adams, P.D., Winn, M.D., Storoni, L.C. & Read, R.J. Phaser crystallographic software. *J Appl Crystallogr* **40**, 658-674 (2007).
29. Adams, P.D., Afonine, P.V., Bunkoczi, G., Chen, V.B., Davis, I.W., Echols, N., Headd, J.J., Hung, L.W., Kapral, G.J., Grosse-Kunstleve, R.W., McCoy, A.J., Moriarty, N.W., Oeffner, R., Read, R.J., Richardson, D.C., Richardson, J.S., Terwilliger, T.C. & Zwart, P.H. PHENIX: a comprehensive Python-based system for macromolecular structure solution. *Acta Crystallogr D Biol Crystallogr* **66**, 213-21 (2010).
30. Kantardjieff, K.A. & Rupp, B. Matthews coefficient probabilities: Improved estimates for unit cell contents of proteins, DNA, and protein-nucleic acid complex crystals. *Protein Sci* **12**, 1865-71 (2003).
31. Weichenberger, C.X. & Rupp, B. Ten years of probabilistic estimates of biocrystal solvent content: new insights via nonparametric kernel density estimate. *Acta Crystallogr D Biol Crystallogr* **70**, 1579-88 (2014).
32. Afonine, P.V., Grosse-Kunstleve, R.W., Echols, N., Headd, J.J., Moriarty, N.W., Mustyakimov, M., Terwilliger, T.C., Urzhumtsev, A., Zwart, P.H. & Adams, P.D. Towards automated crystallographic structure refinement with phenix.refine. *Acta Crystallogr D Biol Crystallogr* **68**, 352-67 (2012).
33. Moriarty, N.W., Grosse-Kunstleve, R.W. & Adams, P.D. electronic Ligand Builder and Optimization Workbench (eLBOW): a tool for ligand coordinate and restraint generation. *Acta Crystallogr D Biol Crystallogr* **65**, 1074-80 (2009).
34. Bernstein, F.C., Koetzle, T.F., Williams, G.J., Meyer, E.F., Jr., Brice, M.D., Rodgers, J.R., Kennard, O., Shimanouchi, T. & Tasumi, M. The Protein Data Bank. A

computer-based archival file for macromolecular structures. *Eur J Biochem* **80**, 319-24 (1977).
